# Coevolving Mutations in Chronic SARS-CoV-2 Infections

**DOI:** 10.64898/2026.06.11.731720

**Authors:** Ryan Hisner, Ravindra K Gupta, Darren P Martin

## Abstract

The SARS-CoV-2 pandemic has been marked by two outstanding features that have had major impacts on global public health: the repeated emergence and growth of highly divergent saltation variants with no known close relatives and the development of adverse health effects extending beyond the period of acute infection, called long Covid, in a proportion of the population. Chronic infections in immunocompromised hosts are the most likely explanation for the emergence of saltation variants, and some evidence indicates that viral persistence contributes to long Covid. Knowledge of intrahost evolution during prolonged SARS-CoV-2 infection is therefore vital for understanding the global evolution of SARS-CoV-2 and for deciphering the nature of long Covid and promising avenues for treatment. We assembled a collection of over 3000 independent, full-length SARS-CoV-2 sequences deriving from posited or confirmed chronic infections. We describe 14 distinct mutation patterns (MPs) that repeatedly appear in these sequences—each involving mutations in multiple genomic regions—including four CD8 T cell-escape MPs and two MPs that represent adaptation to tissue compartments outside the upper-respiratory tract. The existence of these MPs promises new insights into the life cycle and evolution of SARS-CoV-2 and the nature of persistent SARS-CoV-2 infection.

## INTRODUCTION

When a virus infects and replicates in its host, mutations sporadically arise within its progeny. Although the rates at which mutations occur are especially high in single-stranded RNA viruses, within a host at any given moment the number of genome copies carrying mutations at any given nucleotide position, termed an intrahost single nucleotide variant (iSNV), is normally very low. Accordingly, most normal-duration SARS-CoV-2 infections are expected to harbor only two or fewer iSNVs that exceed 2% prevalence^1–3^. Furthermore, even though low prevalence iSNVs will likely occur at many different genome sites in many different genomes, almost all of this fledgeling within-host diversity is erased upon infection of a new host. The reason for this is that in most cases transmission events involve the initiation of a new infection by a single genome. This extremely tight transmission bottleneck means that, in most instances, only a minute fraction of all iSNVs that arise in any given infection outlast that infection by being transmitted to a new host^2, 4–6^.

Such limited intrahost diversity and tight transmission bottlenecks—which may be a universal feature of aerosol-transmitted pathogens^7^—hinder positive selection and introduce a large random element into viral evolution. Furthermore, due to the proofreading function of the coronavirus NSP14, the average mutation rate per site in coronaviruses is much lower than in other RNA viruses—23.9-fold lower in SARS-CoV-2 than in Influenza A Virus (IAV) according to one analysis^8^. The relatively low mutation rate and tight transmission bottleneck might lead one to expect adaptive evolution to proceed at a lethargic pace in SARS-CoV-2, similar to the endemic human coronaviruses (HCoVs) OC43 or 229E, whose receptor binding domains (RBDs) acquire substitutions at roughly one-fourth the rate of H3N2 IAV^9^. Instead, the SARS-CoV-2 RBD substitution rate has been more than twice as fast as H3N2^9^, begging the question of how such rapid antigenic adaptation is achieved given the constraints imposed by transmission bottlenecks and low replication error rates.

The evolution of SARS-CoV-2 has been marked by sudden “jumps” in which a new, highly divergent variant derived from a lineage that ceased circulating many months or years ago, appears and replaces circulating variants. These saltation variants are marked by rapid evolution, a high percentage of non-synonymous nucleotide mutations, and a high proportion of mutations in spike, particularly in the receptor binding domain (RBD)^46–48^. Mutations and mutational patterns seen in saltation variants closely resemble those seen in many case studies of chronic infections in people who are immunocompromised^10–38^. Many of the most common spike mutations observed in SARS-CoV-2 genomes sampled from people with chronic infections early in the pandemic (e.g. S50L, Δ69-70, R346T, K356T, L368I, K444X, V445X, G446X, N460K, E484X, F486X, N501Y) later became fixed in dominant circulating variants. Furthermore, a large proportion of the most highly divergent saltation variants have originated in southern Africa, which has the highest number of people living with HIV (PLWH) of any region in the world: a number of case studies have documented extensive intrahost evolution in PLWH^23, 31, 38, 43, 45, 48–49^.

The ability of SARS-CoV-2 to infect a wide variety of tissues, in part due to the widespread presence of its primary receptor, ACE2, likely facilitates intrahost adaptive evolution during chronic infections. SARS-CoV-2 has been shown to infect the liver^257–259^, kidney^260–264^, gastrointestinal tract^265–267^, fetal brain^268^, fat cells^269–270^, heart^260, 271–272^, and many other organs and cell types. The existence of numerous viral compartments across different tissues within a host creates a large viral population, extensive intrahost diversity, and a wealth of genetic resources the virus can draw from. When viral populations from different tissue compartments encounter one another, the continual recombination inherent in coronavirus replication can enable the traversal of fitness valleys and the creation of novel mutational combinations.

While it may never be possible to prove with certainty that chronic infections have given rise to the major SARS-CoV-2 anachronistic saltation variants, the wealth of evidence pointing in this direction, combined with the paucity of evidence for any competing hypotheses, has resulted in widespread acceptance of this hypothesis. In addition to the effect chronic infections have had on the evolution of SARS-CoV-2 and the course of the pandemic, such viral persistence is likely responsible for some proportion of the malign long-term health effects of SARS-CoV-2 infection^50–55^, which include both persistent life-altering symptoms such as fatigue and brain fog^56–58^ and long-term increased risk of adverse psychological, neurological, gastrointestinal, and cardiovascular health events^59–64^. Understanding the nature of such chronic infections, therefore, should be an urgent public health priority.

Here we attempt to illuminate the selective processes at play during long-term infections by examining a manually curated set of 3332 SARS-CoV-2 sequences displaying evidence of having undergone substantial evolution within the context of persistent infections. We identify 14 distinct patterns of repeatedly co-occurring sets of mutations that likely arose within persistent infection contexts, and which are therefore likely to have enhanced viral fitness within such contexts. Two of these mutation patterns (MPs) are likely adaptive in tissues outside the upper-respiratory tract and four others likely represent sets of human leukocyte (HLA) specific CD8 T cell escape mutations. These patterns provide insights into the nature of persistent infections and the adaptive processes that drive intrahost SARS-CoV-2 evolution.

## METHODS

### Sequences Datasets

We used an iterative approach to assemble a dataset of sequences from the GISAID database^65^ that had most likely arisen in a chronic infection context. An initial posited chronic-infection (PCI) sequence dataset was assembled manually over the course of over two years between April 2022 and January 2025. Candidate chronic-infection sequences were identified through (i) manual examination of individual sequences and sequence metadata on GISAID, (2) manually scanning 5000-sequence subsamples of the most up-to-date global SARS-CoV-2 phylogenetic tree for sequences separated from the rest of the phylogeny by unusually long branches, and (iii) manual examination of sequences on Nextclade^66^. After collecting several hundred such sequences, manual searches on GISAID and CovSpectrum^65, 67^ for mutations found to be common in PCI sequences (present in >2% of the sequences) but rare in circulating lineages (present in <0.1% of sequences) were performed. ORF1a:K1795Q, for example, is present as a private mutation in 9.2% of the PCI sequences but <0.01% of all good-quality circulating sequences (excluding the Gamma VOC lineage and lineage BS.1) as of 2025-01-12. Approximately 1200 additional sequences carrying mutations that differentiated the PCI sequences from circulating sequences were added to a preliminary expanded posited chronic-infection (EPCI) dataset together with the original set of PCI sequences.

Nextclade CLI was used to create ndjson files of the EPCI sequence collection. Custom written Julia code (https://github.com/ryhisner/MP_EPCI_code) was then used to identify and count the private mutations for each sequence (with private mutations being defined by Nextclade as “the mutations between the query sequence and the sequence corresponding to the nearest neighbor (parent) on the tree.”). Closely related sequences very likely collected from a single individual were identified by a combination of geographical location (same country or same state for US sequences), shared Pango lineage designations, and six or more shared private mutations. Further, sequences carrying >6 reversion mutations were excluded because these were likely either low-quality sequences or recombinants. The phylogenetic placement of the EPCI sequences in the global SARS-CoV-2 tree was examined using USHER^68^ and clusters of EPCI sequences within this tree were manually examined using Nextclade web^66^ to determine whether sequences in these clusters all had a recent common origin or had independently evolved but were spuriously clustered within the global SARS-CoV-2 tree due to presumed long branch attraction^69^.

Any sequences with fewer than eight private amino acid mutations + deletion ranges (where deletion range is defined as any continuous deletion of ≥1 consecutive amino acid residues) were automatically excluded from the EPCI dataset. In addition, each EPCI sequence was required to meet at least one of the following two requirements:

1. To possess at least 10 AA substitutions + deletion ranges
2. To possess at least 8 AA substitutions + deletion ranges and be among the last 5% of sequences collected in its designated Pango lineage, as assigned by Nextclade CLI, with exceptions being made for lineages A, B, and B.1 as the collection date statistics for each of these lineages are frequently marred by incorrect collection date labels and/or low-coverage, low-quality sequences that nevertheless pass stringent QC filters due to reversion to the reference genome in areas of low/no sequencing coverage.

For groups of related sequences (typically from the same individual), the sequence with the most private amino acid mutations was chosen as the single representative for each group. This yielded the final EPCI dataset comprising 3332 sequences each from a different potential chronic infection. The GISAID accession numbers for the final EPCI dataset can be found at https://github.com/ryhisner/EPCI_dataset.

### Mutational Pattern Detection

Initially, six of what ultimately turned out to be 14 distinct mutation patterns (MPs), were detected manually in the course of constructing the EPCI database. The remaining eight MPs were detected using custom Julia code that automatically detects statistically significant co-occurrences of mutations in one small (adjustable) region of the genome with mutations at other individual amino acid (AA) sites in the remainder of the genome https://github.com/ryhisner/MP_EPCI_code).

Specifically this involved the use of an iterative, MP-detecting (MPD) function consisting of two parts (hereafter referred to as F1 and F2) to yield a list of associated genomic regions and the specific mutations contained in each region. F1 detects genomic regions where mutations co-occur in the EPCI dataset, based on a single “seed mutation” that may either be part of a subjectively observed MP in the EPCI dataset or simply a mutation that is found to be unusually common in the EPCI dataset relative to circulating sequences. The genomic region associated with this mutation is initially defined as any AA position within six positions upstream or downstream of its AA position. As new mutations are added to a region, the associated region expands to encompass any AA site within six nucleotide sites of any of the region’s mutations. For example, if the seed mutation is E:T9I, an initial genomic region bound would be set as E:5-13. If E:T11A is then found to be strongly associated with E:T9I, the genomic region bound would expand to E:5-15. The MPD function can be set to use either specific amino acid substitutions (MPD1) or the positions at which amino acid substitutions take place (MPD2).

For both MDP1 and MDP2, a single “seed” mutation must be selected. Then, all sequences with that mutation are collected and their private AA substitutions (as determined by Nextclade CLI) tallied in order to determine if mutations falling at other genome sites/regions show signs of co-occurring with this region. All mutations in the qualifying sequences are then tested to determine if their co-occurrences with the seed mutation/region are statistically unlikely to have occurred due to chance alone. Specifically, a Fisher’s exact test -log10 p-value is calculated for each mutation based on (1) the number of times it co-occurs with the seed mutation in the qualifying sequences, (2) the number of times it occurs without the seed mutation in qualifying sequences, (3) the number of times it occurs in the entire EPCI dataset, and (4) the overall number of qualifying sequences.

Different p-value thresholds are used for mutations occurring within one of the starting regions and mutations that occur outside of these regions, with more stringent requirements for the latter. If a mutation occurs within one of the starting regions, a Fisher’s exact test p-value < 0.01 is the default requirement. For mutations occurring outside the starting regions, a Fisher’s exact test p-value < 0.00001 is the default requirement, and if this requirement is met, a new genomic region is added in subsequent rounds. Finally, a mutation must occur at least three times in the EPCI dataset for it to qualify as being a potential member of any MP.

A private mutation adjustment factor (PMA) is used to compensate for there being varying numbers of private mutations in each sequence. PMA was calculated as follows: The average number of private mutations in all EPCI sequences was divided by the number of private mutations in a given sequence to give the initial private mutation adjustment factor (IPMA). The minimum allowed value for IPMA is 0.6 and the maximum 1.5. A IPMA of 0.6 would mean that the sequence has more than 1.67 times as many private amino acid substitutions as the average EPCI sequence, while an IPMA of 1.5 would mean it has fewer than two-thirds as many private amino acid substitutions as the average EPCI sequence. The IPMA for a sequence is then added to one, and that sum divided by two to obtain the final PMA. Each mutation in a sequence is multiplied by the PMA when counting mutations, so that sequences with larger numbers of private mutations need to have a greater number of MP mutations to qualify than do sequences with lower numbers of private mutations. For example, if the required number of MP mutations for inclusion is two, a sequence that has twice as many private mutations as the average EPCI sequence would have a PMA of 1.25 and would therefore need to have three (≥2.50) MP mutations to qualify.

The minimum number of mutations needed for a sequence to qualify depends on the number of regions present at the beginning of each round, with a greater number of mutations required for each additional starting region. The number of regions in the MP is multiplied by a private mutation requirement factor (PMR) to obtain the required number of mutations a sequence must have in order to qualify. The default PMR value is 0.4 for the first.

After each round of F1, if any mutations have been removed or added, the same process is repeated but with the new set of mutations (and any new regions) included. If no mutations have been added or removed, the function ends, and the final set of mutations and associated regions are fed into F2.

F1 is biased because the sequences were selected based on the possession of the very mutations to be tested, and F2 is designed to compensate for this bias. F2 takes the regions delivered from F1 and tests each of them separately for statistical correlation with the other genomic regions. During each round of F2, there are N iterations, where N is the number of qualifying genomic regions from F1. In each iteration, one region is removed from the full mutation set and the remaining regions are used to select a set of sequences in the same manner as in F1. Only mutations in the removed region (which played no role in selecting the qualifying sequences) are tested. First, each mutation in the removed region is tested to see if it appears more often than might be expected by chance in the qualifying sequences. If any of the mutations in the region have a Fisher’s exact test p-value < 10^-5^, the region qualifies. Each mutation within the region’s range is then tested, and if it has a Fisher’s exact test p-value < 10^-2^, it is included in the mutation region. If no single mutation has a Fisher’s exact test p-value < 10^-5^, the region is disqualified.

No new regions are added in F2. In this manner the statistical association of the mutations in each region with the remaining regions/mutations is tested independently.

When all regions have been analyzed, the new set of mutations is compared to the initial set of mutations, and if there have been any changes, a new round begins, with each region going through the same process. This continues until, at the end of a full round, there have been no mutations added or removed. The code for this process can be found at https://github.com/ryhisner/MP_EPCI_code.

Due to the extreme density of mutations in the receptor-binding domain (RBM), from S:335-528, all mutations in that region were excluded. In addition, 98 mutations were excluded from the permutation-based test that follows below due to their miscategorization by NextClade or being common sequencing errors (Table S1). Inclusion of these mutations has no effect on the 14 MPs that were ultimately identified, but they produce a large number of artifactual or trivial MPs when testing all other EPCI mutations. In addition, all mutations in the overlapping parts of ORF9b and N that are simultaneously caused by a single nucleotide mutation were excluded. Finally, for any MP involving only two mutations, these mutations were required to co-occur in at least four EPCI sequences to qualify.

As a control of biases introduced during the initial stages of identifying MPs where seed mutations were selected based on how commonly they occurred in the EPCI dataset relative to the overall dataset of circulating SARS-CoV-2 sequences, all private mutations that occur at least five times in the EPCI dataset were tested in the same manner as the original seed-mutations, with each mutation serving as a seed mutation in the MPD function. This control test was called the all-mutations control test (AMCT).

To empirically estimate the probabilities of MPs of varying sizes arising by chance alone we devised a permutation test involving 100 simulated datasets designed to capture key features of the EPCI dataset but where the linkage of mutations within the real sequences were broken by randomly distributing mutations found in the EPCI dataset across the sequences. Specifically, in each one of these artificial EPCI datasets, (hereafter referred to as APCI datasets) a list of artificial sequences of the same lengths as corresponding sequences in the EPCI dataset was created. Each artificial sequence contained the same total number of private substitutions, private spike substitutions, private RBD substitutions, and private RBM substitutions as a corresponding sequence from the EPCI list. Each sequence was randomly assigned AA substitutions from each region from an array of all private AA substitutions in the EPCI dataset. After being used, a substitution was removed from the array, so that collectively, the substitutions assigned to the artificial sequences exactly match the private substitutions contained in the EPCI dataset. A total of 100 different APCI datasets were created and each subjected to the same statistical tests as in MPD1 and MPD2.

The results of the APCI simulations indicate a very low false-detection rate. Running the MDS function on the 100 APCI datasets returned 44 total MPs, or 0.44 per APCI dataset compared to 29 MPs detected in the EPCI dataset (after removal of all duplicate MPs and any MPs that shared more than half of their mutations with a larger MP).

Nearly all of these MPs consisted of just two mutations. The average APCI MP contained 2.45 mutations in 2.34 regions, compared to 10.48 mutations in 4.24 regions in the EPCI dataset. The EPCI dataset contains 20 MPs involving five or more mutations while the 100 APCI datasets produced zero MPs with at least five mutations. See Table S2a and S2b for detailed statistics and the corresponding results for the position-only version of the MPD.

The average and median number of non-RBD mutations in APCI MP sequences (25.21 and 23.32, respectively) were higher than in EPCI MP sequences (21.54 and 18.26), suggesting that the greater number of MPs detected in the EPCI dataset is not due to the uneven distribution of the number of mutations in EPCI sequences.

Detailed EPCI and APCI results can be found at https://github.com/ryhisner/MP_Substitutions Main_Results and https://github.com/ryhisner/MP_Positions Main_Results.

### Bronchoalveolar Lavage Mutational Signature Detection

One of the 14 MPs that were ultimately detected differs distinctly from the others. Its most prominent features are an extremely high concentration of mutations from E:5-42, specific mutations in NSP4 (especially S209F, S209P, T295I, A307V, and H313Y), a high density of M mutations, and a marked absence of mutations in the spike RBD—particularly the RBM—compared to other EPCI sequences. It also includes a large number of mutations scattered throughout the genome. Examining sequences marked by this pattern, it was observed that many of them were bronchoalveolar lavage (BAL) samples. After noticing this, a more comprehensive search among the EPCI sequences was performed, revealing a total of 41 BAL samples, nearly all of which possessed multiple mutations in this heretofore mysterious MP.

Since a large proportion of SARS-CoV-2 samples do not contain information about the sample source, many other sequences with the BAL MP may have in fact been BAL samples. We designed an objective way of characterizing the BAL MP using a custom Julia code.

The 41 known BAL-sample sequences were examined, and any mutation occurring in at least three of them and which occurred in the BAL sequences at a rate ≥3 times greater than it occurred in the EPCI list overall was included in the initial group of seed mutations. Any sequence containing at least two of these seed mutations was identified and grouped together with the 41 known BAL sequences. The private mutations in these sequences (as determined by Nextclade CLI) were tallied and tested for statistical significance. Any mutations meeting the statistical requirements described below were then added to the initial seed mutations and used to select a new group of sequences in the next round.

Throughout the entire, multi-round process, each mutation was required to have a Fisher’s exact test p-value of 0.001 (10^-3^). In order to select the mutations most highly associated with BAL sequences first, an initial Fisher’s exact test -log10 p-value of 6.0 was set. In each subsequent round, the minimum Fisher’s exact test threshold either decreased or remained constant, depending on the number of new mutations added in the previous round. If more than two new mutations were added, the Fisher’s exact test p-value threshold was held constant. If two or three mutations were added, it decreased by 0.1, and if zero or one mutation was added, it was lowered by 0.2, until a lower limit was reached. The primary analysis used a minimum Fisher’s exact test -log10 p-value threshold of 3.0, while higher and lower minimum values were utilized for sensitivity tests.

In order to exclude mutations that are statistically overrepresented in BAL-MP sequences but which are also very common in non-BAL-MP EPCI sequences, a minimum BAL-MP percentage of 45% was required for qualification. If fewer than 45% of the EPCI sequences a mutation occurs in are BAL sequences, then the mutation is disqualified. The minimum BAL-MP percentage is made to start out at a lower level (∼33%) and gradually rises as more mutations are added to the BAL-MP list, reaching its maximum level of 45% if/when the BAL-MP mutation list reaches 85 mutations.

To eliminate the bias inherent in testing the statistical overrepresentation of a given mutation in a set of sequences that were selected based, at least in part, on the presence or absence of that mutation, a separate list of sequences was created for each mutation where this dependency was removed. Specifically, for each list the mutation being considered was excluded from the criteria upon which the list of sequences was selected. For example, during this process a sequence was required to have two or more mutations from the previous round’s BAL-signature mutation list to qualify. However, if, for example, mutation A was selected as a BAL-associated mutation in the previous round(s), in the subsequent round(s) when testing mutation A, any sequence that possessed mutation A was required to also have at least two other mutations from the BAL-signature list: i.e. the presence of mutation A did not contribute to the count of mutations used to judge whether the sequence should be classified as having a BAL-associated MP. As in the primary MP coding function, a private mutations adjustment factor (PMA) was introduced to partially adjust for the number of private mutations in each sequence (see description above).

## RESULTS AND DISCUSSION

Given the strict limits placed on adaptive evolution in circulating viruses, including limited intrahost diversity and tight transmission bottlenecks, valleys in the fitness landscape requiring multiple simultaneous mutations can present formidable barriers to adaptive viral evolution. Recombination between genetically distant variants during coinfection of the same cell represents one means for escaping this evolutionary straightjacket. Infections of individual cells with divergent SARS-CoV-2 variants is rare: partly because SARS-CoV-2 infections are usually so short that people only rarely become infected with two genetically divergent variants, and partly because when individual cells are infected by a SARS-CoV-2 virion, they rapidly develop resistance to being infected by additional virions (something called superinfection exclusion)^70^. During chronic infections, however, more viral diversity will accumulate, fitter variants will have more time to rise in frequency, the chances of coinfections occurring will increase, the rates at which individual cells become infected with genetically distinct variants will rise^48^, and advantageous combinations of mutations (including mutations involving more than one nucleotide change in the same codon) will be far more likely to occur. Researchers examining repeated RT-PCR tests in a prospective, representative survey in the UK estimated that persistent infections lasting at least 60 days occur in 0.1-0.5% of all infections^256^.

We therefore undertook systematic searches to: (1) identify publicly available sequences that were likely derived from chronically infected individuals (referred to as EPCI sequences); (2) identify convergent mutations within the EPCI sequences and (3) identify patterns of co-occurrence of different convergent mutations between widely separated genome regions: patterns we refer to collectively as MPs.

We initially identified seven MPs through a preliminary manual examination of EPCI sequences we named NTD-DS, 8Fold, Rd3N, Y1-6, 2Macs, BAL, and 6-7a. Based on the manual exploratory process we designed a genome-region aware co-evolution and convergent evolution detection algorithm to more thoroughly search for additional MPs: an effort that revealed an additional seven MPs which we named 9b8, NuBS, D612, 1Macs, CWW, HR1M, and BsHel (Table 1).

**Table 1:** Genomic regions included in 11 of the 14 MPs and the number of EPCI sequences exhibiting each MP.

|  | Genome region number |  |  |  |  |  |  |  |  |
| --- | --- | --- | --- | --- | --- | --- | --- | --- | --- |
| Name | 1 | 2 | 3 | 4 | 5 | 6 | 7 | 8 | EPCI Count<sup>1</sup> |
| **8Fold** | ORF1a:1318-1326 | ORF1a:1638-1639 | ORF1a:2274-2280 | ORF1a:3441, 3443 | ORF1a:4085-4090 | ORF1a:4164-4167 | ORF1b:730-731 | ORF3a:211-215 | 435 |
| **9b8** | ORF1a:239 | ORF1a:3949-3952 | ORF9b:64-68 | ORF9b:92-95 |  |  |  |  | 81 |
| **NuBS** | ORF1a:2134-2138 | ORF1a:4083, 4087, 4090 | N:326 |  |  |  |  |  | 107 |
| **D612** | ORF1a:1500-1507 | ORF1a:3648-3657 | ORF1b:72-76 |  |  |  |  |  | 74 |
| **Rd3N** | ORF1a:1540-1543 | ORF1a:1566-1572 | ORF1a:4396-4398 | ORF1b:820-824 | N:270-271 |  |  |  | 165 |
| **Y1-6** | ORF1a:2466-2472, 2475 | ORF1a:3802-3203, 3808 |  |  |  |  |  |  | 72 |
| **2Macs** | ORF1a:232-233 | ORF1a:1272-1276 | ORF1a:1361-1372 |  |  |  |  |  | 85 |
| **6-7a** | ORF1a:1706,1709,1714,1715 | ORF1a:1769, 1771, 1773 | ORF1a:K1795Q | ORF1a:2209, 2214-2215 | ORF1a:2324-2329 | ORF1a:2345-2353 | ORF7a:22 | ORF7a:39, 43 | 107 |
| **1Macs** | ORF1a:110, 114 | ORF1a:959 | ORF1a:1220-1224 | ORF1a:1394-1397 |  |  |  |  | 62 |
| **BsHel** | ORF1a:2165-2166 | ORF1b:1220 |  |  |  |  |  |  | 34 |
| **HR1M** | S:936 | ORF3a:255-258 | M:155 | ORF1a:3201 |  |  |  |  | 49 |
| **BAL** | Various<sup>2</sup> |  |  |  |  |  |  |  | 467 |
| **CWW** | Various<sup>2</sup> |  |  |  |  |  |  |  | 22 |
| **NTD-DS** | S:9, 12-17, 21, 64, 136-145, 151-152, 258 |  |  |  |  |  |  |  | 136 |

<sup>1</sup> Total MP sequences in the EPCI dataset. A sequence was required to have two amino acid mutations at the indicated positions in MPs with five or fewer regions, and three amino acid mutations at the indicated positions for MPs with more than five regions. The number of qualifying MP sequences for the BAL and CWW MPs were determined similarly but using specific mutations (rather than all mutations at a given site). NTD-DS sequences were required to have two NTD disulfide-altering mutations (listed in the NTD-DS section).
<sup>2</sup> The BAL MP involves over 80 mutations in dozens of regions and the CWW MP includes at least 17 separate regions (see Tables 5 and 14).

**Table 2:** Full list of sites where substitutions occur significantly more frequently in EPCI sequences with a 8fold-MP than they do in other EPCI sequences.

| Genome region # | Substitution position | Adjusted % frequency in 8fold EPCI sequences<sup>1</sup> | Adjusted % frequency in non-8fold EPCI sequences<sup>2</sup> | Fold Increase<sup>3</sup> | Enrichment p-value<sup>4</sup> |
| --- | --- | --- | --- | --- | --- |
| 1<br>(8z1) | ORF1a:1318 | 1.99% | 0.37% | 5.40 | 5.87 x 10<sup>-3</sup> |
|  | ORF1a:1322 | 51.67% | 7.32% | 7.06 | 3.44 x 10<sup>-77</sup> |
|  | ORF1a:1323 | 29.33% | 4.54% | 6.46 | 8.54 x 10<sup>-38</sup> |
|  | ORF1a:1324 | 8.33% | 2.02% | 4.13 | 1.16 x 10<sup>-7</sup> |
|  | ORF1a:1326 | 3.33% | 0.27% | 12.34 | 1.69 x 10<sup>-6</sup> |
|  | ORF1a:1328 | 1.00% | 0.00% | >29.67 | 1.53 x 10<sup>-3</sup> |
| 2<br>(8z2) | ORF1a:1637 | 2.42% | 0.61% | 4.00 | 9.51 x 10<sup>-3</sup> |
|  | ORF1a:1638 | 76.66% | 12.55% | 6.11 | 1.20 x 10<sup>-117</sup> |
|  | ORF1a:1639 | 21.60% | 3.67% | 5.89 | 1.07 x 10<sup>-24</sup> |
| 3<br>(8z3) | ORF1a:2274 | 6.48% | 1.09% | 5.95 | 2.79 x 10<sup>-11</sup> |
|  | ORF1a:2279 | 1.02% | 0.07% | 13.97 | 2.54 x 10<sup>-3</sup> |
|  | ORF1a:2280 | 1.02% | 0.07% | 14.00 | 2.52 x 10<sup>-3</sup> |
| 4<br>(8z4) | ORF1a:3441 | 1.58% | 0.14% | 10.98 | 1.68 x 10<sup>-4</sup> |
|  | ORF1a:3443 | 3.56% | 0.04% | 98.98 | 5.88 x 10<sup>-14</sup> |
| 5<br>(8z5) | ORF1a:4085 | 1.48% | 0.14% | 10.38 | 4.87 x 10<sup>-4</sup> |
|  | ORF1a:4086 | 1.27% | 0.11% | 11.89 | 9.89 x 10<sup>-4</sup> |
|  | ORF1a:4087 | 16.56% | 2.67% | 6.19 | 4.66 x 10<sup>-28</sup> |
|  | ORF1a:4088 | 6.14% | 0.18% | 34.39 | 5.36 x 10<sup>-20</sup> |
|  | ORF1a:4090 | 17.58% | 1.32% | 13.33 | 3.94 x 10<sup>-44</sup> |
| 6<br>(8z6) | ORF1a:4164 | 19.22% | 2.39% | 8.04 | 2.55 x 10<sup>-33</sup> |
|  | ORF1a:4165 | 24.80% | 1.73% | 14.31 | 1.80 x 10<sup>-56</sup> |
|  | ORF1a:4166 | 19.74% | 1.28% | 15.39 | 2.32 x 10<sup>-45</sup> |
|  | ORF1a:4167 | 5.68% | 0.31% | 18.25 | 3.31 x 10<sup>-14</sup> |
| 7<br>(8z7) | ORF1b:730 | 11.65% | 0.83% | 14.10 | 1.13 x 10<sup>-30</sup> |
|  | ORF1b:731 | 0.99% | 0.07% | 13.85 | 2.64 x 10<sup>-3</sup> |
|  | ORF1b:734 | 1.00% | 0.11% | 9.25 | 6.11 x 10<sup>-3</sup> |
| 8<br>(8z8) | ORF3a:210 | 1.84% | 0.22% | 8.47 | 1.57 x 10<sup>-4</sup> |
|  | ORF3a:211 | 1.43% | 0.22% | 6.60 | 2.50 x 10<sup>-3</sup> |
|  | ORF3a:213 | 12.94% | 2.39% | 5.41 | 2.61 x 10<sup>-20</sup> |
|  | ORF3a:215 | 1.64% | 0.11% | 15.11 | 5.29 x 10<sup>-5</sup> |

<sup>1</sup> Total proportion of EPCI sequences with the 8fold-MP (after excluding the mutation under consideration from the identification of sequences with an 8fold-MP) and accounting for sequences with dropout or ambiguous nucleotides at the site under consideration.
<sup>2</sup> Total proportion of EPCI sequences without the 8fold-MP (after excluding the mutation under consideration from the identification of sequences with an 8fold-MP) and accounting for sequences with dropout or ambiguous nucleotides at the site under consideration.
<sup>3</sup> Fold increase of the mutation in EPCI sequences with the 8fold-MP relative to the mutation's frequency in EPCI sequences without the 8fold-MP.
<sup>4</sup> Unadjusted Fisher's Exact test p-value.

**Table 3:** Full list of substitutions that occur significantly more frequently in EPCI sequences with an 8fold-MP than they do in other EPCI sequences.

| MP region # | Substitution | Adjusted % frequency in 8fold EPCI sequences<sup>1</sup> | Adjusted % frequency in non-8fold EPCI sequences<sup>2</sup> | Fold Increase<sup>3</sup> | Enrichment p-value<sup>4</sup> |
| --- | --- | --- | --- | --- | --- |
| 1 (8z1) | ORF1a:R1318G | 1.97 | 0.37 | 5.32 | 6.28 x 10<sup>-3</sup> |
|  | ORF1a:T1322K | 0.99 | 0.00 | >29.45 | 1.57 x 10<sup>-3</sup> |
|  | ORF1a:T1322A | 6.60 | 0.98 | 6.77 | 5.37 x 10<sup>-9</sup> |
|  | ORF1a:T1322I | 43.89 | 5.75 | 7.63 | 8.26 x 10<sup>-67</sup> |
|  | ORF1a:D1323G | 12.58 | 2.56 | 4.92 | 4.60 x 10<sup>-13</sup> |
|  | ORF1a:D1323N | 12.91 | 1.72 | 7.52 | 5.67 x 10<sup>-18</sup> |
|  | ORF1a:N1324H | 1.33 | 0.07 | 19.82 | 1.79 x 10<sup>-3</sup> |
|  | ORF1a:N1324D | 1.33 | 0.07 | 19.82 | 1.79 x 10<sup>-3</sup> |
|  | ORF1a:N1324S | 3.33 | 0.71 | 4.72 | 5.57 x 10<sup>-4</sup> |
|  | ORF1a:I1326V | 1.99 | 0.14 | 14.70 | 1.83 x 10<sup>-4</sup> |
|  | ORF1a:I1326T | 1.66 | 0.07 | 24.50 | 2.37 x 10<sup>-4</sup> |
|  | ORF1a:T1328N | 0.99 | 0.00 | >29.13 | 1.61 x 10<sup>-3</sup> |
| 2 (8z2) | ORF1a:T1638N | 15.11 | 3.12 | 4.84 | 1.90 x 10<sup>-14</sup> |
|  | ORF1a:T1638A | 2.88 | 0.40 | 7.15 | 2.57 x 10<sup>-4</sup> |
|  | ORF1a:T1638I | 57.55 | 8.72 | 6.60 | 7.20 x 10<sup>-80</sup> |
|  | ORF1a:D1639A | 1.78 | 0.17 | 10.59 | 1.62 x 10<sup>-3</sup> |
|  | ORF1a:D1639N | 19.57 | 2.89 | 6.78 | 3.81 x 10<sup>-24</sup> |
| 3 (8z3) | ORF1a:T2274I | 5.56 | 1.03 | 5.41 | $2.62 \times 10^{-9}$ |
| | ORF1a:A2279V | 0.79 | 0.00 | >21.63 | $1.18 \times 10^{-3}$ |
| | ORF1a:T2280I | 0.60 | 0.00 | >16.25 | $7.53 \times 10^{-3}$ |
| 4 (8z5) | ORF1a:T4087I | 11.55 | 2.55 | 4.54 | $1.43 \times 10^{-15}$ |
| | ORF1a:T4087A | 1.89 | 0.18 | 10.56 | $5.56 \times 10^{-5}$ |
| | ORF1a:T4088I | 5.15 | 0.29 | 17.95 | $9.50 \times 10^{-15}$ |
| | ORF1a:T4090A | 1.25 | 0.00 | >34.88 | $1.95 \times 10^{-5}$ |
| | ORF1a:T4090I | 14.97 | 1.36 | 11.01 | $4.82 \times 10^{-35}$ |
| 5 (8z6) | ORF1a:T4164N | 10.10 | 1.08 | 9.34 | $1.30 \times 10^{-19}$ |
| | ORF1a:T4164I | 7.64 | 1.33 | 5.76 | $2.45 \times 10^{-11}$ |
| | ORF1a:D4165N | 5.24 | 0.52 | 10.07 | $1.75 \times 10^{-10}$ |
| | ORF1a:D4165G | 12.83 | 0.83 | 15.42 | $2.18 \times 10^{-29}$ |
| | ORF1a:D4165V | 1.57 | 0.00 | >45.33 | $4.93 \times 10^{-6}$ |
| | ORF1a:D4165A | 4.97 | 0.28 | 17.94 | $2.35 \times 10^{-12}$ |
| | ORF1a:D4166G | 9.55 | 0.48 | 19.85 | $1.01 \times 10^{-22}$ |
| | ORF1a:D4166A | 1.40 | 0.10 | 13.62 | $1.27 \times 10^{-3}$ |
| | ORF1a:D4166N | 7.87 | 0.41 | 19.07 | $1.37 \times 10^{-18}$ |
| | ORF1a:D4166S | 0.84 | 0.00 | >24.52 | $2.57 \times 10^{-3}$ |
| | ORF1a:D4166Y | 1.69 | 0.10 | 16.35 | $2.03 \times 10^{-4}$ |
| | ORF1a:N4167S | 3.10 | 0.04 | 88.64 | $5.15 \times 10^{-11}$ |
| 6 (8z7) | ORF1b:T730I | 11.18 | 0.87 | 12.91 | $6.53 \times 10^{-29}$ |
| 7 (8z8) | ORF3a:Q213K | 10.87 | 2.11 | 5.14 | $6.98 \times 10^{-17}$ |
| | ORF3a:Y215H | 1.18 | 0.04 | 32.49 | $1.70 \times 10^{-4}$ |

<sup>1</sup> Total proportion of EPCI sequences with the 8fold-MP (after excluding the mutation under consideration from the identification of sequences with an 8fold-MP) and accounting for sequences with dropout or ambiguous nucleotides at the site under consideration.
<sup>2</sup> Total proportion of EPCI sequences without the 8fold-MP (after excluding the mutation under consideration from the identification of sequences with an 8fold-MP) and accounting for sequences with dropout or ambiguous nucleotides at the site under consideration.
<sup>3</sup> Fold increase of the mutation in EPCI sequences with the 8fold-MP relative to the mutation's frequency in EPCI sequences without the 8fold-MP.
<sup>4</sup> Unadjusted Fisher's Exact test p-value.

There are three points worth stressing about these fourteen MPs: (1) the convergent evolution evident at many of the individual genome sites represented within each of the MPs is strongly suggestive of selection-driven adaptation to some specific niche or combination of niches); (2) the substantially lower frequencies of the individual mutations comprising these MPs in non-EPCI sequences is indicative of the mutations being selectively more favourable during chronic infections than they are during acute infections; and (3) the high frequencies of co-occurrence of groups of mutations at widely separated genome sites within individual MPs is suggestive of extensive coevolution between genome sites during adaptation to these chronic-infection specific niches. Below we attempt to infer, given the observed mutations comprising each MP, the genome regions involved and our current understanding of how these genome regions might be biologically connected in the context of the specific niches within which each of the MPs evolved.

### CD8 T-cell Escape-Associated Mutation Patterns (8Fold, 9b8, NuBS, D612)

Selection pressures within a persistently infected host may differ in several ways from those present in acute infections. In circulating viruses, for example, there is little selection pressure to evade T cell immunity. Given the existence of hundreds of different HLA types among the human population, mutations in CD4 or CD8 T cell epitopes will be unlikely to evade immunodominant T cell responses in more than a modest proportion of the population^71^. Furthermore, early in the pandemic, as much as half of SARS-CoV-2 transmission occurred in the presymptomatic phase^72–73^. Though this fraction has likely dropped as population immunity has spread, it is still likely that most transmission takes place prior to onset of the T cell-driven arm of the adaptive immune response^71^. In addition, while individual spike mutations can sometimes lead to dramatic antibody escape^74^, the number and diversity of T cell epitopes make escape more difficult, with one study estimating that escape mutations in eight T cell epitopes are required to escape 80% of the CD4 and CD8 T cell response^75^.

Substantial T cell escape during acute infection is therefore extremely unlikely to occur. However, in chronic infection, particularly within immunocompromised hosts lacking a sufficient antibody response, T cell evasion can become a major driver of adaptive evolution^76–79^. Furthermore, there are case studies in which chronic infection led to extensive mutations at immunodominant T cell epitopes^11, 286^. Accordingly, chronic infection associated MPs that we have identified include four MPs that are plausibly indicative of escape from common HLA types.

### The 8Fold MP is likely associated with HLA-A*01:01 escape

A*01:01 is one of the best-studied HLA types, and is the second most common HLA type globally^80^. Immunodominant A*01:01 epitopes in the SARS-CoV-2 genome have been identified in multiple studies^81–83^, and the results of CD8 T cell epitope studies from 2020 and 2021 have been comprehensively summarized in Grifoni et al., 2021^71^.

One of the most extensive and frequent MPs observed in EPCI sequences is The Eightfold Way (8Fold), so named because it involves eight distinct genomic regions. While the exact boundaries of each of the eight regions are somewhat vague , the vast majority of 8Fold mutations occur in the following eight regions: ORF1a:1322-1324 (8z1), ORF1a:1638-1639 (8z2), ORF1a:2274-2280 (8z3), ORF1a:3441-3443 (8z4), ORF1a:4087-4090 (8z5), ORF1a:4164-4167 (8z6), ORF1b:730 (8z7), and ORF3a:211-215 (8z8) (Figure 1). We discovered and documented the 8Fold MP prior to discovering that all of its eight regions are HLA*01:01 CD8 T cell epitopes; 7/8 of which are immunodominant. Furthermore, the two most extensively documented and most immunodominant epitopes (ORF1a:1321-1329 and ORF1a:1637-1646) are the two most common 8Fold mutations in EPCI sequences.

**Figure 1:**
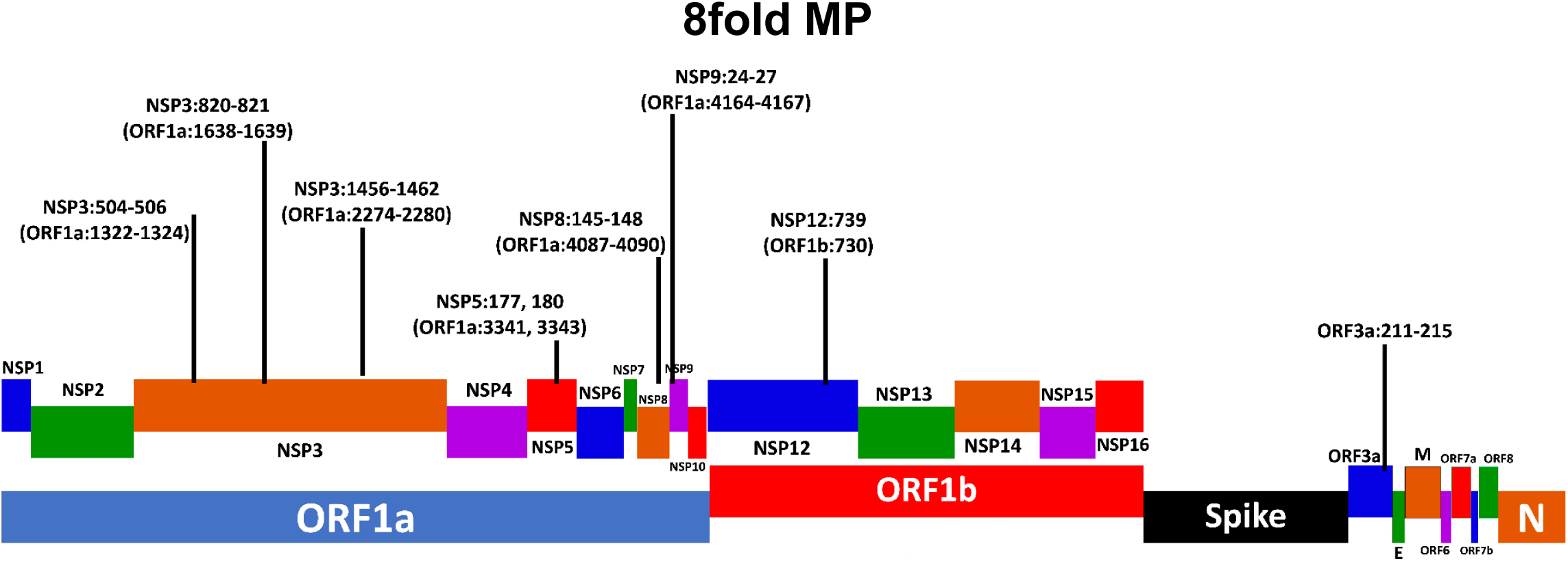
Schematic representation of the SARS-CoV-2 genome with the 8fold MP-associated mutations labeled.

The most common of all 8Fold MP mutations is ORF1a:T1638I, which occurs in the papain-like protease (PLpro) region of NSP3. In addition to being a CD8 T cell-escape mutation, it may also confer another intrahost advantage. PLpro cleaves ISG15 and K48-linked ubiquitin chains from proteins^84^. The SARS-CoV-1 and SARS-CoV-2 PLpro domains cleave ISG15 at comparable rates. But the SARS-CoV PLpro’s deubiquitination (DUB) activity is far greater than that of the SARS-CoV-2 PLpro. This difference has largely been traced to the ORF1a:1795 residue, which is lysine (K) in SARS-CoV-2 but glutamine (Q) in all other sarbecoviruses.

ORF1a:K1795Q was shown to dramatically increase the SARS-CoV-2 PLpro’s DUB activity, to a level that even exceeded that of SARS-CoV-1^85^. Notably, one of the most common mutations in EPCI sequences is ORF1a:K1795Q, a reversion to the universal sarbecovirus residue. However, ORF1a:1638, which is threonine (T) in SARS-CoV-2, is leucine (L) in SARS-CoV-1, and ORF1a:T1638L was shown to more than double the DUB activity of PLpro in SARS-CoV-2^86^. Like ORF1a:T1638L, ORF1a:T1638I is a mutation to a more hydrophobic residue and could increase DUB activity, which appears to be advantageous within chronically infected hosts.

In an in silico prediction of ORF1ab epitopes with the highest binding to molecules from 61 different HLA class I types, eight of the top 30 predicted ORF1ab epitopes were A*01:01 epitopes, including five of the seven ORF1ab 8fold regions^87^. Similar results have been seen in physiological measurements in convalescent patients^82, 88^. In a 2021 meta-analysis of T cell epitopes, seven of 27 epitopes identified in four or more studies were HLA*01:01, with six of those including 8fold-MP regions^71^, all of which are immunodominant. Another study identified eight A*01:01 CD8 T cell epitopes, seven of them including 8fold-MP regions^89^. In another study looking at 18 convalescent patients, the two highest-ranking HLA scores, four of the top six, six of the top 21, and seven of the top 28 epitopes included 8fold-MP regions^90^.

One remarkable, documented chronic-infection displaying the 8fold pattern was a BA.5.11 from California, USA^273^, which, in addition to having nine mutations across the 8fold regions, also had N:D81H, a mutation in a highly conserved region of N that was identified in two studies as an immunodominant A*01:01 epitope^71, 75, 81^,.

Mutations in 8z1 (8fold MP region ORF1a:1322-1324), and 8z2 (8fold MP region ORF1a:1638-1639) show a particularly strong connection; of 633 EPCI sequences with at least one mutation in 8z1, 586 (92.6%) also have a mutation in 8z2. Furthermore, when looking exclusively at EPCI sequences with at least two private mutations in 8fold-MP regions, 586/599 (98.0%) with a mutation in 8z1 also have at least one mutation in 8z2. In one study that looked at 35 patients who had recovered from COVID-19, the most commonly detected and highest magnitude responses were to 8z1 and 8z2, with 8z2 being the top ranked by both measures. This mirrors the frequency of mutations in EPCI sequences, where mutations in 8z2 are more common than any other region of the genome, with ORF1a:1638 being the most frequently mutated position (585 sequences) in the entire genome, while ORF1a:1639 is the 17th-most mutated site (171) outside of spike. ORF1a:1322 is the 2nd-most mutated (371) and ORF1a:1323 the tenth-most mutated site outside of spike (216).

The immunodominance of 8z1 and 8z2 is also illustrated by their near-universal presence in sequences with three or more 8fold mutations. Of 437 EPCI sequences with mutations in three or more 8fold regions, 417 (95.4%) have mutations in both 8z1 and 8z2, and 432 (98.9%) have a mutation in 8z2. When ranking associated EPCI mutations by their Fisher’s Exact Test p-values, seven of the top 14 involve ORF1a:T1638I (Table S4).

### The 9b8 MP is potentially associated with HLA-B*18:01 escape

The 9b8 (ORF**<u>9b</u>**, NSP**<u>8</u>**) MP includes the regions, ORF9b:64-68, ORF9b:92-96, and ORF1a:3949-3952 (Figure 2). While ORF9b has been largely ignored in most T cell-epitope discovery studies, in one study, ORF9b:86-96 was found to be the most reactive epitope in the entire SARS-CoV-2 genome when exposed to PBMCs from A*02:01-positive individuals (the most common HLA-A type globally) who had recovered from SARS-CoV-2 infection^91^. Strikingly, all three 9b8 regions were identified as B*18:01 CD8 T cell epitopes^91^. ORF1a:A239V is also associated with the 9b8 MP, with seven of eight EPCI sequences with ORF1a:A239V also possessing at least one 9b8 mutation. ORF1a:232-242 is among the top predicted B*18:01 epitopes in a comprehensive in silico examination of potential spike and ORF1ab epitopes.^87^

**Figure 2:**
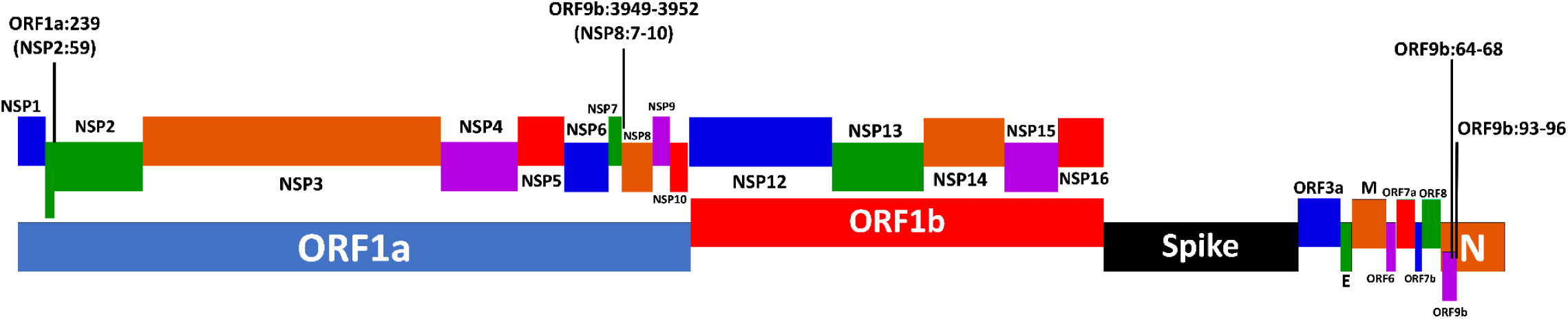
Schematic representation of the SARS-CoV-2 genome with the 9b8 MP-associated mutations labeled.

### The NuBS MP is likely associated with HLA-A*26:01 escape

The NuBS (**<u>Nu</u>**cleocapsid, NSP3 **<u>B</u>**etacoronavirus-**<u>S</u>**pecific Domain) MP consists of N:326 and ORF1a:2134-2136 (Figure 3). Both regions are located within known A*26:01 T cell epitopes (N:323-333 and ORF1a:2132-2141), with the latter being immunodominant. A third region containing mutations that modestly but clearly tend to co-occur with mutations occurring in the two NuBS regions, is ORF1a:2592-2597, which also sits within identified A*26:01 epitopes (ORF1a:2585-2594 and ORF1a:2594-2602).

**Figure 3:**
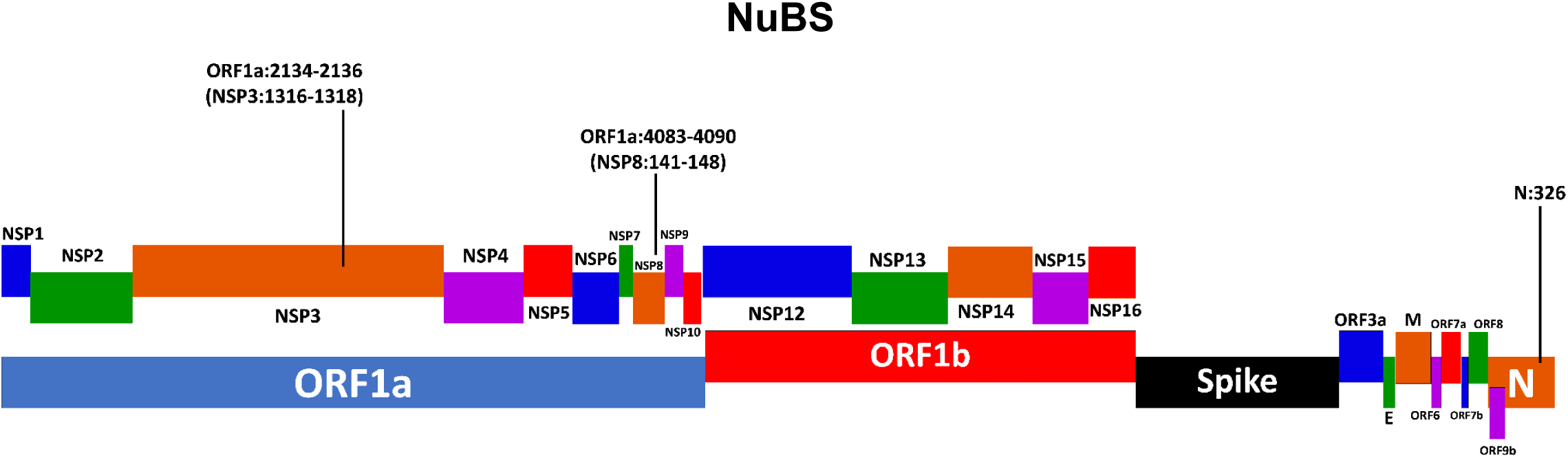
Schematic representation of the SARS-CoV-2 genome with the NuBS MP-associated mutations labeled.

One immunocompromised patient (IgG4-related disease, rituximab therapy) infected with an FL.1.5.1 variant for a minimum of 143 days exhibited multiple NuBS-MP mutations across five sequences^36^. All five sequences had N:P326L, while two had ORF1a:T4090I and three ORF1a:T4087I. Interestingly, three sequences featured different mutations in the other NuBS region: ORF1a:P2134S, D2136A, and Δ2131-2139, suggesting that multiple intrahost variants had each evaded CD8 T cell responses independently, by acquiring different escape mutations^36^.

### The D612 MP is potentially associated with HLA- B*38:01 and/or HLA-B*39:01 escape

The D612 MP (NSP3 **<u>D</u>**PUP, NSP**<u>6</u>**, NSP**<u>12</u>**) involves ORF1a:1500-1507, ORF1a:3648-3653, and ORF1b:72-76 (Figure 4). Peptides from ORF1a:1498-1508 and ORF1b:72-82 were identified through in silico investigation to have strong binding to B*38:01 and B*39:01^87^.

**Figure 4:**
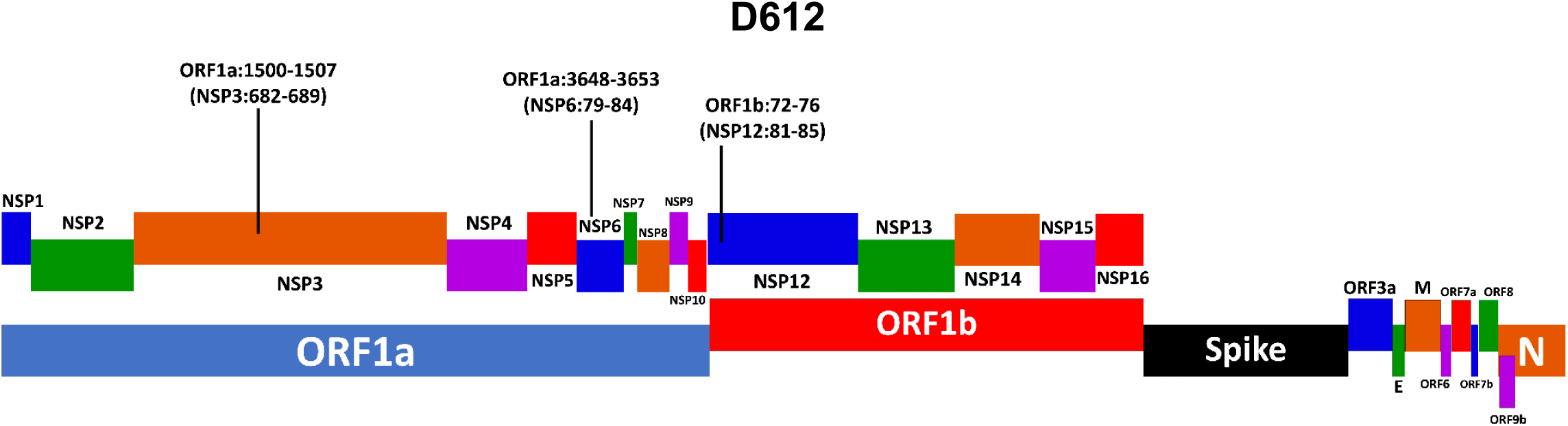
Schematic representation of the SARS-CoV-2 genome with the D612 MP-associated mutations labeled.

Key mutations comprising the BAL MP were initially detected through manual examination of EPCI sequences that were known to be derived from bronchoalveolar lavage samples. Following refinement of the BAL MP via a systematic automated search for sequences carrying BAL MP associated mutations (see Methods), it became apparent that the most distinctive characteristics of the finally determined BAL MP (Figures 5-7) are high substitution concentrations in the envelope (E) protein and the membrane (M) protein, a large number of substitutions at several NSP4 residues, and a marked absence of mutations in the spike receptor binding domain (RBD, S:335-528). The most common spike mutations in sequences carrying the BAL-MP are not concentrated in any one region, and include S50L, P330S, Y508H, T573I, and A653V. Two regions where BAL spike mutations do cluster to some degree are S:367-377 and S:1169-1186. Several mutations in the latter region, most notably L1186F, occur almost exclusively in EPCI sequences that clearly carry the BAL MP.

One well documented chronic infection involving a patient with B-cell lymphoma undergoing B cell-depletion therapy exhibited numerous classic BAL-MP mutations^284–285^ (Table 4). Three lower endotracheal samples from this patient (EPI_ISL_516999, EPI_ISL_517000, EPI_ISL_529236) featured 12 private amino acid substitutions (determined by placement on the global Usher tree^68^), six of which are BAL-MP mutations, including three of the top four and four and four of the top 10 mutations with the strongest statistical association with the BAL-MP (as measured by Fisher’s Exact test). Furthermore, another mutation in these sequences, ORF1a:I3835M (NSP6:I266M) occurs at a position that is mutated in just six EPCI sequences, five of which contain multiple BAL-MP mutations.

**Figure 5:**
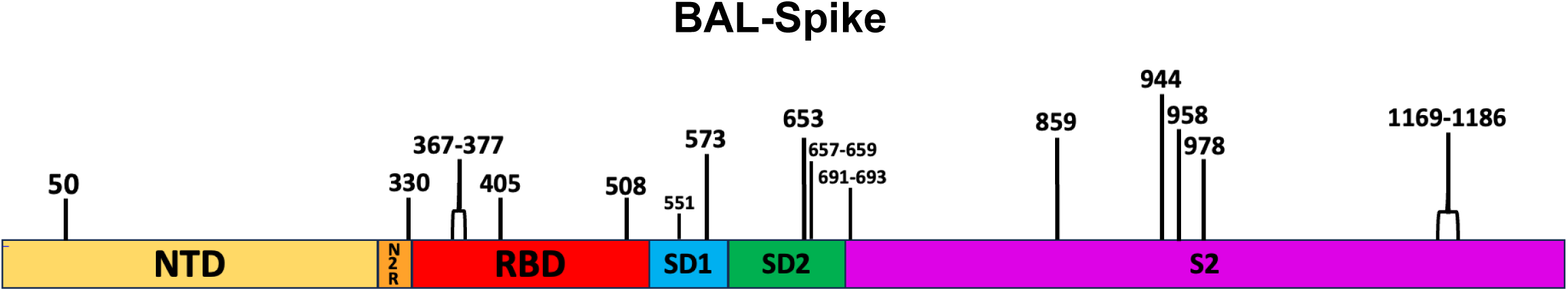
Schematic representation of the SARS-CoV-2 spike gene with BAL MP-associated mutations labeled.

**Figure 6:**
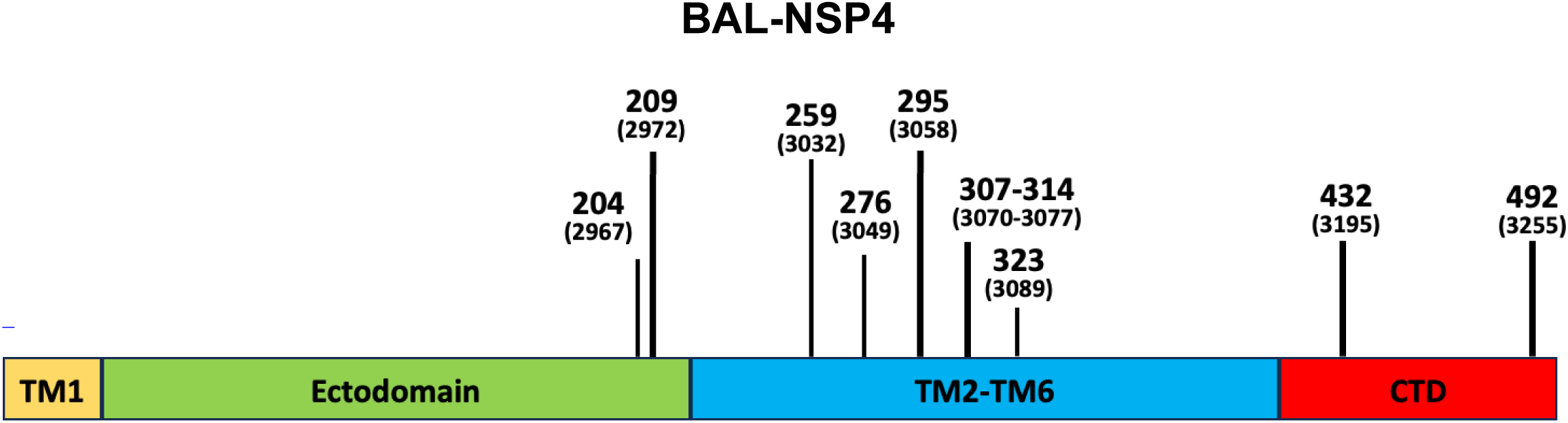
Schematic representation of the SARS-CoV-2 NSP4 gene with BAL MP-associated mutations labeled. TM1 = transmembrane region 1; TM2-TM6 = transmembrane regions 2 through 6; CTD = C-terminal domain

**Figure 7:**
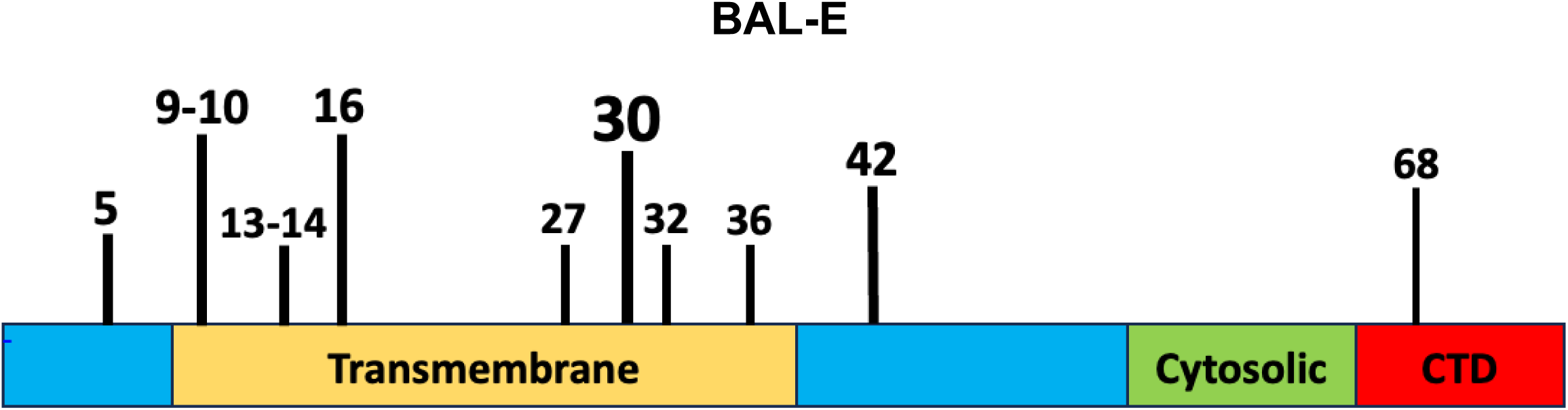
Schematic representation of the SARS-CoV-2 envelope gene with BAL MP-associated mutations labeled. Cytosolic = cytosolic domain; CTD = C-terminal domain^101^

**Figure 8:**
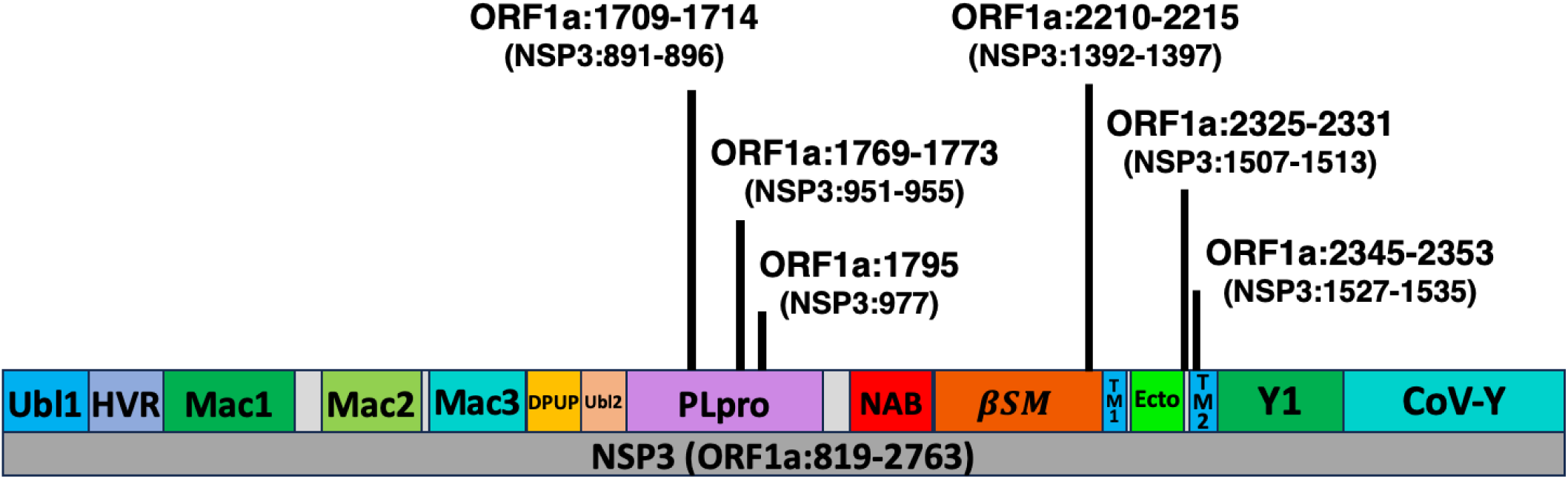
Schematic representation of NSP3 6-7a MP-associated mutations labeled. Ubl1 = ubiquitin-like domain 1; HVR = hypervariable region (also called acidic domain); Mac1 = macrodomain 1; Mac2 = macrodomain 2; Mac3 = macrodomain 3; DPUP = Domain preceding Ubl2 and PLpro; Ubl2 = ubiquitin-like domain 2; PLpro = papain-like protease; NAB = nucleic acid-binding domain; βSM = betacoronavirus-specific marker domain; TM1 = transmembrane domain 1; Ecto = ectodomain; TM2 = transmembrane domain 2; Y1 = Nidovirus-conserved domain of unknown function; CoV-Y = Coronavirus-specific C-terminal domain

**Figure 9:**
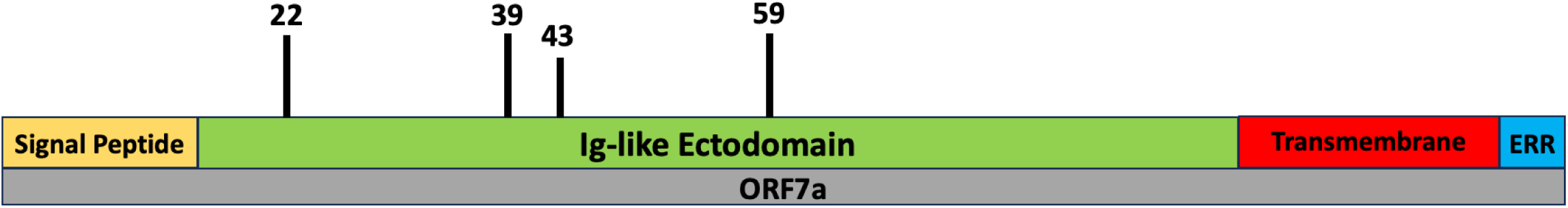
Schematic representation of the SARS-CoV-2 ORF7a gene with 6-7a MP-associated mutations labeled. Ig-like Ectodomain = immunoglobulin-like ectodomain; ERR = C-terminal ER-retention Motif‘^182^

**Figure 10:**
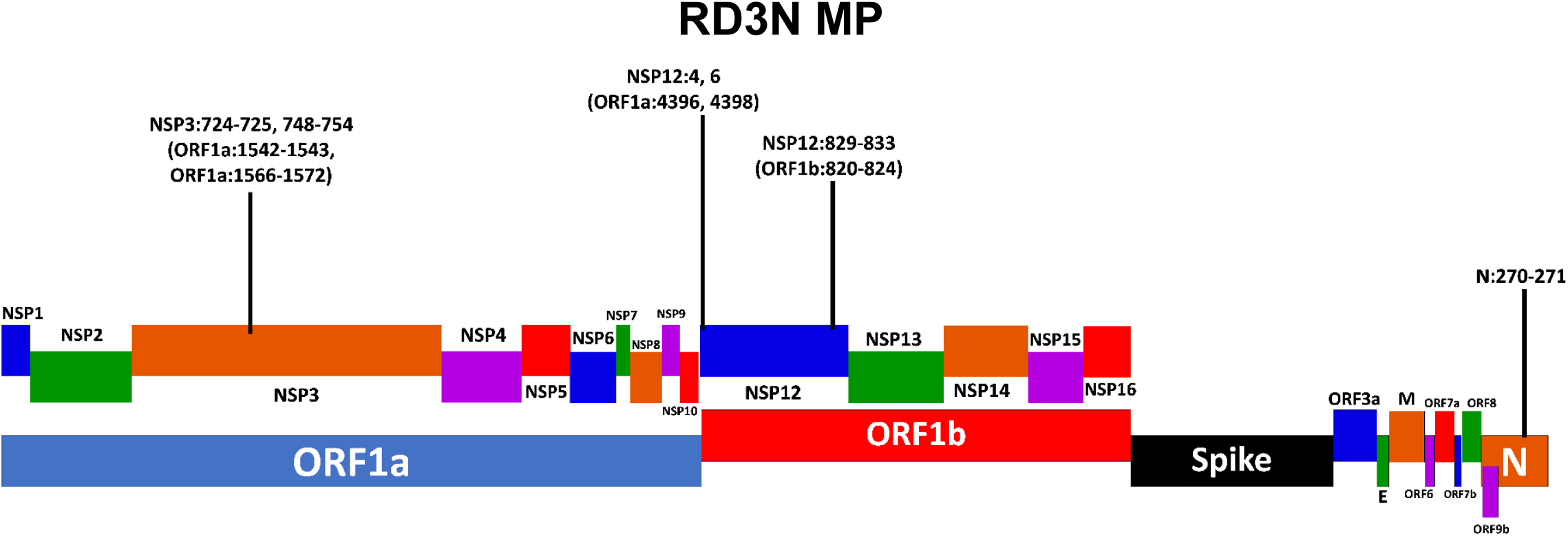
Schematic representation of the SARS-CoV-2 genome with the 8fold MP-associated mutations labeled.

**Table 4:** Mutations in chronic infection documented by Kemp et al, 2021. ^284^

| Gene | Private amino acid substitutions in virus from chronically infected lymphoma patient (BAL-MP mutations in bold) |
| --- | --- |
| Spike | S13I, W64G, **P330S** |
| E | **L19I, T30I** |
| M | **H125Y** |
| ORF1a | S1218N, **T3058I**, D3479N, **I3835M** |
| ORF1b | V783I, **Y1247C** |

**Table 5:**
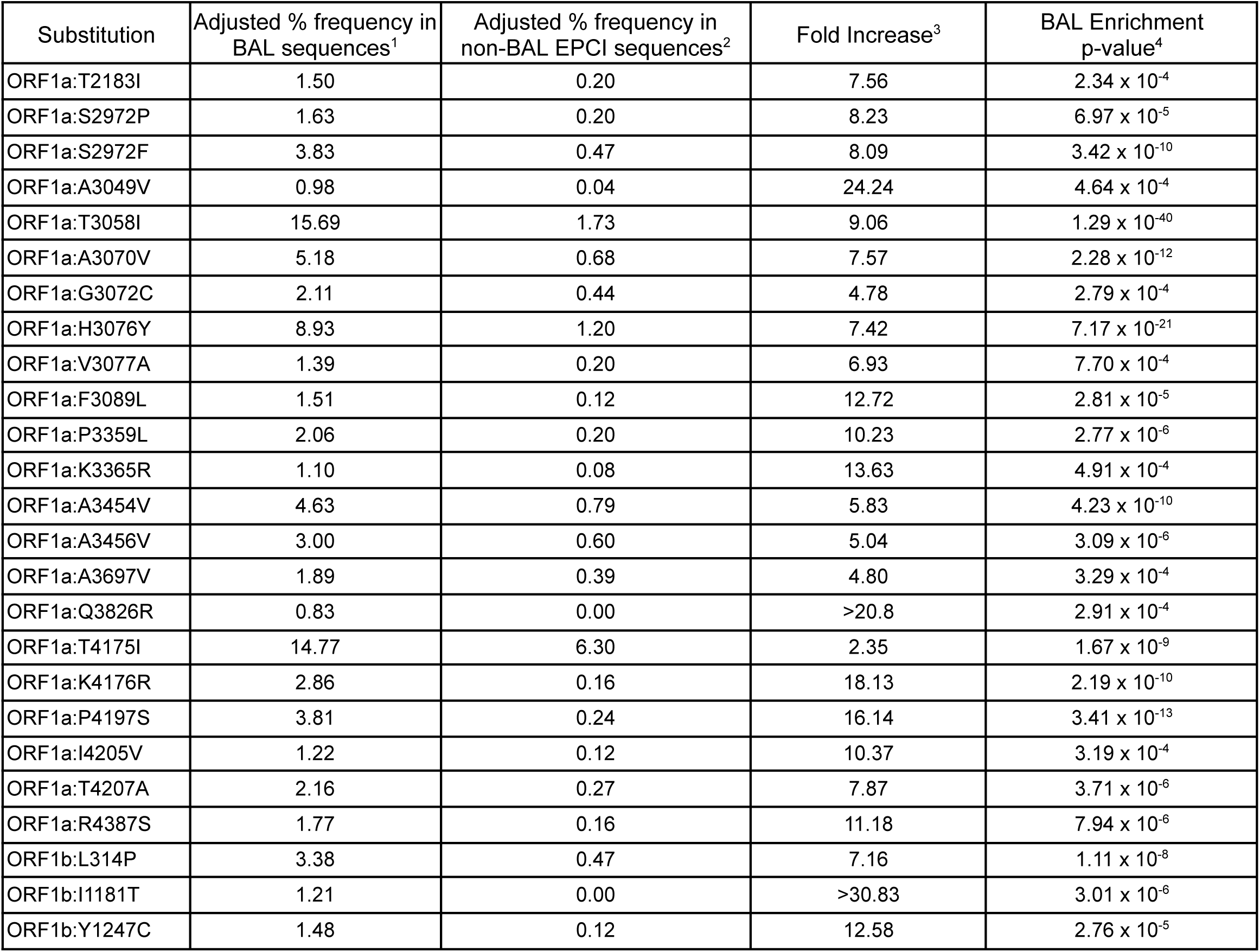
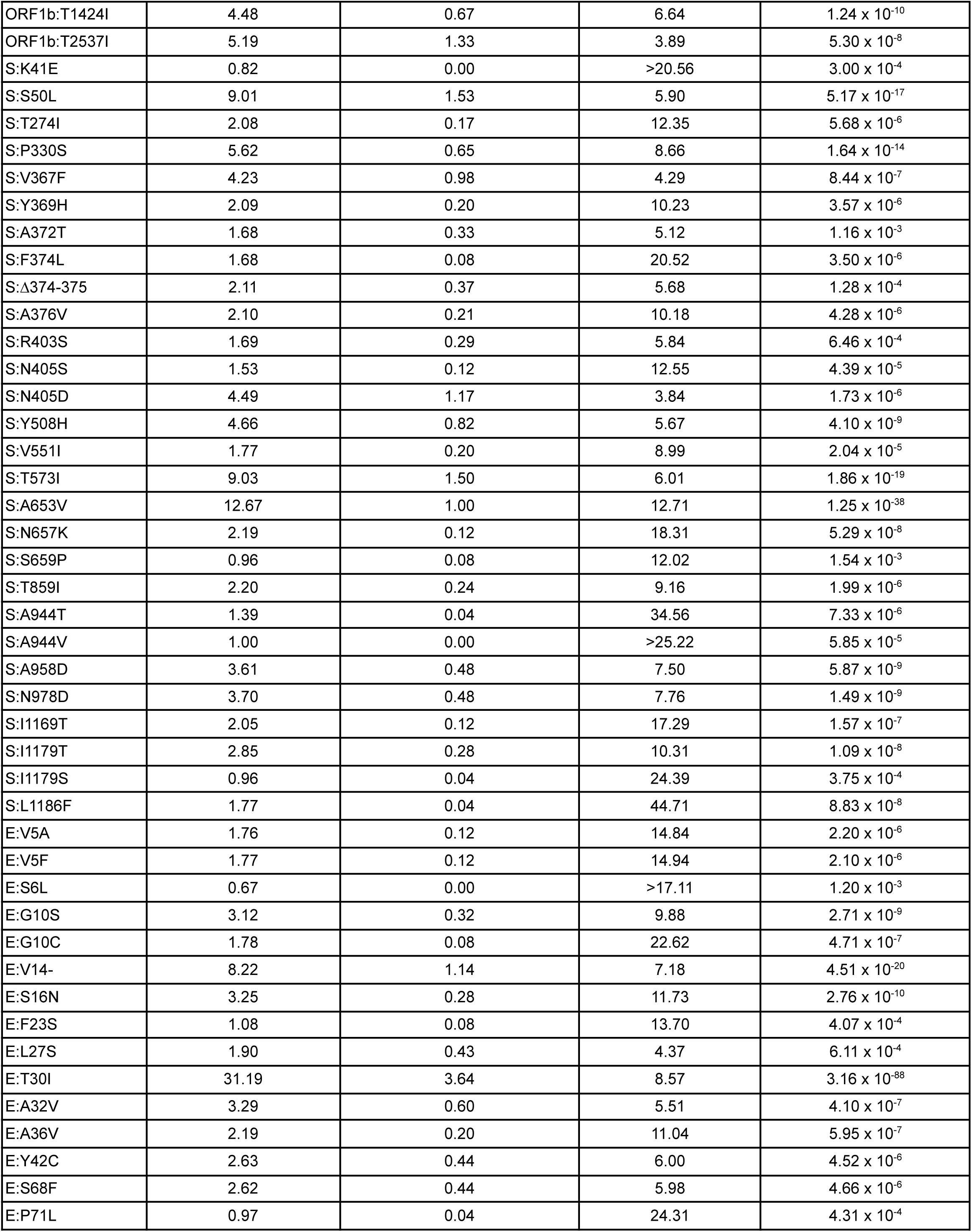
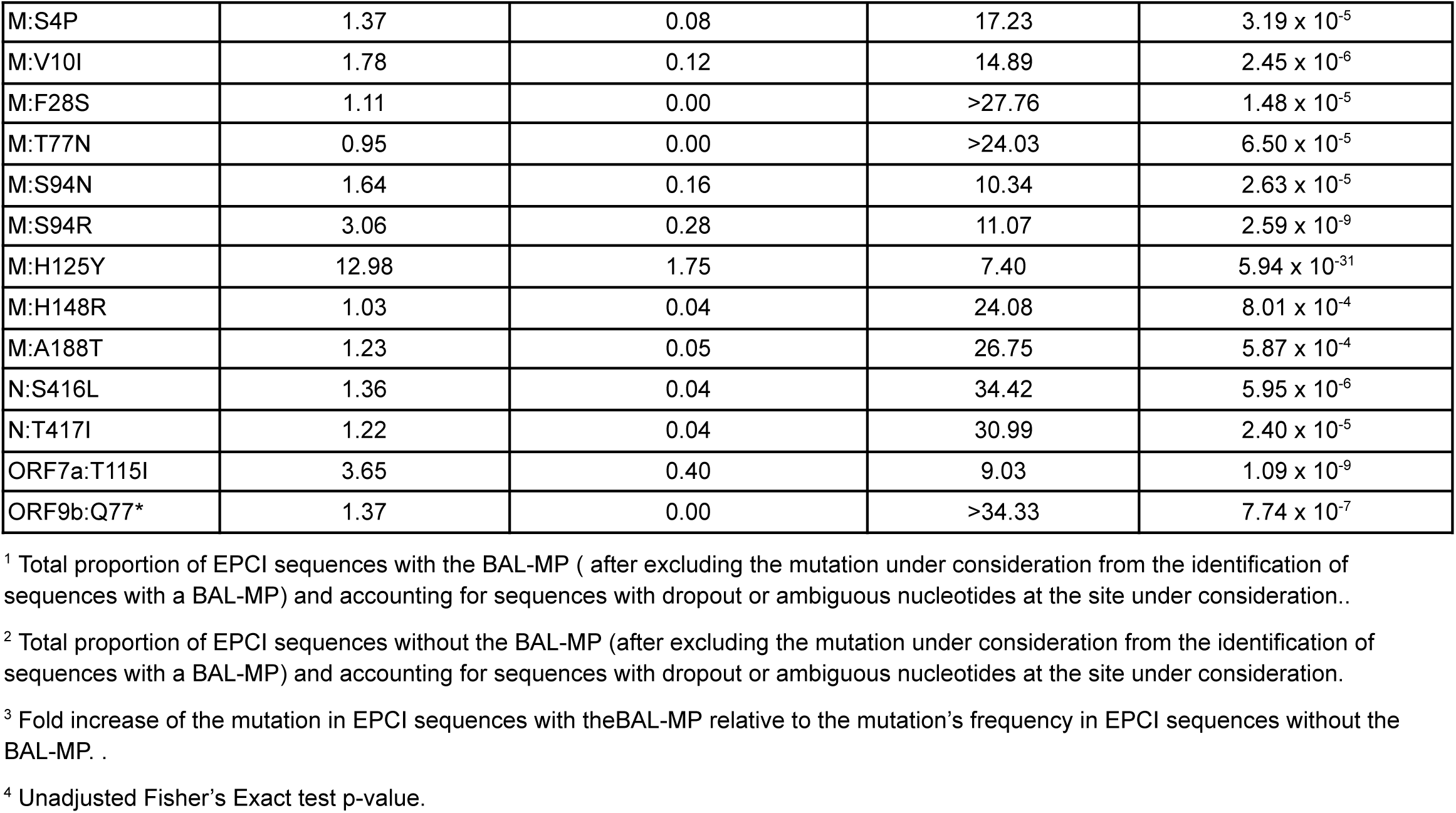
Full list of substitutions that occur significantly more frequently in EPCI sequences with a BAL-MP than they do in other EPCI sequences.

**Table 6:** Full list of substitutions that occur significantly more frequently in EPCI sequences with a 6-7a-MP than they do in other EPCI sequences.

| MP region # | Substitution | Adjusted % frequency in 6-7a EPCI sequences<sup>1</sup> | Adjusted % frequency in non-6-7a EPCI sequences<sup>2</sup> | Fold Increase<sup>3</sup> | Enrichment p-value<sup>4</sup> |
| --- | --- | --- | --- | --- | --- |
| 1(R1) | ORF1a:E1706D | 2.70 | 0.06 | 43.14 | 5.90 x 10<sup>-3</sup> |
|  | ORF1a:N1709S | 17.57 | 0.91 | 19.32 | 4.28 x 10<sup>-12</sup> |
|  | ORF1a:N1709T | 5.41 | 0.09 | 57.48 | 1.62 x 10<sup>-5</sup> |
|  | ORF1a:I1714A | 4.05 | 0.03 | 128.43 | 8.99 x 10<sup>-5</sup> |
|  | ORF1a:I1714V | 28.38 | 1.74 | 16.35 | 3.74 x 10<sup>-18</sup> |
|  | ORF1a:I1714T | 18.92 | 2.09 | 9.06 | 2.80 x 10<sup>-9</sup> |
|  | ORF1a:I1714S | 4.05 | 0.16 | 25.69 | 1.18 x 10<sup>-3</sup> |
| 2(R2) | ORF1a:M1769V | 6.00 | 0.16 | 37.68 | 6.07 x 10<sup>-7</sup> |
|  | ORF1a:M1769T | 4.00 | 0.00 | >125.6 | 1.71 x 10<sup>-6</sup> |
|  | ORF1a:M1769I | 5.00 | 0.06 | 78.50 | 1.01 x 10<sup>-6</sup> |
|  | ORF1a:T1773A | 1.90 | 0.03 | 59.71 | 6.11 x 10<sup>-3</sup> |
|  | ORF1a:T1773I | 8.57 | 0.26 | 33.59 | 1.10 x 10<sup>-9</sup> |
| 3(R3) | ORF1a:K1795Q | 27.59 | 9.01 | 3.06 | 1.64 x 10<sup>-6</sup> |
| 4(R4) | ORF1a:F2209L | 3.57 | 0.10 | 36.61 | 9.33 x 10<sup>-5</sup> |
|  | ORF1a:S2214L | 5.41 | 0.03 | 166.49 | 2.11 x 10<sup>-8</sup> |
|  | ORF1a:S2214A | 1.80 | 0.00 | >55.5 | 2.40 x 10<sup>-3</sup> |
|  | ORF1a:F2215L | 6.25 | 0.03 | 192.56 | 8.44 x 10<sup>-10</sup> |
| 5(R5) | ORF1a:V2324G | 2.60 | 0.00 | >80.94 | 1.15 x 10<sup>-3</sup> |
|  | ORF1a:A2325T | 9.59 | 0.29 | 33.21 | 4.77 x 10<sup>-8</sup> |
|  | ORF1a:A2325V | 32.88 | 1.03 | 32.02 | 3.50 x 10<sup>-26</sup> |
|  | ORF1a:F2328V | 12.00 | 0.45 | 26.71 | 1.69 x 10<sup>-9</sup> |
|  | ORF1a:F2328L | 8.00 | 0.32 | 24.93 | 1.83 x 10<sup>-6</sup> |
|  | ORF1a:L2329F | 12.99 | 0.45 | 28.91 | 1.09 x 10<sup>-10</sup> |
| 6(R6) | ORF1a:A2345V | 7.34 | 0.36 | 20.50 | 1.62 x 10<sup>-7</sup> |
|  | ORF1a:M2347L | 1.82 | 0.03 | 55.87 | 6.94 x 10<sup>-3</sup> |
|  | ORF1a:Q2348R | 1.82 | 0.03 | 55.95 | 6.92 x 10<sup>-3</sup> |
|  | ORF1a:L2349V | 3.64 | 0.00 | >112.0 | 2.68 x 10<sup>-6</sup> |
|  | ORF1a:Y2353H | 2.70 | 0.03 | 83.76 | 3.14 x 10<sup>-4</sup> |
|  | ORF1a:Y2353C | 2.70 | 0.10 | 27.92 | 1.49 x 10<sup>-3</sup> |
| 7 | ORF7a:E22D | 12.62 | 1.56 | 8.07 | 7.33 x 10<sup>-8</sup> |
| 8 | ORF7a:T39I | 37.23 | 5.69 | 6.55 | 1.69 x 10<sup>-18</sup> |
|  | ORF7a:N43K | 4.26 | 0.16 | 26.56 | 1.54 x 10<sup>-4</sup> |

<sup>1</sup> Total proportion of EPCI sequences with the 6-7a-MP (after excluding the mutation under consideration from the identification of sequences with an 6-7a-MP) and accounting for sequences with dropout or ambiguous nucleotides at the site under consideration.
<sup>2</sup> Total proportion of EPCI sequences without the 6-7a-MP (after excluding the mutation under consideration from the identification of sequences with an 6-7a-MP) and accounting for sequences with dropout or ambiguous nucleotides at the site under consideration.
<sup>3</sup> Fold increase of the mutation in EPCI sequences with the 6-7a-MP relative to the mutation's frequency in EPCI sequences without the 6-7a-MP.
<sup>4</sup> Unadjusted Fisher's Exact test p-value.

**Table 7:** Full list of sites where substitutions occur significantly more frequently in EPCI sequences with a 6-7a-MP than they do in other EPCI sequences.

| MP region # | Substitution position | Adjusted % frequency in 6-7a EPCI sequences<sup>1</sup> | Adjusted % frequency in non-6-7a EPCI sequences<sup>2</sup> | Fold Increase<sup>3</sup> | Enrichment p-value<sup>4</sup> |
| --- | --- | --- | --- | --- | --- |
| 1(R1) | ORF1a:1706 | 2.3 | 0.06 | 36.76 | $8.04 \times 10^{-3}$ |
| | ORF1a:1709 | 34.1 | 1.64 | 20.80 | $5.75 \times 10^{-28}$ |
| | ORF1a:1714 | 59.8 | 3.77 | 15.86 | $1.05 \times 10^{-26}$ |
| | ORF1a:1715 | 5.7 | 0.25 | 22.83 | $2.61 \times 10^{-5}$ |
| 2(R2) | ORF1a:1769 | 12.1 | 0.19 | 63.94 | $2.14 \times 10^{-18}$ |
| | ORF1a:1770 | 1.5 | 0.03 | 47.62 | $9.42 \times 10^{-3}$ |
| | ORF1a:1771 | 3.0 | 0.16 | 19.05 | $5.43 \times 10^{-4}$ |
| | ORF1a:1773 | 12.9 | 0.19 | 67.94 | $1.02 \times 10^{-19}$ |
| 3(R4) | ORF1a:2209 | 5.1 | 0.03 | 159.92 | $3.03 \times 10^{-9}$ |
| | ORF1a:2214 | 5.9 | 0.03 | 183.00 | $1.35 \times 10^{-10}$ |
| | ORF1a:2215 | 5.9 | 0.06 | 91.56 | $6.46 \times 10^{-10}$ |
| 4(R5) | ORF1a:2324 | 4.2 | 0.13 | 33.16 | $8.83 \times 10^{-5}$ |
| | ORF1a:2325 | 46.3 | 0.89 | 52.06 | $2.90 \times 10^{-23}$ |
| | ORF1a:2328 | 18.8 | 0.83 | 22.69 | $6.56 \times 10^{-17}$ |
| | ORF1a:2329 | 15.8 | 0.44 | 35.55 | $3.54 \times 10^{-16}$ |
| 5(R6) | ORF1a:2345 | 6.6 | 0.36 | 18.63 | $7.21 \times 10^{-8}$ |
| | ORF1a:2348 | 2.2 | 0.06 | 34.22 | $1.36 \times 10^{-3}$ |
| | ORF1a:2349 | 2.9 | 0.00 | >91.35 | $5.93 \times 10^{-6}$ |
| | ORF1a:2351 | 3.7 | 0.00 | >114.71 | $2.37 \times 10^{-7}$ |
| | ORF1a:2353 | 8.0 | 0.06 | 125.46 | $7.01 \times 10^{-14}$ |
| 6 | ORF7a:22 | 12.3 | 1.46 | 8.46 | $1.64 \times 10^{-9}$ |
| 7 | ORF7a:39 | 32.5 | 5.73 | 5.67 | $2.76 \times 10^{-18}$ |
| | ORF7a:43 | 5.7 | 0.76 | 7.46 | $2.22 \times 10^{-4}$ |

<sup>1</sup> Total proportion of EPCI sequences with the 6-7a-MP (after excluding the mutation under consideration from the identification of sequences with an 6-7a-MP) and accounting for sequences with dropout or ambiguous nucleotides at the site under consideration.
<sup>2</sup> Total proportion of EPCI sequences without the 6-7a-MP (after excluding the mutation under consideration from the identification of sequences with an 6-7a-MP) and accounting for sequences with dropout or ambiguous nucleotides at the site under consideration.
<sup>3</sup> Fold increase of the mutation in EPCI sequences with the 6-7a-MP relative to the mutation's frequency in EPCI sequences without the 6-7a-MP.
<sup>4</sup> Unadjusted Fisher's Exact test p-value.

**Table 8:** Full list of substitutions that occur significantly more frequently in EPCI sequences with a Rd3N-MP than they do in other EPCI sequences.

| MP region # | Substitution | Adjusted frequency in Rd3N EPCI sequences<sup>1</sup> | Adjusted frequency in non-Rd3N EPCI sequences<sup>2</sup> | Fold Increase<sup>3</sup> | Enrichment p-value<sup>4</sup> |
| --- | --- | --- | --- | --- | --- |
| 1 | ORF1a:T1542I | 10.87 | 2.12 | 5.1 | 1.74 x 10<sup>-6</sup> |
|  | ORF1a:T1543I | 23.36 | 3.33 | 7.0 | 1.76 x 10<sup>-16</sup> |
| 2 | ORF1a:R1566M | 1.28 | 0.00 | >40.49 | 4.40 x 10<sup>-3</sup> |
|  | ORF1a:T1567I | 4.49 | 0.57 | 7.9 | 2.10 x 10<sup>-4</sup> |
|  | ORF1a:V1570M | 1.92 | 0.09 | 20.2 | 3.69 x 10<sup>-3</sup> |
|  | ORF1a:T1572I | 4.49 | 0.19 | 23.6 | 1.21 x 10<sup>-6</sup> |
| 3 | ORF1a:A4396V | 16.05 | 1.06 | 15.2 | 6.41 x 10<sup>-11</sup> |
|  | ORF1a:A4396T | 7.41 | 0.19 | 39.7 | 3.01 x 10<sup>-7</sup> |
|  | ORF1a:S4398P | 3.61 | 0.22 | 16.6 | 3.25 x 10<sup>-3</sup> |
|  | ORF1a:S4398L | 55.42 | 4.04 | 13.7 | 1.45 x 10<sup>-38</sup> |
| 4 | ORF1b:L820F | 11.82 | 0.59 | 20.0 | 1.15 x 10<sup>-11</sup> |
|  | ORF1b:P821S | 10.00 | 1.09 | 9.2 | 3.20 x 10<sup>-7</sup> |
|  | ORF1b:Y822H | 8.18 | 0.62 | 13.1 | 3.96 x 10<sup>-7</sup> |
|  | ORF1b:D824E | 8.33 | 0.34 | 24.3 | 7.25 x 10<sup>-9</sup> |
|  | ORF1b:D824N | 3.70 | 0.34 | 10.8 | 2.19 x 10<sup>-3</sup> |
|  | ORF1b:S826A | 1.82 | 0.00 | >58.38 | 2.17 x 10<sup>-3</sup> |
| 5 | N:V270I | 2.19 | 0.09 | 23.3 | 2.50 x 10<sup>-3</sup> |
|  | N:V270L | 5.84 | 0.41 | 14.3 | 1.77 x 10<sup>-6</sup> |
|  | N:T271I | 20.29 | 2.36 | 8.6 | 2.80 x 10<sup>-16</sup> |

<sup>1</sup> Total proportion of EPCI sequences with the Rd3N-MP (after excluding the mutation under consideration from the identification of sequences with an Rd3N-MP) and accounting for sequences with dropout or ambiguous nucleotides at the site under consideration.
<sup>2</sup> Total proportion of EPCI sequences without the Rd3N-MP (after excluding the mutation under consideration from the identification of sequences with an Rd3N-MP) and accounting for sequences with dropout or ambiguous nucleotides at the site under consideration.
<sup>3</sup> Fold increase of the mutation in EPCI sequences with the Rd3N-MP relative to the mutation's frequency in EPCI sequences without the Rd3N-MP.
<sup>4</sup> Unadjusted Fisher's Exact test p-value.

**Table 9:** Full list of sites where substitutions occur significantly more frequently in EPCI sequences with a Rd3N-MP than they do in other EPCI sequences.

| MP region # | Substitution Position | Adjusted frequency in Rd3N EPCI sequences<sup>1</sup> | Adjusted frequency in non-Rd3N EPCI sequences<sup>2</sup> | Fold Increase<sup>3</sup> | Enrichment p-value<sup>4</sup> |
| --- | --- | --- | --- | --- | --- |
| 1 | ORF1a:1542 | 9.87 | 2.13 | 4.64 | 6.06 x 10<sup>-6</sup> |
|  | ORF1a:1543 | 22.37 | 3.52 | 6.35 | 4.19 x 10<sup>-16</sup> |
| 2 | ORF1a:1566 | 3.61 | 0.25 | 14.22 | 6.20 x 10<sup>-5</sup> |
|  | ORF1a:1567 | 4.82 | 0.64 | 7.59 | 8.81 x 10<sup>-5</sup> |
|  | ORF1a:1570 | 4.82 | 0.19 | 25.29 | 1.56 x 10<sup>-7</sup> |
|  | ORF1a:1572 | 6.63 | 0.35 | 18.96 | 3.14 x 10<sup>-9</sup> |
| 3 | ORF1a:4396 | 29.00 | 1.22 | 23.77 | 1.41 x 10<sup>-27</sup> |
|  | ORF1a:4398 | 58.00 | 4.69 | 12.36 | 4.90 x 10<sup>-46</sup> |
| 4 | ORF1b:820 | 10.29 | 0.56 | 18.23 | 9.51 x 10<sup>-12</sup> |
|  | ORF1b:821 | 11.76 | 0.94 | 12.50 | 1.79 x 10<sup>-11</sup> |
|  | ORF1b:822 | 7.35 | 0.69 | 10.65 | 5.67 x 10<sup>-7</sup> |
|  | ORF1b:824 | 14.81 | 0.56 | 26.23 | 1.33 x 10<sup>-18</sup> |
|  | ORF1b:826 | 1.47 | 0.00 | >46.84 | 3.33 x 10<sup>-3</sup> |
| 5 | N:270 | 7.50 | 0.51 | 14.83 | 3.36 x 10<sup>-9</sup> |
|  | N:271 | 20.89 | 2.34 | 8.92 | 2.44 x 10<sup>-18</sup> |

<sup>1</sup> Total proportion of EPCI sequences with the Rd3N-MP (after excluding the mutation under consideration from the identification of sequences with an Rd3N-MP) and accounting for sequences with dropout or ambiguous nucleotides at the site under consideration.
<sup>2</sup> Total proportion of EPCI sequences without the Rd3N-MP (after excluding the mutation under consideration from the identification of sequences with an Rd3N-MP) and accounting for sequences with dropout or ambiguous nucleotides at the site under consideration.
<sup>3</sup> Fold increase of the mutation in EPCI sequences with the Rd3N-MP relative to the mutation's frequency in EPCI sequences without the Rd3N-MP.
<sup>4</sup> Unadjusted Fisher's Exact test p-value.

**Table 10:** Full list of substitutions that occur significantly more frequently in EPCI sequences with a 2Macs-MP than they do in other EPCI sequences.

| MP region # | Substitution | Adjusted % frequency in 2Macs EPCI sequences<sup>1</sup> | Adjusted % frequency in non-2Macs EPCI sequences<sup>2</sup> | Fold Increase<sup>3</sup> | Enrichment p-value<sup>4</sup> |
| --- | --- | --- | --- | --- | --- |
| 1 | ORF1a:R232L | 1.09 | 0.03 | 33.3 | 4.19 x 10<sup>-3</sup> |
|  | ORF1a:E233K | 1.92 | 0.00 | >58.83 | 5.69 x 10<sup>-6</sup> |
|  | ORF1a:E233D | 1.15 | 0.00 | >35.3 | 9.51 x 10<sup>-4</sup> |
|  | ORF1a:E233A | 1.54 | 0.03 | 47.1 | 3.46 x 10<sup>-4</sup> |
|  | ORF1a:E233G | 5.00 | 0.00 | >152.95 | 6.32 x 10<sup>-15</sup> |
| 2 | ORF1a:S1272G | 3.65 | 0.36 | 10.3 | 2.61 x 10<sup>-5</sup> |
| | ORF1a:D1273G | 12.15 | 0.68 | 17.9 | $2.15 \times 10^{-19}$ |
| | ORF1a:D1273N | 3.27 | 0.23 | 14.4 | $2.02 \times 10^{-5}$ |
| | ORF1a:I1274M | 0.92 | 0.00 | >28.28 | $8.69 \times 10^{-3}$ |
| | ORF1a:I1276T | 4.13 | 0.62 | 6.7 | $9.22 \times 10^{-5}$ |
| 3 | ORF1a:S1361P | 6.35 | 0.50 | 12.6 | $3.22 \times 10^{-6}$ |
| | ORF1a:Q1365R | 4.92 | 0.32 | 15.6 | $2.69 \times 10^{-5}$ |
| | ORF1a:Q1365P | 5.74 | 0.16 | 36.4 | $1.09 \times 10^{-7}$ |
| | ORF1a:I1367L | 8.73 | 1.20 | 7.3 | $2.82 \times 10^{-6}$ |
| | ORF1a:L1368I | 6.35 | 0.44 | 14.4 | $1.50 \times 10^{-6}$ |
| | ORF1a:G1369R | 8.87 | 0.41 | 21.6 | $4.69 \times 10^{-10}$ |
| | ORF1a:S1372C | 5.00 | 0.32 | 15.8 | $2.52 \times 10^{-5}$ |
| | ORF1a:S1372A | 17.50 | 1.68 | 10.4 | $7.12 \times 10^{-14}$ |

<sup>1</sup> Total proportion of EPCI sequences with the 2Macs-MP (after excluding the mutation under consideration from the identification of sequences with an 2Macs-MP) and accounting for sequences with dropout or ambiguous nucleotides at the site under consideration.
<sup>2</sup> Total proportion of EPCI sequences without the 2Macs-MP (after excluding the mutation under consideration from the identification of sequences with an 2Macs-MP) and accounting for sequences with dropout or ambiguous nucleotides at the site under consideration.
<sup>3</sup> Fold increase of the mutation in EPCI sequences with the 2Macs-MP relative to the mutation's frequency in EPCI sequences without the 2Macs-MP.
<sup>4</sup> Unadjusted Fisher's Exact test p-value.

**Table 11:** Full list of substitutions that occur significantly more frequently in EPCI sequences with a 1Macs-MP than they do in other EPCI sequences.

| MP region # | Substitution | Adjusted % frequency in 1Macs EPCI sequences<sup>1</sup> | Adjusted % frequency in non-1Macs EPCI sequences<sup>2</sup> | Fold Increase<sup>3</sup> | Enrichment p-value<sup>4</sup> |
| --- | --- | --- | --- | --- | --- |
| 1 | ORF1a:H110Y | 29.41 | 0.63 | 46.4 | 2.89 x 10<sup>-29</sup> |
|  | ORF1a:I114T | 25.58 | 0.79 | 32.3 | 2.19 x 10<sup>-23</sup> |
| 2 | ORF1a:K958R | 1.87 | 0.03 | 60.1 | 6.04 x 10<sup>-3</sup> |
|  | ORF1a:P959L | 4.76 | 0.16 | 30.6 | 1.28 x 10<sup>-5</sup> |
|  | ORF1a:P959S | 7.62 | 0.41 | 18.8 | 2.25 x 10<sup>-7</sup> |
| 3 | ORF1a:P1220L | 6.82 | 0.29 | 23.8 | 2.81 x 10<sup>-6</sup> |
|  | ORF1a:P1220S | 13.64 | 0.51 | 26.8 | 3.29 x 10<sup>-12</sup> |
|  | ORF1a:P1220H | 2.27 | 0.03 | 71.5 | 4.23 x 10<sup>-3</sup> |
|  | ORF1a:S1221P | 2.20 | 0.00 | >69.05 | 1.57 x 10<sup>-3</sup> |
|  | ORF1a:E1223K | 3.16 | 0.06 | 49.6 | 4.69 x 10<sup>-4</sup> |
|  | ORF1a:Q1224R | 2.11 | 0.06 | 33.1 | 9.85 x 10<sup>-3</sup> |
| 4 | ORF1a:E1394D | 4.76 | 0.07 | 70.4 | 1.71 x 10<sup>-6</sup> |
|  | ORF1a:T1395N | 2.15 | 0.03 | 63.8 | 5.37 x 10<sup>-3</sup> |
|  | ORF1a:T1395I | 25.81 | 0.27 | 95.8 | 2.83 x 10<sup>-31</sup> |
|  | ORF1a:T1395A | 2.15 | 0.00 | >63.85 | 1.83 x 10<sup>-3</sup> |
|  | ORF1a:A1397V | 12.38 | 0.14 | 91.5 | 1.84 x 10<sup>-16</sup> |

<sup>1</sup> Total proportion of EPCI sequences with the 1Macs-MP (after excluding the mutation under consideration from the identification of sequences with an 1Macs-MP) and accounting for sequences with dropout or ambiguous nucleotides at the site under consideration.
<sup>2</sup> Total proportion of EPCI sequences without the 1Macs-MP (after excluding the mutation under consideration from the identification of sequences with an 1Macs-MP) and accounting for sequences with dropout or ambiguous nucleotides at the site under consideration.
<sup>3</sup> Fold increase of the mutation in EPCI sequences with the 1Macs-MP relative to the mutation's frequency in EPCI sequences without the 1Macs-MP.
<sup>4</sup> Unadjusted Fisher's Exact test p-value.

**Table 12:** Full list of substitutions that occur significantly more frequently in EPCI sequences with a Y1-6-MP than they do in other EPCI sequences.

| MP region # | Substitution | Adjusted % frequency in Y1-6 EPCI sequences<sup>1</sup> | Adjusted % frequency in non-Y1-6 EPCI sequences<sup>2</sup> | Fold Increase<sup>3</sup> | Enrichment p-value<sup>4</sup> |
| --- | --- | --- | --- | --- | --- |
| 1 | ORF1a:S2466G | 1.63 | 0.03 | 50.7 | $8.36 \times 10^{-3}$ |
| | ORF1a:D2467E | 3.23 | 0.03 | 100.6 | $1.98 \times 10^{-5}$ |
| | ORF1a:E2468D | 6.50 | 0.06 | 101.4 | $2.90 \times 10^{-10}$ |
| | ORF1a:V2469I | 2.42 | 0.00 | >75.39 | $1.10 \times 10^{-4}$ |
| | ORF1a:A2470S | 4.03 | 0.10 | 41.8 | $7.79 \times 10^{-6}$ |
| | ORF1a:R2471G | 5.93 | 0.10 | 61.7 | $1.58 \times 10^{-8}$ |
| | ORF1a:R2471I | 1.69 | 0.00 | >52.85 | $2.64 \times 10^{-3}$ |
| | ORF1a:R2471S | 8.47 | 0.03 | 264.2 | $6.07 \times 10^{-14}$ |
| | ORF1a:R2471K | 4.24 | 0.06 | 66.1 | $2.35 \times 10^{-6}$ |
| | ORF1a:D2472E | 6.45 | 0.23 | 28.6 | $3.86 \times 10^{-8}$ |
| | ORF1a:L2475I | 11.29 | 0.03 | 351.6 | $2.03 \times 10^{-19}$ |
| 2 | ORF1a:R3802C | 30.95 | 0.63 | 49.4 | $5.04 \times 10^{-31}$ |
| | ORF1a:R3802H | 14.29 | 0.47 | 30.4 | $9.33 \times 10^{-13}$ |
| | ORF1a:R3802S | 2.38 | 0.06 | 38.0 | $7.55 \times 10^{-3}$ |
| | ORF1a:R3802L | 2.38 | 0.00 | >75.93 | $1.30 \times 10^{-3}$ |
| | ORF1a:Y3803H | 10.34 | 0.16 | 65.9 | $1.57 \times 10^{-11}$ |
| | ORF1a:L3808F | 18.39 | 0.53 | 34.5 | $2.43 \times 10^{-17}$ |

<sup>1</sup> Total proportion of EPCI sequences with the Y1-6-MP (after excluding the mutation under consideration from the identification of sequences with an Y1-6-MP) and accounting for sequences with dropout or ambiguous nucleotides at the site under consideration.
<sup>2</sup> Total proportion of EPCI sequences without the Y1-6-MP (after excluding the mutation under consideration from the identification of sequences with an Y1-6-MP) and accounting for sequences with dropout or ambiguous nucleotides at the site under consideration.
<sup>3</sup> Fold increase of the mutation in EPCI sequences with the Y1-6-MP relative to the mutation's frequency in EPCI sequences without the Y1-6-MP.
<sup>4</sup> Unadjusted Fisher's Exact test p-value.

### Noteworthy BAL MP spike mutations

All mutations that eliminate the N657 glycan, such as N657K and S659P, are overrepresented in BAL MP sequences. Compared to ancestral SARS-CoV-2, in Omicron the N657 glycan is less occupied and less complex, possibly due to the nearby H655Y mutation^92^. A mutation eliminating the N657 glycan has been found to reduce infectivity^93^. Another study found S659 to be the major site of O-glycosylation in spike, and that its removal reduced spike stability and packaging efficiency^94^. As O-glycosylation has been found to occur immediately after N-linked glycans^95^, mutation at either N657 or S659 may eliminate the posited S659 O-linked glycan as well as the N657 glycan. How the loss of this glycan might be advantageous in the environment of the deep lung is not clear.

### Noteworthy BAL MP envelope mutations

E is a minor component of virions but is required, together with M, for the production of virus-like particles (VLPs)^96–97^. E oligomerizes to form pentamers in membranes, where it forms an ion channel (or viroporin). In contrast to other enveloped viruses, betacoronaviruses exit cells by the unconventional lysosomal pathway^98–100^. E is transported from the endoplasmic reticulum (ER) to the trans-Golgi compartment, from which a pool of E interacts with the adapter protein complex, AP-1, and is transported to lysosomes^101^, where its ion-channel activity raises lysosomal pH, creating a hospitable environment for exiting virions^101–102^.

Mutations that abolish E’s ion-channel activity severely inhibit coronaviral release from cells^103–104^. In mice and in cell culture, recombinant SARS-CoV viruses with ion-channel-inactivating mutants rapidly acquired compensatory mutations that restored viroporin activity and pathogenicity, a clear indication of its importance to viral replication.

With the partial exception of S68F, mutations in the E CTD are uncommon in BAL-MP sequences. Instead, the mutations in E are concentrated in E:5-42, which encompasses the transmembrane (TM) domain (roughly E:8-38), with E:T30I being by far the most common (in ∼40-45% of BAL-MP sequences). Notably, E:T30I has evolved when SARS-CoV-2 was repeatedly passaged in Calu3, Vero E6, Huh7, Caco2, and LLC-MK2 cell _lines_105-106.

In addition to governing E’s ion-channel activity, the TM domain contains an amphipathic helix crucial to oligomerization. In SARS-CoV (whose first 54 AA are identical to the SARS-CoV-2 Wuhan-hu-1 E sequence), the E:V25F mutation disrupts E oligomerization. When viruses with this mutation were passaged in cell culture and in mice, compensatory mutations rapidly arose^103^. Interestingly, one prominent reversion mutation was E:T30I, which outgrew the original virus by day two. In the E pentamer, E:30 faces toward the helix of an adjacent E protein, and other top BAL-MP E mutations, including ΔV14, S16N, A32V, and A36V, are similarly laterally oriented with respect to the E ion channel. Mutation of the N15 residue in E has been shown to eliminate or dramatically reduce its ion channel activity in coronavirus E proteins^102, 104, 297–298^. While mutations at N15 in BAL-MP EPCI sequences are less common than at other residues, seven different N15 substitutions (D, I, H, K, S, T, Y) occur in 16 EPCI sequences, and these mutations are almost entirely absent from all other SARS-CoV-2 sequences, suggesting that reduced ion-channel activity may be adaptive in the context of chronic infection of the deep lung.

Other top BAL-MP E mutations, such as V5A/F, G10S/C, and Y42C, are located just outside the transmembrane domain. All coronavirus E proteins contain 2-4 cysteine residues in the region immediately outside the C-terminal side of the transmembrane domain^107–108^. These cysteines are palmitoylated and have been shown to stabilize E^100, 108^. Mutations abolishing these cysteines resulted in impaired viral particle formation and reduced growth as well as markedly increased proteasomal degradation of E, possibly because palmitoylation regulates membrane association and E oligomerization^108–109^. SARS-CoV-2 E has cysteines at 40, 43, and 44. The addition of a fourth cysteine via a Y42C substitution may increase palmitoylation, which could conceivably increase the efficiency with which E oligomerizes and/or associates with membranes, though direct evidence of this is lacking.

Another intriguing explanation for the extreme density of E mutations in chronic-infection BAL-MP sequences is suggested by a recent study comparing individuals suffering from long Covid, also called Postacute Sequelae of COVID (PASC), to healthy convalescents (CP)^110^. The PASC group had much lower spike-specific antibody titers than the CP group, yet had higher levels of anti-E antibodies that persisted for at least 6 months post infection.

PASC patients displayed numerous other signs of persistent antigen exposure, including persistently elevated mucosa-associated invariant T cell (MAIT) and circulating T follicular helper cell (cTFH) counts. Furthermore, the PASC group displayed elevated IgG_1_ and IgG_3_ and depressed IgG_4_ levels compared to CP^110^. The low spike RBD mutation rate and extraordinarily high number of E mutations in BAL-MP sequences are precisely what one might expect to see in an environment with low anti-S and high anti-E antibody levels—as seen in these PASC patients.

### The BAL-MP may represent specialization for infecting alveolar macrophages

It has been shown that SARS-CoV-2 productively infects alveolar macrophages (AMs) *in vivo* and *in vitro*^111–117^, early in infection. A number of lines of evidence suggest that the BAL MP may represent adaptation to AM infection. One longitudinal study in which macaques were infected with wild-type or Omicron BA.1/BA.2 viruses exhibited striking parallels to chronic-infection case studies involving BAL MP sequences^111^. The infected macaques appeared to recover fully and both NP and endotracheal samples were PCR-negative from 21 d.p.i. onward. However, the infected macaques exhibited increased levels of inflammatory cytokines compared to healthy controls (HC) at 221 d.p.i. (similar to what is seen in human PASC) and distinct alterations in AMs, including elevated expression of IL-10 and MHC-E. Upon examination, 17/25 macaques harbored SARS-CoV-2 RNA-positive AMs, and 20/25 had spike-protein-positive AMs. All AMs with high levels of viral RNA replicated productively when cultured in BAL-fluid (BALF) macrophages, evidenced by increased dsRNA and NSP3 protein levels, as did 3/8 BALF AM samples in which no viral RNA could be detected.

Evidence that similarly prolonged LRT infections may often go undetected in humans comes from dozens of case studies documenting respiratory symptoms and radiological evidence of infection (e.g. lung infiltrates and ground-glass lung opacities) in patients whose NP swabs repeatedly test negative by RT-PCR but whose BAL samples test positive^145–166, 288^. In some cases, both NP and BAL samples were negative in patients later confirmed to have suffered prolonged infection, either through autopsy^160^ or by subsequent positive tests and sequencing analysis, as in one remarkable case study documenting an infection that lasted over a year^10^.

In that case, the patient was infected and hospitalized in July 2022, and sequencing revealed the variant to be BF.7. Despite ongoing fever, cough, and shortness of breath in subsequent months, the patient repeatedly tested negative via both NP and BAL. However, in May 2023, simultaneous BAL, NP, and endotracheal-aspirate samples were taken and tested positive. Sequencing revealed distinct URT and LRT intrahost variants, each descended from the original BF.7 strain. The BAL sequence from this patient contains at least five mutations from the BAL-MP at ≥50% prevalence (ORF1b:T2537I, S:P330S, E:L21F, E:T30I, and N:T417I), and possesses far more minority variants than the NP sequence. The NP sequence has two BAL- MP mutations not found in the BAL sequence (S:V551I and M:H125Y), suggesting that the deep lung may serve as a reservoir of viral genetic diversity from which the URT viral reservoir occasionally draws^10^.

Furthermore, in the macaque study viruses infecting AMs spread directly between AMs through syncytia formation and cell membrane protrusions connecting AMs, many of which contained viral proteins^111^. This type of cell-cell spread would prevent antibodies from binding to, and neutralizing, the spike protein, which could explain the marked absence of Ab-evasive spike mutations in sequences carrying the BAL MP. In addition, while AMs are robustly activated by the E protein through a TLR2- and TLR4-dependent manner, AMs do not react to spike protein^118–119^.

Infection of AMs has been shown to be ACE2-dependent. While many studies have failed to detect ACE2 expression in AMs through scRNA-seq, others have conclusively shown that AMs can express ACE2 and that SARS-CoV-2 infection of AMs is mediated by ACE2^112, 117^. While spike RBD mutations are rare in sequences carrying the BAL MP, a number of non-RBD spike mutations are strikingly common, and these mutations can have large effects on ACE2 affinity by modulating the frequency of RBD-up conformations. While the exact effect of a mutation on ACE2 affinity depends on the spike background in which it occurs, the effects of most mutations are similar across variants^121^. Using deep mutational scanning (DMS) results for XBB.1.5 and KP.3.1.1^120–121^, the top BAL-MP spike mutations, on average, confer a substantial increase in ACE2 affinity (see table S3). Since non-RBD spike mutations increase ACE2 affinity by favoring more open RBD conformations, they tend to increase vulnerability to neutralization, and this is also reflected in the DMS data for BAL-MP spike mutations.

Given the low level of ACE2 expression in AMs, increasing ACE2 affinity may be an important adaptation for AM cell entry. The ability of SARS-CoV-2 to spread directly from cell to cell in AMs would largely obviate the need to evade antibodies, tilting the balance of evolutionary pressure toward a more open RBD and increased ACE2 affinity.

In addition to ACE2, the cellular protease TMPRSS2 has been shown to be vital for efficient cell entry *in vivo* and in primary human lung cells ^290–292, 299^. For efficient cell entry, the SARS-CoV-2 spike must be cleaved at the S2’ site (S:815/816), which facilitates dramatic conformational changes that result in fusion with the cell membrane and delivery of the viral genome to the cytoplasm^293^. TMPRSS2 is the primary cellular protease responsible for S2’ cleavage, but AMs do not express significant levels of TMPRSS2 mRNA^294^. However, primary human nasal epithelial cells and lung epithelial cells release extracellular vesicles (EVs) containing ACE2 and TMPRSS2 which facilitate viral infection. EVs from lung organoids consisting of alveolar type II epithelial cells deliver ACE2 and TMPRSS2 to AMs, endothelial cells, and pericytes, facilitating SARS-CoV-2 infection of these cells, which do not otherwise display TMPRSS2 on their cell surface^294–295^. Although the exact processes by which ACE2 and TMPRSS2 are transferred and viral infection is enhanced are not yet clear, the discovery of EV-mediated delivery of ACE2 and TMPRSS2 to otherwise non-permissible cells provides a ready explanation for how SARS-CoV-2 proficiently infects AMs in a native pulmonary environment.

IFN-γ was shown to inhibit replication in BALF AMs, but BALF NK cells were deficient in IFN-γ expression^111^. SARS-CoV-2 has been shown to inhibit the cell-surface expression of MHC-I, a common viral tactic to evade detection by CTLs. The absence of MHC-I on the surface of an infected cell typically activates NK cells, prompting them to kill the cell with the “missing self marker” through the infusion of cytotoxic molecules. However, SARS-CoV-2 evades NK-cell cytotoxicity through multiple mechanisms, including downregulation of activating NK receptors and suppression of IFN expression^111, 122–127^.

In AMs, an additional NK cell–evasion tactic is increased cell-surface expression of MHC-E, a non-canonical class I MHC molecule. In primary human lung tissue and in a non-human primate (NHP) model, SARS-CoV-2 infection upregulated MHC-E expression, which inhibited NK cell cytotoxicity^111^. In NHPs, persistent AM infection was promoted by the countervailing effect of MHC-E on IFN-γ^111^. IFN-γ expression suppressed viral replication in cultured BALF AMs, but it also increased MHC-E expression, which inhibited NK cell degranulation, preventing the death of infected AMs. This “Catch-22” scenario could conceivably lead to cycles in which immune activation and IFN-γ reduce viral titers in the alveolar compartment but fail to clear the virus since the very immune reaction combating the virus also prevents its elimination by prolonging the life of the AMs it infects.

SARS-CoV-2’s ability to suppress IFN responses has been shown to be particularly thorough in lung tissue. While SARS-CoV-2 was found to evoke a robust IFN response in primary human nasal turbinate organoids, type I and type II IFN responses in 3D lung organoids and primary human lung tissue has been found to be low or absent^112, 116, 123, 128^, . In contrast, H1N1 influenza A virus provoked strong IFN responses in the same lung organoids, despite a lower viral load than SARS-CoV-2-infected organoids. Furthermore, treatment of infected AMs with IFN-α had no effect on viral replication^112^.

This offers a possible explanation for several rare and unusual features seen in some sequences carrying the BAL MP: mutations abolishing ORF6 expression, ORF9b expression, or NSP15 endonuclease activity. ORF6 and ORF9b are both potent suppressors of innate immune responses. ORF6 interacts with the nuclear pore complex to impede nuclear trafficking, stifling the nuclear import of host transcription factors^126, 129–131^. ORF9b binds to the mitochondrial membrane protein, TOM70, preventing its interaction with HSP90, an essential step in the activation of MAVS and the subsequent phosphorylation and nuclear import of IRF3, which ultimately culminates in the secretion of IFN-β. NSP15 cleaves dsRNA, which triggers host innate immune responses, including IFN production, and mutations that deactivate NSP15’s endonuclease activity have been shown to enhance host innate immune responses and decrease viral replication^132–133^.

One of the most remarkable instances of convergent evolution in the first two years of the COVID-19 pandemic was the independent acquisition by dozens of SARS-CoV-2 lineages—including nearly every variant of concern (VOC) and variant of interest (VOI)—of mutations downgrading the N Kozak sequence, which would be expected to increase the frequency of leaky scanning of N, which is the primary mode of ORF9b expression^283^. These same N-Kozak mutations are extremely common in EPCI sequences involving pre-VOC variants^283^. Increased ORF9b protein expression in major VOC has been demonstrated, including in the only two VOC/VOI which lacked a mutation in the N Kozak sequence: Beta and Gamma^124–126, 134^.

Some EPCI sequences are marked by ORF6/ORF9b stop codons, mutation of the ORF6/ORF9b start codon, frameshifting deletions in ORF6, or mutations at or adjacent to an NSP15 endonuclease active site residue (Table S5a). A majority of these sequences possess multiple additional BAL-MP mutations—2.40, 2.61, and 2.98 per sequence for ORF6, NSP15, and ORF9b loss-of-function mutations, respectively (compared to an average of 1.01 BAL-MP mutations per EPCI sequence) (Table S5b). The presence of these ORF6, ORF9b, and NSP15 mutations in BAL sequences—despite their rarity in EPCI sequences overall—is readily explained if these viruses primarily infect AMs and viral replication in AMs is unaffected by type-I IFN expression, as has been demonstrated previously^112^.

While the BAL-MP is normally not seen in circulating sequences, one designated Pango variant, B.1.616, possessed several major BAL MP mutations, including E:F20L, E:T30I, M:H125Y, ORF1b:T2537I, a frameshifting deletion in ORF6, and a five-amino acid extension of ORF7a^135^. There are 14 total EPCI sequences with a similar four-to-five AA extension of ORF7a. Remarkably, 13 of these 14 sequences display a distinct BAL-mutation signature, with multiple additional BAL-MP mutations (table S6). B.1.616 also has S:Q949R, which, on a XBB.1.5 or a KP.3.1.1 genomic background causes a very large increase in ACE2 affinity^120–121^.

Remarkably, ORF7a stop-codon mutations that are either identical to or very similar to the one found in B.1.616 appear in 20 EPCI sequences, nearly all with strong BAL-mutation signatures, and which extend the ORF7a tail by either four or five amino acids (usually WLNFH, RLNFH, or WNFH). The C-terminal ORF7a tail (at amino acid positions 117-121) encodes an ideal ER-retention (ERR) motif (KxKx[D/E]) that governs ORF7a intracellular trafficking and is critical for activating the NF-kB immune response^179–183^, which enhances SARS-CoV-2 replication^296^. In a mouse model in which ORF7a was added to the prototypical betacoronavirus MHV, mutations to the two lysine residues in the ORF7a tail attenuated disease, reduced viral loads *in vivo* and *in vitro*, and decreased levels of pro-inflammatory cytokines^183^, though surprisingly, no change in ORF7a localization was detected. Notably, the only other ORF7a BAL-MP mutation is T115I (Table 5), very close to the ORF7a ERR motif (at 117-121). Speculatively, a downregulated immune response resulting from ablation of the ERR could prove adaptive in the context of prolonged infection. However, the proviral functions of the ORF7a CTD tail are poorly understood, and the effects of the CTD-tail extensions seen in B.1.616 and many BAL-signature EPCI sequences are unknown.

Notably, B.1.616 is the only variant that was usually not detectable by NP-swab PCR tests, though it was readily detected in BAL samples, a clear sign of its deep-lung tropism. Furthermore, B.1.616 was among the most virulent variants known, lethal in 46% of detected cases (albeit in an elderly cohort, median age 81 years) and with a hazard ratio for poor outcomes of 4.0 relative to VOC (mostly Alpha)^135^. Examination of BAL samples from B.1.616-infected patients showed that infected and uninfected AMs appeared in the same samples, while an autopsy specimen revealed that lung regions with infected AMs were located adjacent to healthy, uninfected regions^116^. Such discrete areas of infection may provide for the evolution of multiple viral lineages within the lungs of a single patient. This may explain why, in a chronically infected patient from which NP and BAL samples were collected at the same time, the BAL sample showed far greater genetic diversity than the NP sample^10^.

The existence of multiple diverse intrahost variants in chronically infected hosts, possibly inhabiting different tissue compartments, is well documented^10, 19, 25, 28–29, 49, 137^. Adaptation to different bodily compartments may introduce additional diversity, as documented in cryptic wastewater variants involving long-term infections, many of which acquired mutations found in Omicron and other VOC months or years before they appeared in circulation^242–245^, suggesting that within-host evolution may occur in the gastrointestinal tract over many months with variants arising there later recolonizing to the respiratory tract^138^.

Within the respiratory tract, the simultaneous existence of very different SARS-CoV-2 variants in the lungs to those found in the nasal cavity has been shown in a chronically infected individual^10^. Experiments with influenza A virus and SARS-CoV-2 in mice indicate that even within the same tissue compartment, intermixing of different viral populations is restricted to a narrow boundary^70, 139^, which is consistent with semi-independent evolution of numerous intrahost populations during long-term infection.

Coronavirus replication involves the production of subgenomic RNA (sgRNA) molecules encoding all four structural proteins and numerous accessory proteins via template switching in which the viral RNA-dependent RNA polymerase (RdRP) disconnects from viral gRNA at a transcription regulating site (TRS) and reattaches at a leader TRS (TRS-L) site at the 5’ end of the genome, skipping over thousands of intervening nucleotides^140–141^.

Since this intragenomic recombination is an inherent aspect of their replication/transcription, coronaviruses readily recombine, as phylogenetic evidence makes clear^142–144^. Regular recombination at the boundaries of distinct intrahost viral populations during chronic infection presents a favorable environment for the production of novel genotypes unlikely to evolve during acute infection and circulation.

### The 6-7a Mutation Pattern

The majority of MPs in EPCI sequences do not appear to be related to T cell escape and are clearly linked, like the BAL-MP, to infection of a specific tissue compartment. Deciphering what adaptive advantage they provide is therefore necessarily speculative, but in some cases known interactions between the proteins or protein domains within which the mutations occur indicates that coevolution between functionally interacting protein regions is a plausible explanation for the MPs. In some cases, the coevolution implied by the MPs might be evidence of undiscovered aspects of the SARS-CoV-2 life cycle.

One MP that seems likely to involve functional relationships between the genomic regions involved is the 6-7a-MP, so called because it involves six different NSP3 regions and several specific residues in ORF7a. The NSP3 domains involved are the PLpro domain: (ORF1a:1709-1714 hereafter referred to as R1, ORF1a:1769-1773 referred to as R2, and ORF1a:1795, R3), the C-terminal subdomain of the betacoronavirus-specific marker (BMS) domain (ORF1a:2209-2215, R4), the ectodomain TM-like helix (ORF1a:2325-2329, R5), and the TM2 domain (ORF1a:2345-2353, R6)^277–278^, while the related ORF7a residues are ORF7a:22, 39, 43, and 59. None of these regions are known immunodominant T cell epitopes^71^, suggesting either that the different regions interact in some way during the viral life cycle, or that the mutations occurring within the regions are adaptive in the context of a particular internal tissue environment.

One of the 6-7a mutations, ORF1a:K1795Q, is known to increase PLpro’s ability to deubiquitinate K48-linked ubiquitin chains and is a reversion to the residue found in all other known sarbecoviruses (as discussed above), which suggests the linked mutations also have functional effects unrelated to the evasion of T cells. R2 includes ORF1a:M1769, which was shown in SARS-CoV to be a crucial residue for binding ubiquitin, with which it directly interacts^167–168^. This suggests the possibility that, like K1795Q, mutations in R2 may be modulating PLpro’s DUB and deISGylating activity, perhaps shifting its substrate preference or enhancing its modest ability to bind K63-linked ubiquitin chains.

Like most positive-sense RNA viruses, SARS-CoV-2 transforms host-cell organelle membranes into replication organelles in which genomic RNA (gRNA) and subgenomic RNA (sgRNA) transcription takes place^169^. In coronaviruses (CoVs), NSP3 and NSP4 manipulate endoplasmic reticulum (ER) membranes to form double-membraned vesicles (DMVs)^170–172^. The CoV replication complex, consisting of NSP7-NSP16, produces gRNA and sgRNA within the DMV, which shields double-stranded RNA (dsRNA) replication intermediates from detection by host-cell immune sensors such as MDA5 and RIG-I ^173–176^.

NSP3 and NSP4 form dodecameric, DMV-spanning pores through which viral RNA is extruded into the cytoplasm, where it can be translated by ribosomes or, in the case of gRNA, encapsidated by nucleocapsid for assembly into virions. NSP4 forms the inner portion of the pore and NSP3 the outer portion, while NSP3 and NSP4 ectodomains interact within the DMV intermembrane space^177–178^. The cytoplasmic side of the DMV pore is topped by a pronged crown formed by the first eight of NSP3’s 16-17 domains (Ubl1, Mac1, Mac2, Mac3, DPUP, Ubl2, PLpro, NAB).

The disordered betacoronavirus-specific domain (ßSM) links this crown region with the NSP3 ectodomain, which is followed by a transmembrane (TM) domain that leads back to the cytoplasmic side of the DMV, where the highly conserved Y1 and CoV-Y regions form the base of the DMV crown^177–178^.

While several of the 6-7a mutations are always to specific AA residues (ORF1a:K1795Q, ORF7a:E22D, ORF7a:T39I), others display great variety, both in position and residue. Among EPCI sequences, for example, although the bulk of ORF1a:N1709 mutations have been either N1709S or N1709D (62/78, 79.5%), 10 different AA substitutions have occurred at this site—D, E, G, H, K, I, R, S, T, and Y—five of which require two nucleotide substitutions. Similarly, six different AA mutations have occurred at ORF1a:I1714 in EPCI sequences (A, L, M, S, T, V), four at ORF1a:F2328 (C, L, S, V), four at ORF1a:L2329 (F, M, S, V), five at ORF1a:Y2353 (C, F, H, N, S), and five at ORF7a:N43 (I, K, S, T, and Y), all of which suggests some degree of diversifying evolution.

Structurally, within the most comprehensive and precise structure of the DMV pore available (PDB: 8YAX), the three PLpro regions are near enough to each other that a direct interaction is plausible, with the side chains of R1 and R2 15-30 Å apart and the R2 and R3 side chains 10-15 Å distant. Similarly, R5 and R6 side chains are within 12-20 Å of each other. R4 is disordered and is not represented in the DMV structure model, but the side chain of ORF1a:F2223, which is within eight AA of R4, is 55 and 47 Å apart from the nearest R5 and R6 side chains. The PLpro regions (R1-R3) are separated from R5 and R6 by more than 100 Å, making any direct interaction in the context of the final DMV pore structure implausible. However, the dynamics of DMV formation are largely uncharacterized, so it is conceivable that these regions either directly interact or act synergistically during DMV formation. Furthermore, although the dominant role of NSP3 is likely in the context of the formation of replication organelles and the DMV-pore structure, it may also carry out auxiliary functions in other cellular contexts, though the exact nature of such roles remains unknown.

The reason(s) that sites in ORF7a might be coevolving with sites in NSP3 is less evident. The function of ORF7a in the viral life cycle is uncertain. It is not essential to viral replication or transmission as numerous SARS-CoV-2 variants, including several of the last XBB* subvariants (including GE.1.2.1, GW.5.1.1, and FE.1.1) and BA.3.2, are entirely missing ORF7a. Among the many possible functions attributed to ORF7a are, among others, stimulation of NF-kB^179–183^, antagonism of type-I IFN responses^184–187^, downregulation of MHC-I surface expression^188–189^, inhibition of antiviral effects of BST2 (tetherin)^190–194^, disruption of the antiviral protein SERINC^187, 195–196^, and inhibition of autophagy^197–200^. However, most of these posited functions and activities were inferred based on experiments involving the overexpression of ORF7a, which could potentially have precipitated spurious phenomena not found in the context of normal infection.

Three specific ORF7a mutations—E22D, T39I, and F59I—are strongly associated with ORF1a:K1795Q in EPCI sequences. ORF7a residue 59 was shown, in the context of ORF7a overexpression, to govern the ability of various sarbecovirus ORF7a proteins to downregulate the surface expression of MHC-1 in 293T cells^189^. Unlike sarbecoviruses with a hydrophobic residue (F or I) at ORF7a:59, SARS-CoV and others with a polar residue (S or T) at ORF7a:59 showed no ability to inhibit MHC-I. Fascinatingly, of the eight other sarbecovirus ORF7a proteins tested, only Rf1 displayed a superior ability to reduce MHC-I cell-surface expression compared to SARS-CoV-2, and Rf1 was also the only virus tested with isoleucine (I) at residue 59. Perhaps relevant in this context is that in a comparison between ancestral and major early variants (WA1, Alpha (B.1.1.7), Beta (B.1.351), Gamma (P.1), Epsilon (B.1.427/9), and Iota (B.1.526)), Gamma, the only variant with the ORF1a:K1795Q mutation, was most effective at suppressing MHC-I surface expression, despite having the highest expression levels of MHC-I mRNA^201^. The association of these ORF7a mutations with ORF1a:K1795Q and the other 6-7a mutations may therefore represent adaptations to a host environment in which there is intense selection pressure to inhibit MHC-I antigen presentation and CD8 T cell responses.

Another possible connection between K1795Q and ORF7a mutations is modulation of the pro-inflammatory NF-kB pathway. The region responsible for ORF7a’s established ability to activate NF-kB is not known, though it does not depend on two key C-terminal lysine residues (K117 and K119) that are important to other ORF7a functions^183^. SARS-CoV PLpro’s ability to inhibit NF-kB is due to its ability to cleave K48-linked diubiquitin from ubiquitin chains. As K1795Q dramatically increases the SARS-CoV-2 PLpro’s DUB activity, and therefore presumably its ability to antagonize NF-kB, the ORF7a mutations that frequently accompany K1795Q in EPCI sequences may either moderate or facilitate this PLpro-driven NF-kB inhibition.

### The Rd3N Mutation Pattern

Another multi-protein MP seen in EPCI sequences is Rd3N (for <u>R</u>NA-<u>d</u>ependent RNA polymerase, NSP<u>3</u>, <u>n</u>ucleocapsid), and involves mutations in the DPUP domain of NSP3 (ORF1a:1542-1543), the NSP12 NTD (ORF1a:4396-4398), the NSP12 thumb domain (ORF1b:820-824), and the nucleocapsid CTD (N:270-271). None of these narrow regions have been identified as major T cell epitopes, and there is no known interaction between these protein domains in the coronavirus life cycle. All three proteins, however, are involved in replication and transcription.

Nucleocapsid, in addition to its role of encapsidating the viral genome for assembly in virions, plays numerous other roles during infection, including antagonizing stress granules^279–281^, evasion of cellular antiviral pattern-recognition receptors (PRRs)^280–282^, and regulating the production of viral gRNA and sgRNA^207–209, 220–222^. It consists of five major domains: an N-terminal disordered region (N1), a folded RNA-binding domain (N2), a disordered central region containing a serine-arginine (SR) rich segment (N3), a folded region important for dimerization (N4), and a C-terminal disordered region important for higher-order oligomerization and N-M interactions (N5). Phosphorylation of the central SR region of the N3 domain plays a vital role in governing N’s transition between its different roles during infection.

The naked gRNA of most positive-strand RNA viruses is capable of initiating full-blown infection, but the infectiousness of coronavirus gRNA is dramatically reduced—as much as 1000-fold—compared to that of intact virions^202–203^. However, the addition of N restores gRNA to full infectiousness. Exchanging the N3 domain of the model betacoronavirus MHV with those of SARS-CoV or Bovine coronavirus (BCoV) was shown to be lethal and severely deleterious, respectively, but in both cases revertant mutants in the SR region of the N3 domain

(N:176-206 in SARS-CoV-2)^203–205^, and/or the N-terminal Ubl1 region of NSP3 (ORF1a:819-929) largely restored infectivity: the first indication of a crucial NSP3-N interaction in coronaviruses, which was later characterized structurally^204–205^. In SARS-CoV-2 specifically, a leucine-rich helix from 219-230 and a polar strand from 243-255 fold around Ubl1, causing dramatic compaction of N^204^.

Like many positive-strand RNA viruses, coronaviruses manipulate host-cell membranes to form replication organelles. NSP3 and NSP4 of SARS-CoV-2 create DMVs from the ER while simultaneously forming dodecameric pores that connect DMVs to the cytoplasm. NSP3 forms the outer part of the pore, with its N-terminal domains composing a “crown” topped by prongs. Ubl1 is disordered and extends from the tips of DMV-pore prongs, and this is where the N-Ubl1 interaction takes place. The NSP3 mutations (ORF1a:T1542I, T1543I, T1567I, V1570M, T1572I) in the Rd3N-MP are also found in the DMV-pore crown but in the DPUP domain of NSP3 and a linker region connecting DPUP and Ubl2 (not to be confused with Ubl1), which is located below the prongs to which Ubl1 is connected (see PDB 8YAX^178^). The function of DPUP is not known, though it may modulate the RNA binding ability of the adjacent Mac3 domain^206^.

The vast majority of N mutations in the Rd3N-MP are V270L and T271I, which reside in the N-terminal part of the globular N4 domain (N:263-364) within a region called the CTD basic patch (CBP, N:247-279)^207^. N:270-271 lie outside the region known to directly interact with Ubl1. However, both the N2 and N4 domains of N have been shown to be required for N’s ability to stimulate replication^205, 208–209^, implying they play an indirect role in the N-NSP3 interaction. In fact, when each of the five individual MHV N domains was expressed individually in MHV-infected cells, only the CTD was recruited to DMVs^210^, likely through dimerizing or oligomerizing with the WT N from the infecting virus.

Furthermore, two recent studies have shown that ISGylation of lysine residues in the N4 and N5 domains impedes N oligomerization and reduces single-cycle SARS-CoV-2 replicon RNA production^211–212^ without interfering with N-Ubl1 binding^211^, proving that N oligomerization/dimerization is required for efficient SARS-CoV-2 transcription/replication, a function that could potentially be affected by N:270-271 mutations. ISGylation involves the attachment of ISG15, an interferon-stimulated gene (ISG) whose production is enhanced during viral infection.

In addition, advanced imaging of structural viral proteins during SARS-CoV-2 infection indicates double layers of N can form around DMVs^99^. As there are only eight to ten pores per DMV^172^, a double-layer of N surrounding DMVs would likely involve extensive oligomerization or possibly liquid-liquid phase separation (LLPS), where certain components (such as proteins and RNA) are concentrated in membraneless compartments. The ability of N to induce LLPS has been extensively documented and is regulated by phosphorylation of the SR region of the N3 domain^207, 213–218^.

The NSP12 mutations in the Rd3N-MP are at opposing ends of the protein, with ORF1a:4396-4398 located at the extreme N-terminal part of NSP12 (residues 4-6) and ORF1b:820-824 (NSP12:829-833) in the C-terminal thumb region^219^. While these two regions are distant from one another and therefore cannot directly interact, they do appear on the same face of NSP12 and might therefore both interact with a third molecule.

Although N has long been thought to regulate viral RNA production, it is unclear whether it enhances sgRNA or gRNA production (or perhaps each at different time points during infection)^220–222^. N has been shown to phase separate with replicase components when overexpressed^215^, but the relevance of this *in vivo* is unclear. The viral replicase complex, which includes, at minimum, one copy of NSP7 and NSP12 and two copies each of NSP8 and NSP13, is primarily found within DMVs during infection, and N would need to be enclosed in DMVs during their formation in order to access the replicase. As the sgRNA expressing N dominates sgRNA populations early in infection^99^, this conceivably could occur, but there is no direct evidence that N localizes inside DMVs. Since the Ubl1 domain of NSP3 is located on the outside of the DMVs, N bound to Ubl1 is unlikely to directly influence the viral replication machinery.

Detailed study of the localization of SARS-CoV-2 proteins during infection indicate that NSP4-NSP16 are all predominantly located within DMVs, close to the inner DMV membrane and at angular positions that correlate closely with DMV pores^176^. This likely results from uncleaved NSP4-12/NSP4-16 polyprotein positioning on the interior membrane of forming DMVs, with NSP5-driven polyprotein cleavage only occurring after DMV closure. Structural analysis and Alphafold-assisted modeling lend additional support to this model^174^, as does the detection of unaccounted for density in the interior of DMV pores^177^. Co-immunoprecipitation of NSP4, NSP7, NSP8, NSP9, NSP12, and NSP13 by NSP3 antibodies and of NSP3, NSP4, NSP7, NSP9, NSP12, and NSP13 by anti-NSP8 antibodies is also consistent with the existence of a DMV pore-replicase complex^173^.

Once the ORF1a/ORF1ab polyprotein is cleaved within the DMV interior, the individual NSPs would coalesce to form the full replicase complex and would have no way of accessing the cytoplasm. However, NSP6 and NSP15 carry out important proviral functions in the cytoplasm. NSP6 recruits lipid droplets and ER vesicles to the DMV exterior, providing lipid and membrane material necessary for DMV growth, while NSP15 cleaves viral dsRNA in the cytosol in order to prevent its detection by cellular pattern-recognition receptors like RIG-I and MDA5, which trigger IFN production and the establishment of an antiviral state. This implies that at least some polyprotein cleavage must occur in the cytoplasm.

While it is clear that the majority of viral RNA production occurs in DMVs, it seems certain that immediately after viral entry, some degree of replication takes place in the cytoplasm before a sufficient quantity of NSP3 and NSP4 (the minimal required proteins for DMV formation) have been produced. For example, in the JHMV strain of MHV, the cellular helicase DDX1 associates with phosphorylated N early in infection to preferentially stimulate the production of gRNA and long sgmRNA, and DDX1 knockdown seriously impedes JHMV replication^220^. DDX1 also facilitates the synthesis of long RNAs in the gammacoronavirus IBV, and preventing phosphorylation of SARS-CoV-2 N protein also results in a reduction in NSP3, spike, and membrane proteins, while N protein levels were unaffected^300^, all of which suggest this role of DDX1 may be universal in coronaviruses^220^.

Similarly, the cellular protein SND1 binds preferentially to the 5’ end of negative-strand SARS-CoV-2 viral RNA and is required for early viral RNA synthesis, especially sgRNA production^223^. Furthermore, SND1 knockout reduced sgRNA levels relative to gRNA early in infection but had no effect on the sgRNA/gRNA ratio late in infection^223^. Together with the lack of any known mechanism to transport SND1 or DDX1 into DMVs, these findings suggest a substantial degree of cytosolic RNA production takes place in coronavirus-infected cells.

Similarly, it seems possible that the NSP3 and N regions mutated in the Rd3N-MP could interact with the Rd3N NSP12 regions during this cytosolic RNA replication process, though any interaction within DMVs appears impossible.

### The 2Macs Mutation Pattern

An MP we call 2Macs involves mutations in NSP2 (ORF1a:232-233), the NSP3 Mac2 domain (ORF1a:1272-1273), and a linker between the NSP3 Mac2 and Mac3 domains (ORF1a:1361-1372; Figure 11).

**Figure 11:**
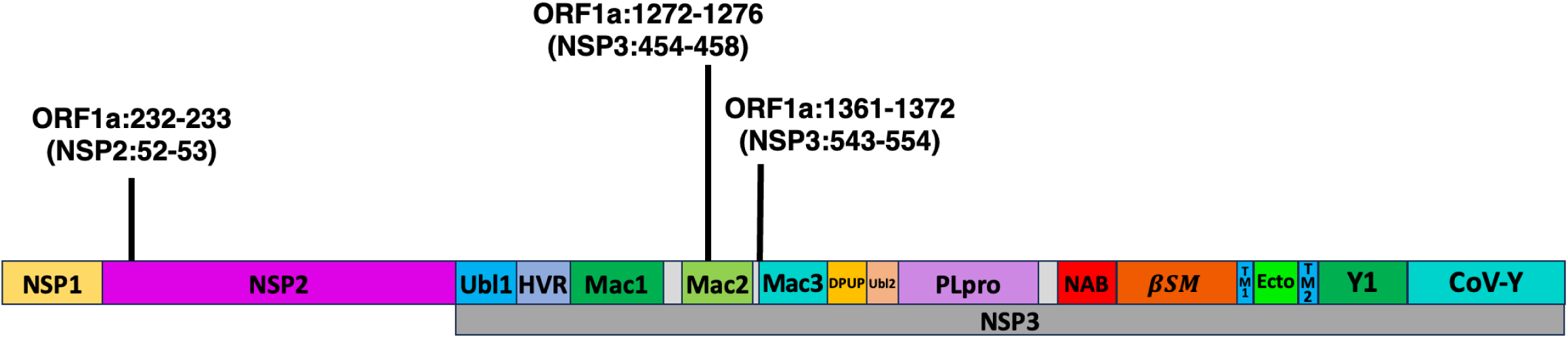
NSP1, NSP2, and NSP3 with the 2Macs MP-associated mutations labeled. Ubl1 = ubiquitin-like domain 1; HVR = hypervariable region (also called acidic domain); Mac1 = macrodomain 1; Mac2 = macrodomain 2; Mac3 = macrodomain 3; DPUP = Domain preceding Ubl2 and PLpro; Ubl2 = ubiquitin-like domain 2; PLpro = papain-like protease; NAB = nucleic acid-binding domain; βSM = betacoronavirus-specific marker domain; TM1 = transmembrane domain 1; Ecto = ectodomain; TM2 = transmembrane domain 2; Y1 = Nidovirus-conserved domain of unknown function; CoV-Y = Coronavirus-specific C-terminal domain

**Figure 12:**
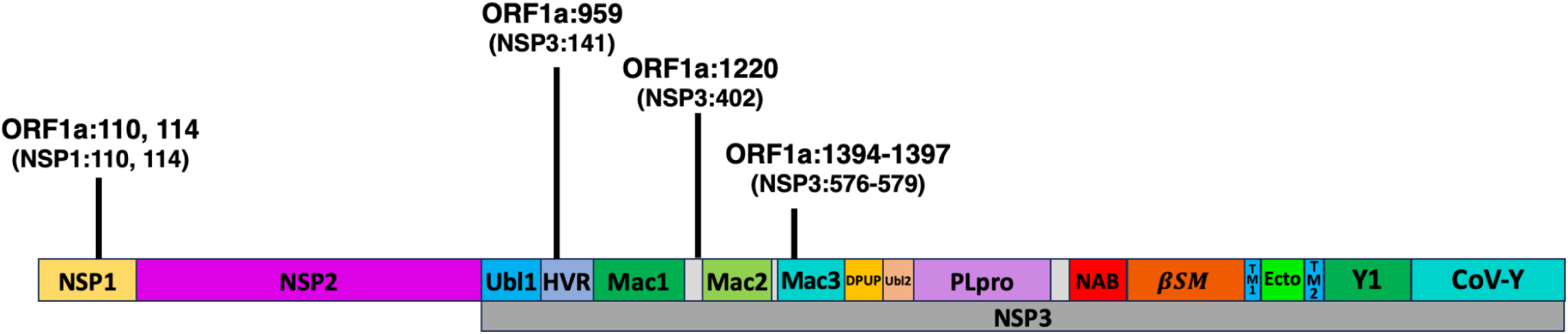
NSP1, NSP2, and NSP3 with the 1Macs MP-associated mutations labeled. Ubl1 = ubiquitin-like domain 1; HVR = hypervariable region (also called acidic domain); Mac1 = macrodomain 1; Mac2 = macrodomain 2; Mac3 = macrodomain 3; DPUP = Domain preceding Ubl2 and PLpro; Ubl2 = ubiquitin-like domain 2; PLpro = papain-like protease; NAB = nucleic acid-binding domain; βSM = betacoronavirus-specific marker domain; TM1 = transmembrane domain 1; Ecto = ectodomain; TM2 = transmembrane domain 2; Y1 = Nidovirus-conserved domain of unknown function; CoV-Y = Coronavirus-specific C-terminal domain

NSP2 is one of the most enigmatic SARS-CoV-2 proteins. At 638 amino acids, it is the fourth-largest SARS-CoV-2 protein but has no well established function. Although it is dispensable for replication in both SARS-CoV and MHV, its deletion substantially attenuates peak viral titers^224–225^. In MHV, NSP2 localizes to the outer membrane of DMVs^226–227^, and recent imaging of SARS-CoV-2 proteins during infection also found that NSP2 formed rings around DMVs and colocalized with NSP3^176^, a finding perhaps relevant to the 2Macs-MP. The same study found that late in infection, NSP2 relocated to Golgi bodies that had been fragmented and dispersed.

While many overexpression studies of NSP2 in SARS-CoV-2 have been performed, only recently was NSP2 function interrogated in the context of viral infection. Consistent with overexpression studies^228–230^, it was found that NSP2 associated with the host translation suppressor protein GIGYF2. While NSP2 associated with DMVs in the context of infection, when ectopically expressed, it diffused throughout the cytosol, suggesting it is recruited to DMVs directly or indirectly by another viral protein. Since the N-terminal half of NSP3 projects off the cytosolic side of the DMV membrane, it could plausibly be responsible for NSP2’s localization there early in infection.

Similarly, while endogenous GIGYF2 and its cofactor ZNF598 are distributed throughout the cytosol, in SARS-CoV-2 infection, they are recruited to the surface of DMVs by NSP2. Infection with an NSP2-deletion virus (ΔNSP2) failed to recruit GIGYF2 and ZNF598 to DMVs and led to a 60-fold reduction in viral RNA at 5 hpi, a replication defect that was repaired with ectopic expression of NSP2^231^. GIGYF2 was found to bind to and enhance the translation of M and ORF6 during infection but not when NSP2, M, and ORF6 were ectopically expressed^231^.

The roles of the Mac2 (ORF1a:1231-1358) and Mac3 (ORF1a:1369-1493) domains of NSP3, apart from their structural role in the DMV pore, are not known with certainty. Both bind to G-quadruplex-forming RNA^206, 238^, though what effect this might have, if any, during infection is not known. Deletion of Mac2 in a SARS-CoV replicon only moderately reduced replication but deletion of Mac3 destroyed all replication, as did the mutation of three charged Mac3 residues crucial for Mac3 binding of G-quadruplexes. Together (but not individually), Mac2 and Mac3 bind to and stabilize the ubiquitin E3 ligase RCHY1, which ubiquitinates the tumor-suppressor and antiviral p53 protein in SARS-CoV replicon experiments^232^.

Mac2 has also been shown to form a complex with Paip1 and PABP1, which enhances translation of viral but not host proteins^233^. A conservative mutation in the linker region connecting Mac3 and DPUP (ORF1a:S1494T) resulted in a highly attenuated SARS-CoV-2 mutant, and the defect was traced to an inability of the S1494T-mutant to enhance Paip1-PABP1 association^234^. The N-terminal region of Mac2 binds Paip1, and Paip1 binding of the S1494T mutant was unimpaired, implying that viral-protein translation enhancement depends on the joint cooperation of Mac2, Mac3, and perhaps DPUP.

In a study that aimed to identify all host and viral proteins that associate with DMVs during MHV infection, many of the identified host proteins were translation factors. Furthermore, siRNA knockdown of the translation factors that were identified resulted in a dramatic reduction in viral replication and viral titers despite having no effect on cell viability and only slightly reducing the translation of host proteins. Active translation takes place near DMVs, particularly early in infection, with activity peaking at 6 hpi; the same time period when NSP2 has its greatest effect on viral replication.

As NSP2, Mac2, and Mac3 are all implicated in enhancing viral protein translation and all localize to the perimeter of the DMVs during infection, it seems reasonable to suspect that the clear association between mutations in these regions of EPCI sequences may be the result of a functional interaction between them. The Mac2-Mac3-linker (Mac23L) region is the only of the three regions to have been identified as an immunodominant T-cell epitope^71^, and mutations in EPCI sequences are more common there than in the other two regions.

Mutations in the NSP2 region, for example, appear almost exclusively in sequences that also have mutations in the Mac23L region. The 29 EPCI sequences with one of the 2Macs NSP2 mutations (ORF1a:R232L, E233A, E233D, E233G, E233K) collectively have 33 Mac23L-region mutations, with 27 sequences have at least one (93.1%), strongly suggesting that these NSP2 mutations are deleterious in isolation but either compensate for loss-of-function immune-escape Mac23L mutations, or interact synergistically with Mac23L mutations.

### The 1Macs Mutation Pattern

The 1Macs (NSP**<u>1</u>**, NSP3 hypervariable region (HVR), **<u>Mac</u>**1-**<u>Mac</u>**2 Linker, **<u>Mac</u>**3) mutation pattern is notable for mutations converging on amino acid states found in RaTG13, one of the most closely related sarbecoviruses to SARS-CoV-2^235^. The mutations involved are ORF1a:H110Y and ORF1a:I114T in NSP1, ORF1a:P959S/L in NSP3-HVR, ORF1a:P1220L/S in the Mac1-Mac2 linker, and ORF1a:E1394D, T1395I, and A1397V in Mac3.

RATG13 residues at the first four of these sites are ORF1a:110Y, 114T, 959S, and 1220L. RaTG13 matches the SARS-CoV-2 Wuhan reference genome at all three Mac3 sites listed, though it does have M at ORF1a:1393 where SARS-CoV-2 has V. While several sarbecoviruses have 114T and 959S (including WIV1 and pangolin-CoV MP789), 110Y and 1220L are unique to RaTG13 (Fig SX).

The three NSP3 regions in 1Macs all occupy the DMV-pore prongs that project outward into the cytoplasm and therefore could plausibly interact with common host or viral factors. The HVR is essential for recruiting FXR1 to DMVs where it antagonizes stress granule formation early in infection, causes DMVs to cluster together through a mechanism involving liquid-liquid phase separation, and recruits host translation factors to DMVs for the efficient translation of viral RNA^236–237^. Notably, the two most important HVR amino acid residues for binding FXR1 are ORF1a:Y955 and F962, on either side of ORF1a:P959^237^. ORF1a:P1220 is in a disordered linker region connecting Mac1 and Mac2, and mutations in linker regions near Mac2 have been shown to affect the ability of Mac2 to enhance viral mRNA translation^234^. ORF1a:1394-1395 and 1397 are located 10-15 residues upstream of a G-quadruplex-binding region containing three conserved charged amino acids essential for SARS-CoV replicon replication^238^.

NSP1 is distributed widely throughout the cytoplasm during infection, where it binds near the mRNA entry site of ribosomes, shutting down translation of host proteins^250–253^. Unlike all other ORF1ab NSPs, NSP1 does not associate with DMVs during infection^227^. It is difficult, therefore, to imagine a direct physical interaction between ORF1a:110/114 and the other three 1Macs regions. However, at least two of the NSP3 regions, like NSP1, interact in some way with translation machinery of host cells.

### The Y1-6 Mutation Pattern

NSP6 contributes to DMV formation by forming zippered membrane channels that funnel ER material to nascent DMVs without allowing entry of large ER proteins^239^. NSP6 also associates with lipid droplets (LDs), which are then recruited to the surface of DMVs, where their lipid contents are used to fuel DMV growth^239–240^. Furthermore, NSP6 was recently shown to recruit ER-associated degradation (ERAD) vesicles called EDEMosomes to the DMV surface, where it provides membrane material for DMV expansion^240^. NSP6’s ability to associate with LDs and ER membranes has been traced to its C-terminal 80 amino acids (ORF1a:3780-3859), which are exposed to the cytoplasm and form two alpha helices shown to be essential for SARS-CoV-2 replication^239–241^. Mutations at three amino acid residues within this region (ORF1a:R3802, Y3803, L3808) correlate strongly with mutations in a highly conserved section of the Y1 domain of NSP3 (ORF1a:2467-2475) in EPCI sequences in what we call the Y1-6 mutation pattern (NSP3-**<u>Y1</u>**, NSP**<u>6</u>**).

The NSP3-Y1 domain is located on the cytoplasmic side of the DMV-pore structure, close to the outer DMV membrane, where it could potentially interact with cytoplasmic proteins, including NSP6^170, 172^–^-178^. While the Y1-6 regions are highly conserved in sarbecoviruses, among the 184 NSP3 and 305 NSP6 sequences we were able to align, we found a small number of sequences with variations in each. Specifically, nine sarbecovirus sequences have one mutation in the NSP6 region of the Y1-6 MP (ORF1a:3802-3808; eight with ORF1a:Y3803H and one with both L3806F and L3808F) and 15 sarbecovirus sequences have variations in the Y1 region of NSP3 (ORF1a:2467-2475; one with ORF1a:R2470H, three with L2475F, ten with L2475Q, and one with L2475V).

Remarkably, all nine sequences with Y1 mutations were also among those with mutations in the NSP6 region (figure 13), suggesting that these regions are also coevolving in other non-SARS-CoV-2 sarbecoviruses.

**Figure 13:**
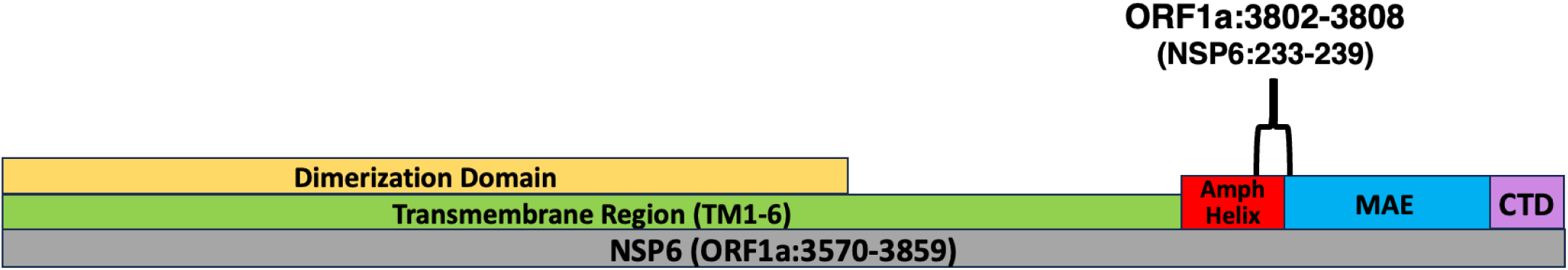
NSP6 with the Y1-6 MP-associated mutations labeled. Amph Helix = amphipathic helix; MAE = membrane-associated element; CDT = cytosolic C-terminal domain

**Figure 14:**
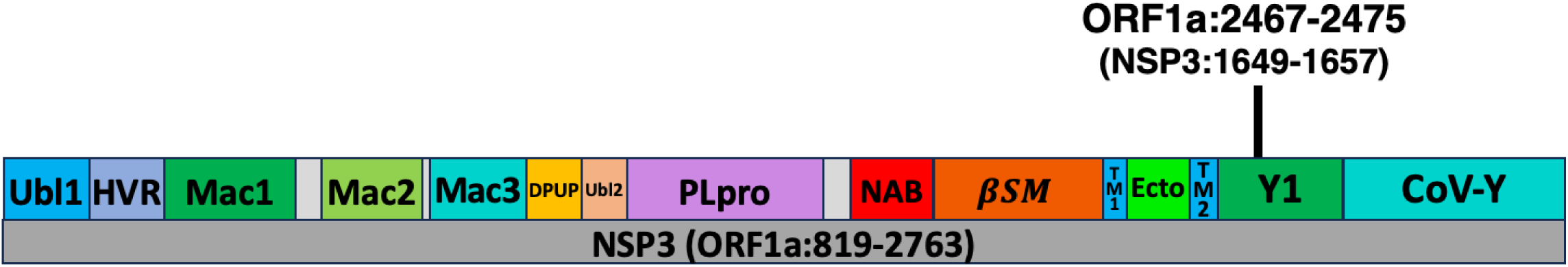
NSP3 with the Y1-6 MP-associated mutations labeled. Ubl1 = ubiquitin-like domain 1; HVR = hypervariable region (also called acidic domain); Mac1 = macrodomain 1; Mac2 = macrodomain 2; Mac3 = macrodomain 3; DPUP = Domain preceding Ubl2 and PLpro; Ubl2 = ubiquitin-like domain 2; PLpro = papain-like protease; NAB = nucleic acid-binding domain; βSM = betacoronavirus-specific marker domain; TM1 = transmembrane domain 1; Ecto = ectodomain; TM2 = transmembrane domain 2; Y1 = Nidovirus-conserved domain of unknown function; CoV-Y = Coronavirus-specific C-terminal domain

The underlying molecular basis of this coevolution is uncertain. Although it is unknown how NSP6 is recruited to DMVs from the cytoplasm, NSP3 is the only known protein on the DMV surface and it is therefore a top candidate for a direct interaction with NSP6 in order to facilitate lipid or membrane-material transfer to DMVs. It is therefore plausible that coevolution of amino acid residues ORF1a:2467-2475 of NSP3 with residues ORF1a:3802-3808 of NSP6 reflects optimization of direct interactions between these residues to ensure efficient DMV formation and growth.

### The NTD-DS Mutational Pattern

A common MP in EPCI sequences is NTD-DS (spike **<u>N</u>**-**t**erminal **<u>d</u>**omain **<u>d</u>**<u>i</u>**<u>s</u>**ulfide), involving the alteration or destruction of the C15-C136 spike NTD disulfide bond by disulfide-altering mutations (DAMs). The cysteine residue at position 15 can be removed through substitution (most commonly C15F), deletion (typically Δ14-16 or Δ14-22), or through alteration of the signal peptide cleavage site. The SARS-CoV-2 signal peptide is typically cleaved between residues 13 and 14^274^, but several substitutions shift the cleavage point so that C15 is cleaved from spike, including P9L, S12P, and S13I^274–276^. Furthermore, five EPCI sequences contain the combination of either P9S or P9T with S12F. Four of these sequences feature a substitution or deletion at C136, while the fifth lacks coverage at S:136-137, which is often indicative of a deletion or a mixed-mutation site. The P9S/T-S12F combination is therefore also considered to result in the loss of C15 in this analysis.

The five major surface-exposed loops of the SARS-CoV-2 spike NTD (N1-N5) compose an antigenic supersite, a major target of neutralizing antibodies^274^ and the location of the C15-C136 disulfide bond, which binds loops N1 and N3 together. The loss of either C15 or C136 (but not both) would result in an exposed, unpaired cysteine residue, which is generally destabilizing^301^. Loss of the remaining unpaired cysteine or the acquisition of a "replacement" cysteine therefore seems likely to confer a fitness increase in such a scenario.

One major 2021 variant, Epsilon (B.1.427/9), which circulated primarily in western USA, lost C15 (through S13I) and acquired a "replacement" cysteine with W152C, which formed a new disulfide bond with C136^275^. The new C136-C152 disulfide bond reshaped the NTD, resulting in total escape from all 10 anti-NTD neutralizing antibodies tested^275^.

Several sublineages of major variants, most notably Alpha (B.1.1.7) and BA.1, have lost the cysteine at S:15 without losing C136 or gaining an additional NTD cysteine, yet circulated at very low levels, indicating that C15 is dispensable, though its loss absent compensatory NTD mutations, is very likely deleterious. A search on Cov-spectrum for high-quality sequences that have C15-erasing mutations but lack any NTD mutations that likely compensate for C15 loss (W64C, ΔC136, C136X, G142C, Y144C, Y145C, S151C, W152C, Y248C, G257C, W258C, G261C) returns 4699 sequences (0.059%), indicating that while loss of C15 is not lethal, it is rare without compensatory mutations being present (Table S7a).

Similarly, a substantial proportion (205, 6.55%) of EPCI sequences possess mutations that result in the loss of C15 without the loss of C136 or the replacement of C15 with a new NTD cysteine. However, the number of EPCI sequences with mutations causing the loss of C15 or C136 that are accompanied by at least one compensatory mutation (123, 3.93%) is far higher than expected by chance alone given the overall frequency of such mutations in the dataset (p-value = 1.03x10^-104^, Table S8a). Furthermore, examination of these sequences reveals that mutations not directly involving cysteines strongly correlate with C15 loss and therefore likely compensate for the loss of the C15-C136 disulfide bond (Table 13).

**Table 13:** Mutations with the strongest association with high-confidence disulphide altering mutations (DAMs) in the expanded posited chronic-infection (EPCI) dataset. Spike RBM mutations are excluded. High-confidence DAMs include the following spike mutations: P9L, P9T, S12P, S13I, C15-, C15Y, C15S, C15F, C15G, C15L, C15R, W64C, F59C, S71C, C136-, D138-, C136R, C136H, C136F, C136Y, P139S, P139T, D142C, G142C, Y144C, Y145C, S151C, W152C, Y248C, W258C, G261C.

| Mutation | % of EPCI seqs with $\geq 1$ DAM that have the listed mutation as a private mutation | % of EPCI seqs with zero DAMs that have the listed mutation as a private mutation | Fold Increase<sup>3</sup> | Enrichment p-value<sup>4</sup> |
| --- | --- | --- | --- | --- |
| S:C136F | 7.91 | 0.04 | 197.72 | $1.08 \times 10^{-28}$ |
| S:P9L | 20.69 | 3.33 | 6.22 | $8.18 \times 10^{-28}$ |
| S:C136- | 6.62 | 0.08 | 82.78 | $1.20 \times 10^{-22}$ |
| S:D138- | 11.17 | 1.34 | 8.31 | $5.27 \times 10^{-20}$ |
| S:L242- | 18.42 | 4.92 | 3.74 | $1.24 \times 10^{-19}$ |
| S:A243- | 18.42 | 5.00 | 3.68 | $2.61 \times 10^{-19}$ |
| S:R403K | 11.40 | 2.84 | 4.01 | $4.59 \times 10^{-13}$ |
| S:T95I | 10.31 | 2.40 | 4.29 | $1.31 \times 10^{-12}$ |
| S:R21I | 3.07 | 0.04 | 76.69 | $9.49 \times 10^{-11}$ |
| S:D215G | 7.02 | 1.44 | 4.87 | $8.91 \times 10^{-10}$ |
| S:W152C | 2.42 | 0.00 | >120.78 | $2.10 \times 10^{-9}$ |
| S:R190S | 4.61 | 0.84 | 5.48 | $2.68 \times 10^{-7}$ |
| S:A688V | 3.95 | 0.64 | 6.16 | $7.18 \times 10^{-7}$ |
| S:N211- | 5.70 | 1.44 | 3.96 | $7.43 \times 10^{-7}$ |
| S:S151C | 1.54 | 0.00 | >76.69 | $4.02 \times 10^{-6}$ |
| S:L212I | 5.48 | 1.52 | 3.60 | $4.33 \times 10^{-6}$ |
| S:V642G | 5.04 | 1.32 | 3.82 | $5.54 \times 10^{-6}$ |
| S:C15- | 7.65 | 2.54 | 3.02 | $5.55 \times 10^{-6}$ |
| S:R190K | 3.07 | 0.44 | 6.97 | $6.24 \times 10^{-6}$ |
| M:D3Y | 3.51 | 0.64 | 5.48 | $8.83 \times 10^{-6}$ |
| S:G213E | 4.17 | 0.96 | 4.34 | $1.15 \times 10^{-5}$ |
| S:E554K | 2.63 | 0.32 | 8.22 | $1.23 \times 10^{-5}$ |
| S:W64C | 2.22 | 0.20 | 11.12 | $1.80 \times 10^{-5}$ |
| S:E406Q | 3.51 | 0.72 | 4.87 | $2.39 \times 10^{-5}$ |
| S:D339A | 1.32 | 0.00 | >65.74 | $2.63 \times 10^{-5}$ |
| M:A2V | 5.26 | 1.64 | 3.21 | $3.02 \times 10^{-5}$ |
| S:P217L | 2.19 | 0.24 | 9.13 | $4.66 \times 10^{-5}$ |
| S:P139S | 2.24 | 0.28 | 8.01 | $8.22 \times 10^{-5}$ |
| S:D215H | 2.63 | 0.48 | 5.48 | $1.46 \times 10^{-4}$ |
| S:H681R | 3.51 | 0.92 | 3.81 | $1.93 \times 10^{-4}$ |
| S:T791I | 4.17 | 1.32 | 3.15 | $2.80 \times 10^{-4}$ |
| N:Q244K | 1.54 | 0.12 | 12.78 | $3.14 \times 10^{-4}$ |
| S:P631S | 1.97 | 0.28 | 7.04 | $3.83 \times 10^{-4}$ |
| S:R158I | 1.10 | 0.04 | 27.39 | 9.01 x 10<sup>-4</sup> |
| S:W258C | 0.88 | 0.00 | >44.02 | 1.11 x 10<sup>-3</sup> |
| ORF1a:A1298V | 0.88 | 0.00 | >43.82 | 1.12 x 10<sup>-3</sup> |
| S:V615A | 0.88 | 0.00 | >43.82 | 1.12 x 10<sup>-3</sup> |
| S:H655Y | 2.85 | 0.80 | 3.56 | 1.40 x 10<sup>-3</sup> |
| S:N556K | 1.54 | 0.20 | 7.67 | 1.56 x 10<sup>-3</sup> |
| S:D574N | 2.63 | 0.76 | 3.46 | 2.70 x 10<sup>-3</sup> |
| S:R683Q | 1.97 | 0.48 | 4.11 | 4.77 x 10<sup>-3</sup> |
| S:W64G | 0.88 | 0.04 | 21.91 | 4.93 x 10<sup>-3</sup> |
| S:K854R | 1.32 | 0.20 | 6.57 | 6.06 x 10<sup>-3</sup> |
| S:S12F | 1.75 | 0.40 | 4.38 | 6.35 x 10<sup>-3</sup> |
| S:H1101R | 1.10 | 0.12 | 9.13 | 6.41 x 10<sup>-3</sup> |
| ORF1a:S2396I | 0.66 | 0.00 | >32.87 | 7.32 x 10<sup>-3</sup> |
| S:C136R | 0.66 | 0.00 | >32.87 | 7.32 x 10<sup>-3</sup> |
| S:P139T | 0.66 | 0.00 | >32.87 | 7.32 x 10<sup>-3</sup> |
| S:T791P | 0.66 | 0.00 | >32.87 | 7.32 x 10<sup>-3</sup> |
| S:D1084E | 0.66 | 0.00 | >32.87 | 7.32 x 10<sup>-3</sup> |

**Table 14:**
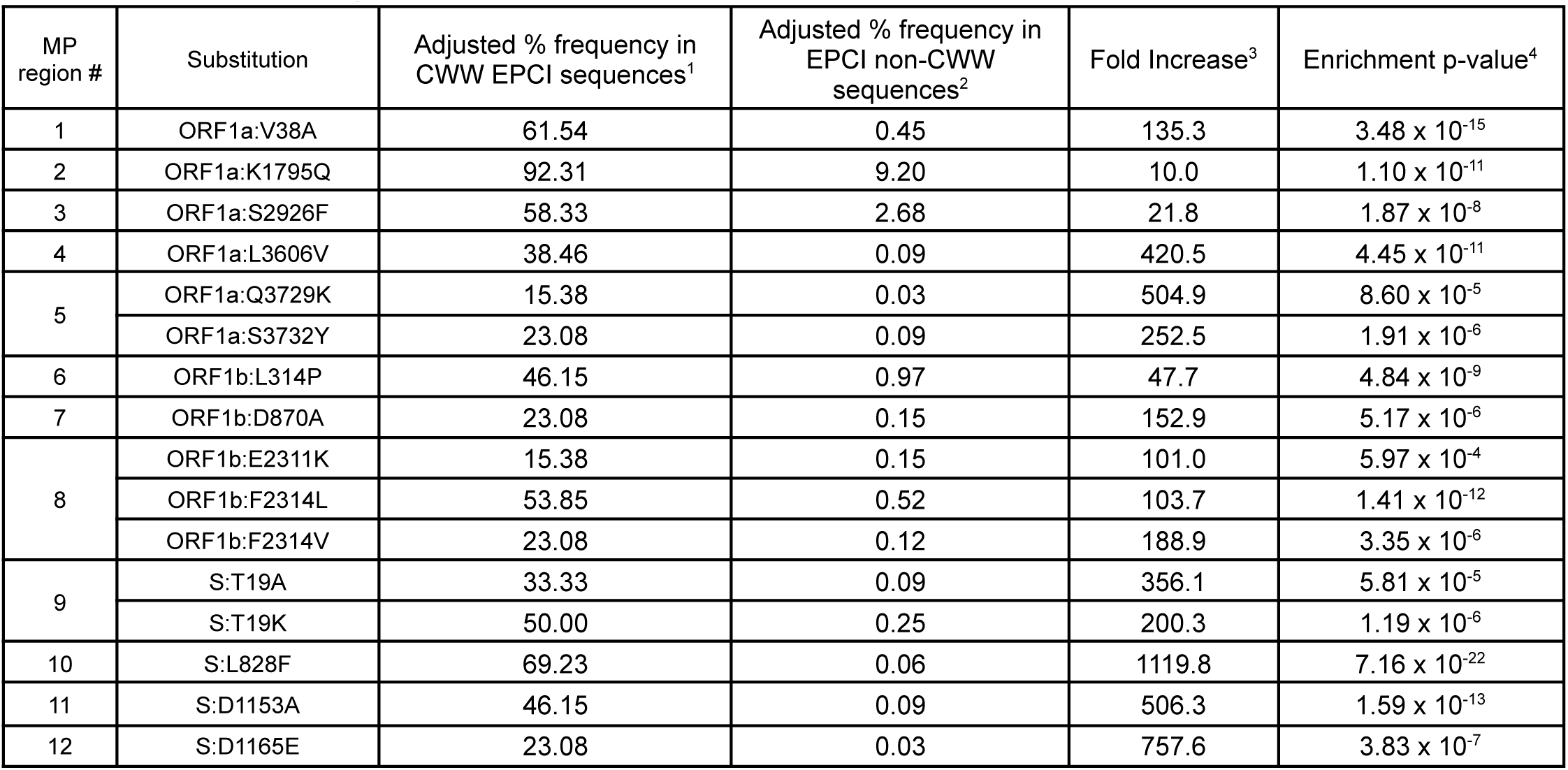
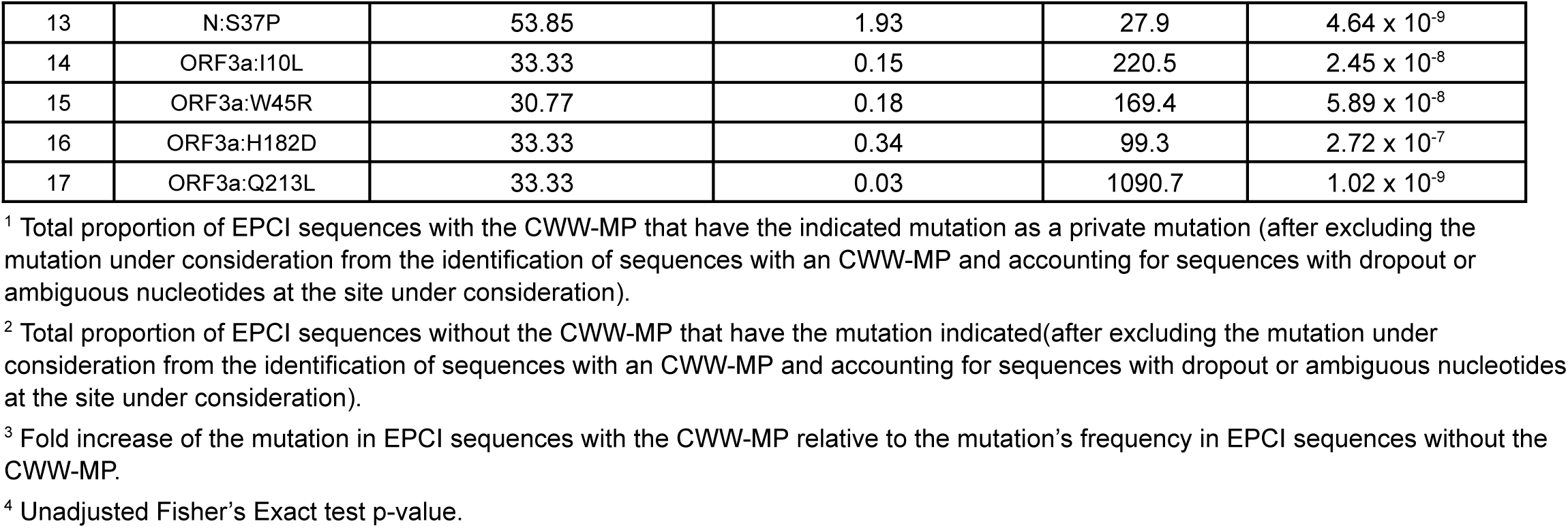
Full list of substitutions that occur significantly more frequently in EPCI sequences with a Cryptic Wastewater MP than they do in other EPCI sequences.

Foremost among these posited compensatory mutations are deletions that retain C136 but delete D138 and/or P139, usually along with at least Δ140-144. Presumably such deletions cause a restructuring of the regions around C136, reducing its exposure and destabilizing tendencies. S:P139S and P139T, both of which create an N-linked glycan motif (N-x-S/T) are also tightly associated with C15-erasing mutations. Of the 18 EPCI sequences with P139S/T (and no sequencing dropout in major disulfide-associated regions), 14 (77.8%) have known C15-erasing mutations (including P9L, C15F, C15R, ΔC15) or a new cysteine (W64C). Another has P9S and S13C, a combination which seems likely to alter signal peptide cleavage, while two of the remaining three possess the private deletions Δ142-143 which creates two consecutive acidic (D) residues at 144-145, and Δ243-244, which may together alter the environment around the disulfide region. The N137 glycan created by P139S/T may act to shield the unpaired C136 residue, dampening its reactivity and stabilizing the NTD.

Several other mutations in or adjacent to the N1, N3, or N5 NTD loops which do not involve the removal or addition of cysteine residues exhibit remarkably tight association with DAMs, including, R21I, T95I, and D215G. Thirteen of 14 EPCI sequences with R21I contain at least one other DAM, and those 13 contain a collective 18 DAMs. When measuring association of individual mutations with at least one DAM, P9L and C136F have the strongest associations (p<10^-26^ and p≤10^-19^, respectively; Fisher’s exact test), but with T95I having the third strongest (p<10^-12^) and D215G having the fourth strongest (p<10^-10^). The only other non-DAMs with p-values < 10^-5^ are ΔA243 (p<10^-8^), R190S (p<10^-7^), and R190K (p<10^-5^).

In contrast to the tolerance for standalone mutations removing C15, the loss of C136 without compensatory mutations appears likely to be lethal. An open Cov-spectrum search for high-quality sequences with a C136 mutation but no major known compensating mutations returns just 40 sequences (Table S7b). Nine of these sequences have acquired new cysteine residues at uncommon positions, including S71C, G75C, F79C, and R214C, while most of the rest display sequencing issues, such as NTD dropout, unrealistically large spike deletions (such as Δ85-149), or what are almost certainly artifactual reversions to the reference genome. EPCI sequences reveal a similar pattern. Of the 74 EPCI sequences with deletions or substitutions at C136, 70 have major known compensatory mutations. Of the remaining four, three have deletions of ≥13 amino acids in S:16-31. These extensive deletions seem very likely to either shift the signal peptide cleavage point beyond C15 or perhaps alter protein folding so that C15 is shunted into an internal, unexposed position in spike. The last of the C136-only EPCI sequences is from a lab whose sequences regularly feature extensive sequencing dropout and artifactual reversions to reference: both of which can be seen in numerous other sequences from the same upload. It therefore appears likely that C136 cannot be lost without either the loss of C15 or the acquisition of an additional cysteine with which C15 can create a new disulfide bond.

Elimination of the C15-C136 disulfide bond has taken on a new relevance with the global emergence of BA.3.2, which possesses P9L and Δ136-147, eliminating both C15 and C136. Notably, the BA.3.2 spike features numerous other mutations that are strongly associated with DAMs in EPCI sequences, including T95I, Δ243-244, R403K, V642G, and H681R (in BA.3.2.1), as well as E554D and A688D, which occur at the same sites as E554K and A688V, both among the top DAM-associated mutations.

An even more remarkable example of the NTD-DS MP is a chronic BA.2 infection suffered by a man living with HIV in Mexico, who had not received antiretroviral treatment in five years^287^. Initially hospitalized with severe Covid-19 in July 2023, the last sequence was collected from this patient on May 22, 2024, 315 days after initial hospitalization and likely more than two years after the initial infection. In addition to having two known DAMs in P9L and Δ136-152, this sequence also possessed an astonishing number of non-DAM mutations strongly associated with the NTD-DS MP, including T95I, G213E, D215G, Δ244, R346T, R403K, N417T, V615A, V642G, H681R, A688V, and M:D3Y. Setting aside the known DAMs, these include the #1, 2, 3, 4, 7, 10, 11, 12, 15, 28, 31, and 32 mutations most strongly associated with the NTD-DS MP (Table S9).

### Other Mutation Patterns

In addition to the MPs we have outlined in some detail, we detected numerous others in EPCI sequences. The Cryptic Wastewater MP (CWW) involves a number of extremely rare mutations that are common in very long-term chronic infections that have been detected in wastewater^242–245^. Remarkably, the CWW MP includes five reversions to consensus sarbecovirus amino acid residues, namely, ORF1a:V38A, ORF1a:K1795Q, ORF1a:L3606V, ORF1b:L314P (reversion), and ORF1b:F2314L, likely reflecting adaptation to the intestinal tract^242^, which is believed to be the primary site of viral replication in bats^246–247^. S:L828F and ORF3a:H182D are perhaps the two most distinctive CWW mutations, occurring almost exclusively in the longest-lasting and most highly mutated chronic-infection sequences.

Other notable MPs include the BsHel MP (NSP3-**<u>B</u>**etacoronavirus-**<u>S</u>**pecific Domainmutation, NSP13 **<u>Hel</u>**icase), which includes mutations from ORF1a:2165-2169 and ORF1b:1218-1220, and an unusual MP we call HR1M (Spike **<u>H</u>**eptad **<u>R</u>**epeat **<u>1</u>** + **<u>M</u>**embrane) that includes S:936 mutations (primarily S:D936H), M:H155N, ORF1a:3201 mutations, and deletions (often frameshifting) at ORF3a:255-256 (usually Δ26155-26158, which corresponds exactly to four unpaired nucleotides in a stem-loop in the genomic secondary RNA structure^248–249^). ORF3a forms giant membraned vesicles associated with early endosomes and the trans-Golgi network that are loaded with S and M proteins, possibly ferrying them to the ERGIC for virion assembly^289^. The formation of 3DBs has been traced to the C-terminal domain of ORF3a, which may explain the connection between S:D936 mutations, M:H155N, and ORF3a CTD deletions^289^. The full list of MPs detected by our algorithm can be found in Table S10.

## Conclusion

By analyzing EPCI sequences obtained by bronchoalveolar lavage (BAL), we present strong evidence for a distinct lower-respiratory tract (LRT) mutational signature in persistent infections. We also show that this same deep-lung MP is present in hundreds of other EPCI sequences, suggesting that prolonged lower-lung infection, probably primarily in the immunocompromised, is a widespread, and likely underrecognized, phenomenon that may contribute to some proportion of long Covid (PASC).

We also confirm the existence of a rare but distinct Cryptic wastewater MP in conventional URT-swab sequences, suggesting that the prolonged SARS-CoV-2 infections detected in wastewater occasionally migrate to the respiratory tract from the GI tract and may, in rare cases, be capable of transmitting from person to person^242, 254^. Notably, Gamma (P.1) possessed several mutations that have been rare in circulating SARS-CoV-2 variants but common in Cryptic wastewater sequences.

We also have detected frequent and extensive CD8 T cell escape mutations for common HLA types, especially A*01:01, indicating strong intrahost selective pressure to evade cellular immune responses. Given the clear overrepresentation of A*01:01 escape mutations, it would be interesting to investigate whether A*01:01 individuals are at higher risk of persistent SARS-CoV-2 infection than other HLA types. Such knowledge could help identify which individuals are more vulnerable to viral persistence and guide treatment strategies.

The function of other MPs detected in EPCI sequences is less clear, though we have outlined possible functional relationships between different regions in these MPs. Each persistent human infection represents, in some sense, an independent experiment in uninterrupted evolution. Researchers have serially passaged viruses in cell culture or in mice in an attempt to discover virulence factors and uncover hidden aspects of coronavirus genetics and the viruses life cycle^105, 255^. However, such passaging is prone to cell culture-specific adaptations such as deletion of the spike furin cleavage site, or the development of mutations rarely or never seen in sequences from humans.

The unprecedented amount of viral genetic data made available during the Covid pandemic has made it possible to decipher evolutionary patterns that occur in thousands of patients chronically infected by SARS-CoV-2.

Frequently, such sequences have foreshadowed mutations that months or years later became fixed in circulating variants. Furthermore, much about the SARS-CoV-2 life cycle remains unknown, and the mutation patterns that emerge during chronic infections are breadcrumb trails that may lead to the discovery of heretofore unknown interactions and functional connections between diverse SARS-CoV-2 proteins.

Lastly, millions of people continue to suffer the effects of long Covid (PASC), for which there are no approved treatments. While there are many hypotheses about the underlying causes of long Covid, they remain shrouded in doubt and uncertainty^56^. One major proposed mechanism of long Covid is the existence of persistent viral reservoirs in hard-to-sample bodily niches^50–55^. Some of the MPs we have described likely represent adaptations to specific tissues or organs, and unraveling the mechanisms and nature of such viral reservoirs may lay foundations for the development of effective long Covid treatments.

### Additional Information

The custom Julia code used in producing and analyzing the data used in this paper can be found at: https://github.com/ryhisner/MP_EPCI_code

## Supporting information

All supplementary information

