## Supplementary material for "Coevolving Mutations in Chronic SARS-CoV-2 Infections": All supplementary information

### Supplementary Data

**Supplementary Table 1**: Excluded mutations and justifications

| Mutation | Reason |
|---|---|
| ORF1a:M2606I | Inherited B.1.2 mutation |
| N:D377Y | Inherited B.1.2 mutation |
| ORF3a:A72S | Inherited B.1.258 mutation |
| ORF1a:E3764D | Inherited B.1.258 mutation |
| ORF1a:I2283T | Artifactual reversion in XEC miscategorized as KP.3 or KP.3.3 by Nextclade |
| ORF1a:A599T | Artifactual "private" mutation in XEC miscategorized as KP.3 or KP.3.3 by Nextclade |
| S:F59S | Artifactual "private" mutation in XEC miscategorized as KP.3 or KP.3.3 by Nextclade |
| ORF8:L60F | Artifactual "private" mutation in AY.44. Over 125000 seqs have both ORF8:L60F & ORF1a:H2125Y |
| ORF1a:H2125Y | Artifactual "private" mutation in AY.44. Over 125000 seqs have both ORF8:L60F & ORF1a:H2125Y |
| S:Q146K | Extremely homoplasic XBB mutation and/or artifact. Impossible to tell if inherited or private. |
| N:M203K | Recombinant Delta/Omicron "mutation" |
| N:G204R | Recombinant Delta/Omicron "mutation" |
| N:R204G | Recombinant Delta/Omicron reversion "mutation" |
| ORF1b:M1156I | BQ.1 mutation misattributed as private in 3 sequences |
| ORF:L61D | Artifactual 3-nuc reversion |
| ORF1a:S135R | Omicron mut misattributed as private in 3 recombinants |
| M:A30T | Very common artifactual reversion in BA.2.86* lineages |
| ORF1a:S1612L | Inherited Beta mutation miscategorized by Nextclade as private |
| ORF7b:E39* | Inherited Beta mutation miscategorized by Nextclade as private |
| S:F18L | All five on list in Beta, likely artifactual |
| ORF1a:A2554V | Inherited AY.44 mutation miscategorized by Nextclade as private |
| ORF1b:H1087Y | Inherited AY.44 mutation miscategorized by Nextclade as private |
| ORF1a:M2796T | Inherited BA.1.17 mutation miscategorized by Nextclade as private |
| ORF1a:P1803S | Inherited BA.1.17 mutation miscategorized by Nextclade as private |
| ORF3a:P104S | Inherited AY.103 mutation miscategorized by Nextclade as private |
| N:A208S | Inherited AY.103 mutation miscategorized by Nextclade as private |
| ORF8:K68* | Inherited B.1.1.7 mutation miscategorized by Nextclade as private |
| ORF1b:P1975S | Inherited B.1.1.7 mutation miscategorized by Nextclade as private |
| ORF1b:V2178F | Inherited BA.1.1 mutation miscategorized by Nextclade as private |
| ORF1b:I97V | Inherited BA.1.1 mutation miscategorized by Nextclade as private |
| ORF1b:A2589T | Inherited BA.5.1 mutation miscategorized by Nextclade as private |
| ORF9b:T83I | Inherited BA.5.1 mutation miscategorized by Nextclade as private |
| ORF1a:T842I | From 5' End Recombination with BA.2 |
| ORF1a:S135R | From 5' End Recombination with BA.2 |
| ORF3a:V48F | Inherited BE.1 mutation miscategorized by Nextclade as private |
| ORF3a:G49C | Inherited BE.1 mutation miscategorized by Nextclade as private |
| ORF1a:A1204T | Inherited BE.1 mutation miscategorized by Nextclade as private |

| | |
|---|---|
| S:A376P | Butchered mutation interpretation by Nextclade for ∆374-375 or ∆374-376 |
| S:T376P | Butchered mutation interpretation by Nextclade for ∆374-375 or ∆374-376 |
| S:F375A | Butchered mutation interpretation by Nextclade for ∆374-375 or ∆374-376 |
| S:S375A | Butchered mutation interpretation by Nextclade for ∆374-375 or ∆374-376 |
| S:D339G | An artifactual reversion >90% of the time, likely 100% |
| S:N417K | An artifactual reversion >90% of the time, likely 100% |
| S:K440N | An artifactual reversion >90% of the time, likely 100% |
| ORF1b:T1555I | Inherited B.1.2 mutation miscategorized by Nextclade as private |
| ORF1a:A4285V | Inherited B.1.2 mutation miscategorized by Nextclade as private |
| ORF1a:N2361K | Inherited B.1.2 mutation miscategorized by Nextclade as private |
| ORF1a:V3782I | Inherited mutation in Canadian BA.1 branch miscategorized by Nextclade as private |
| S:K1191N | Inherited B.1.1.7 mutation miscategorized by Nextclade as private |
| S:R158G | Artifactual Delta reversion |
| ORF1a:T3646A | Inherited Delta mut, miscategorized in recombinants |
| ORF1b:A1918V | Inherited Delta mut, miscategorized in recombinants |
| ORF7b:I2V | Corresponds to ORF7a:*122W |
| ORF1b:P218L | Inherited B.1.1.7 mutation miscategorized by Nextclade as private |
| ORF1a:G2118D | B.1 Inherited mutation miscategorized by Nextclade as private (co-occurs with S:P681H) |
| ORF1a:A1298V | Inherited BA.1 mut, miscategorized by Nextclade as private |
| ORF3a:V112F | Inherited B.1.1 mutation miscategorized by Nextclade as private |
| ORF1b:N2695I | Inherited B.1.1 mutation miscategorized by Nextclade as private |
| ORF3a:I35K | Inherited XBB.1.16.11 mutation miscategorized by Nextclade as private |
| ORF1b:E1871G | Inherited B.1.1.7 mutation miscategorized by Nextclade as private |
| ORF1b:D1903Y | Inherited XBB.1.16.31 mutation miscategorized by Nextclade as private |
| ORF1a:K2219R | Inherited XBB.1.16.31 mutation miscategorized by Nextclade as private |
| ORF1b:K1094R | Inherited B.1.2 mutation miscategorized by Nextclade as private |
| ORF1a:S771I | Inherited XBB.1.5 mutation miscategorized by Nextclade as private |
| N:S26C | Inherited BA.5 mutation miscategorized by Nextclade as private |
| ORF1a:V108I | Inherited BA.5.1 mutation in Scandinavian branch miscategorized by Nextclade as private |
| ORF1b:G128S | Inherited BA.5.1 mutation in Scandinavian branch miscategorized by Nextclade as private |
| ORF1b:N2452S | Inherited BA.5.1 mutation in Scandinavian branch miscategorized by Nextclade as private |
| ORF1b:S1182T | Inherited FL.4 mutation in Australian branch miscategorized by Nextclade as private |
| ORF1b:P1570T | Inherited BA.1.1.1 mutation in Belgian branch miscategorized by Nextclade as private |
| ORF1b:E2355Q | Inherited BA.1.1.1 mutation in Belgian branch miscategorized by Nextclade as private |
| S:T19R | Delta mutation present in recombinants, falsely labeled as private mutation |
| ORF7a:A66V | Inherited B.1.429 mutation miscategorized by Nextclade as private |
| ORF1a:T3459M | Inherited B.1.429 mutation miscategorized by Nextclade as private |
| ORF7b:M1T | Corresponds to ORF7a:*122R |
| ORF1b:A869V | Inherited KF.1 (FL.15.1.1.1) mutation miscategorized by Nextclade as private |
| ORF1a:E1015G | Inherited KF.1 (FL.15.1.1.1) mutation miscategorized by Nextclade as private |
| ORF7b:L4F | Inherited BA.1 mutation on Swedish branch miscategorized by Nextclade as private |
| ORF1a:V365I | Inherited B.1.2 mutation miscategorized by Nextclade as private |
| ORF1b:K1094R | Inherited B.1.2 mutation miscategorized by Nextclade as private |
| ORF1b:D1869Y | Inherited C.37 and AY.85 mutation miscategorized by Nextclade as private |
| ORF3a:L65H | Inherited BN.1.3.1 mutation (Italian branch) miscategorized by Nextclade as private |

| | |
|---|---|
| ORF3a:S220I | Inherited B.1.466 (Indonesia) mutation miscategorized by Nextclade as private |
| ORF1b:A176T | Inherited EG.5.1 mutation miscategorized by Nextclade as private |
| ORF1a:E352D | Inherited B.1.1.176 (Canada) mutation miscategorized by Nextclade as private |
| ORF1b:A434S | Inherited B.1.1.176 (Canada) mutation miscategorized by Nextclade as private |
| ORF1a:G3676C | Inherited B.1.1.176 (Canada) mutation miscategorized by Nextclade as private |
| ORF1a:Q991L | Inherited BA.5.2 (North America) mutation miscategorized by Nextclade as private |

**Supplementary Table S2a**: Detailed results for running the MPD function on 100 APCI substitution datasets.
https://github.com/ryhisner/MP_Substitutions__Main_Results/blob/main/APCI_Substitutions__Results__EPCI_8_10_95__HQCS_5_1_5__2026_05_05__100_Runs.xlsx

**Supplementary Table S2b:** Detailed results for running the MPD function on 100 APCI position-only datasets.
https://github.com/ryhisner/MP_Positions__Main_Results/blob/main/APCI_Positions__Results__EPCI_8_10_95__HQCS_5_1_5__2026_05_05__100_Runs.xlsx

**Supplementary Table S3**: Bloom Lab deep mutational scanning measurements for ACE2 binding and sera escape for major BAL mutations.
https://github.com/ryhisner/MP_BAL_DMS_ACE2

**Supplementary Table S4**: Pairwise associations between mutations in the EPCI dataset, ranked by Fisher's Exact Test p-value for enrichment.
https://github.com/ryhisner/MP_EPCI_pairwise_correlations/blob/main/Top_EPCI_Pairwise_Mut_Correlations__2026_03_15.xlsx

**Supplementary Table 5a**: NSP15, ORF6, and ORF9b loss-of-function (LoF) mutations and BAL-MP statistics

| | |
|---|---|
| NSP15 LoF Muts | ORF1b:S2339A, S2339F, S2339Y, S2339P, K2340N, K2340E, K2340T |
| ORF6 Lof Muts | ORF6:M1T, M1I, M1V, Q8*, W27*, Y49*, Q56*, M58K, E59* |
| ORF9b LoF Muts | ORF9b:M1K, M1T, M1I, M1L, M1V, E7*, Q18*, Q20*, Q34*, E65*, Q77* |

**Supplementary Table 5b : NSP15, ORF6, and ORF9b loss-of-function (LoF) mutation BAL-MP statistics**

| Protein | Avg BAL-MP Muts/LoF seq | Ratio of Avg BAL-MP Muts/LoF seq to Avg BAL-MP Muts/seq for all other EPCI seqs | Collective Percentile Rank of LoF muts in terms of Avg # of BAL muts/seq among all EPCI muts that occur ≥10 times |
|---|---|---|---|
| NSP15 | 2.61 | 2.72 | 94.16% |
| ORF6 | 2.40 | 2.42 | 92.41% |
| ORF9b | 2.98 | 3.07 | 96.55% |

**Supplementary Table 6**: ORF7a C-terminal Extension Mutations and EPCI counts (All mutation counts in this table exclude the ORF7a-extension mutation itself, even if it qualifies as a BAL-MP mutation. For example, if a sequence has ORF7a:*122R plus three other BAL-MP muts, it counts as having three BAL-MP muts, not four.)

| Mutation | Count | Avg BAL-MP Muts/seq | Ratio of Avg BAL-MP Muts/7a-extension seq to Avg BAL-MP Muts/seq for all other EPCI seqs | BAL-MP mut count Percentile Rank for mut among all EPCI muts that occur ≥ "Count" times |
|---|---|---|---|---|
| ORF7a:*122- | 12 | 3.17 | 3.20 | 97.59% |

| | | | | |
|---|---|---|---|---|
| ORF7a:*122R | 4 | 5.00 | 5.00 | 99.90% |
| ORF7a:*122W | 3 | 2.33 | 2.33 | 88.67% |
| ORF7a:*122S | 1 | 3.00 | 3.00 | 96.82% |
| ALL | 20 | 3.40 | 3.43 | 99.59% |

**Supplementary Table S7a**: Open Cov-Spectrum search for sequences with one disulfide-altering mutation (DAM) but no known compensatory mutations. Search performed on March 13, 2026.

| |
|---|
| **Search Query**: !Nextcladepangolineage:BA.3.2* &<br>[exactly-1-of: S:P9L, S:P9T, S:S12P, S:S13I, S:C15, S:W64C, S:C136, S:142C, S:Y144C, S:Y145C, S:S151C, S:W152C, S:Y248C, S:W258C] &<br>[16-of: S:P9P, S:S12S, S:S13S, S:C15C, (S:R21R \| S:R21T), S:W64W, S:C136C, (S:D138D \| S:D138Y \| S:D138H), S:P139P, (S:G142G \| S:G142D \| S:G142-), (S:Y144Y \| S:Y144- \| S:Y144S \| S:Y144N \| S:Y144H), (S:Y145Y \| S:Y145D \| S:Y145H \| S:Y145-), S:S151S, (S:W152W \| S:W152R), (S:Y248Y \| S:Y248N \| S:Y248S \| S:Y248H), (S:G257G \| S:G257D \| S:G257S), S:W258W]<br>**Advanced QC Filters**: Overall score ≤ 29, 0.9 ≤ Coverage (from 0 to 1) |
| **Number of sequences returned**: 4699 (of 7915523 sequences) (0.059%) |
| **Top Pango lineages Returned**: BA.1* (1537), B.1.1.7* [Alpha] (987), B.1.617.2* [Delta] (645) |

**Supplementary Table S7b**: Open Cov-Spectrum search results for sequences with a substitution or deletion at S:C136 and no known compensatory mutations. Search performed on May 23, 2026.

| |
|---|
| **Search Query**: S:C136 &<br>[14-of: S:P9P, S:S12S, S:S13S, S:C15C, S:V16V, S:N17N, (S:R21R \| S:R21T), S:W64W, S:S151S, (S:W152W \| S:W152R), (S:Y248Y \| S:Y248N \| S:Y248S \| S:Y248H), (S:G257G \| S:G257D \| S:G257S), S:W258W, S:G261G]<br>**Advanced QC Filters**: Overall score ≤ 29, 0.9 ≤ Coverage (from 0 to 1) |
| **Number of sequences returned (all times)**: 40 (of 7920222 sequences) (0.00051%) |
| **Number of sequences returned with collection date on or after Jan. 1, 2022**: 6 (of 3884114 sequences) (0.00015%) |
| **Sequences with collection dates after 2022-11-01:** 0 (of 867142 sequences) |
| **Notable features among 40 returned sequences, all times**: S71C (1), G75C (6), F79C (1), R214C (1), dropout artifact (1), unrealistically large deletions (9), clearly false reversions to reference (3), extensive NTD dropout (2) |
| **Notable features among six Omicron-era sequences collected on or after Jan 1, 2022**: S:S71C (1), S:F79C (1), dropout artifact (1) |

**Supplementary Table 8a**: Statistics for enrichment of -Loss Compensatory Mutations in EPCI Sequences with ≥1 C15/C136 Loss Mutations.

<u>C15/C136-Loss Mutations</u>: P9-, P9L, S12P, S13I, C15F, C15R, C15L, C15G, C15Y, C15S, C15-, C136-, C136F, C136H, C136R, C136Y, and the combination of P9S/P9T and S12F.

<u>C15/C136-Loss Compensatory Mutations include all C15/C136-Loss Mutations plus</u>: R21I, F59C, W64C, S71C, F79C, P139S, P139T, G142C, D142C, Y144C, Y145C, S151C, W152C, S247C, Y248C, G257C, W258C, G261C

| Total EPCI Seqs with Coverage at all NTD-DS Mut Sites | Total EPCI Seqs with C15/C136-Loss Muts | Pct of EPCI Seqs with ≥1 C15/C136-Loss Muts | Total EPCI Seqs with ≥1 C15/C136-Loss Muts with ≥1 C15/C136-Loss Compensatory Muts | Total EPCI Seqs with ≥1 C15/C136-Loss Muts and Zero C15/C136-Loss Compensatory Muts | -log10 Fisher Exact Test p-value for Enrichment of C15/C136-Loss Compensatory Muts in EPCI Seqs with ≥1 C15/C136-Loss Muts | Fisher Exact Test p-value for Enrichment of C15/C136-Loss Compensatory Muts in EPCI Seqs with ≥1 C15/C136-Loss Muts | Pct of EPCI Seqs with ≥1 C15/C136-Loss Mut that Possess ≥1 C15/C136-loss Compensatory Muts | Pct of EPCI Seqs with Zero C15/C136-Loss Muts that Possess ≥1 C15/C136-loss Compensatory Muts | C15/C136-Compensatory Mut Fold-Increase in C15/C136-Loss EPCI Seqs Relative to non-C15/C136-Loss EPCI Seqs |
|---|---|---|---|---|---|---|---|---|---|

| | | | | | | | | | |
|---|---|---|---|---|---|---|---|---|---|
| 3130 | 328 | 11.19% | 123 | 205 | $1.03 \times 10^{-104}$ | 103.99 | 37.50% | 0.86% | 43.78 |

**Supplementary Table 8b**: Statistics for enrichment of C15-Loss Compensatory Mutations in EPCI Sequences with ≥1 C15-Loss Mutations.

<u>C15-Loss Mutations</u>: P9-, P9L, S12P, S13I, C15F, C15R, C15L, C15G, C15Y, C15S, C15-, and the combination of P9S/P9T + S12F.

<u>C15-Loss Compensatory Mutations</u>: R21I, W64C, F59C, S71C, F79C, C136-, C136F, C136H, C136R, C136Y, P139S, P139T, G142C, D142C, Y144C, Y145C, S151C, W152C

| Total EPCI C15 with Coverage at all C15-Loss and C15-Loss Compensatory Mut Sites | Total EPCI Seqs with C15-Loss Muts | Pct of EPCI Seqs with ≥1 C15-Loss Muts | Total EPCI Seqs with ≥1 C15-Loss Muts and ≥1 C15-Loss Compensatory Muts | Total EPCI Seqs with ≥1 C15-Loss Muts with Zero C15-Loss Compensatory Muts | -log10 Fisher Exact Test p-value for Enrichment of C15-Loss Compensatory Muts in EPCI Seqs with ≥1 C15-Loss Muts | Fisher Exact Test p-value for Enrichment of C15-Loss Compensatory Muts in EPCI Seqs with ≥1 C15-Loss Muts | Pct of EPCI Seqs with ≥1 C15-Loss Mut that Possess ≥1 C15-loss Compensatory Muts | Pct of EPCI Seqs with Zero C15-Loss Muts that Possess ≥1 C15-loss Compensatory Muts | C15-Compensatory Mut Fold-Increase in C15-Loss EPCI Seqs Relative to non-C15-Loss EPCI Seqs |
|---|---|---|---|---|---|---|---|---|---|
| 3214 | 328 | 9.89% | 112 | 206 | $2.52 \times 10^{-95}$ | 94.60 | 35.22% | 0.86% | 40.80 |

**Supplementary Table 8c**: Statistics for enrichment of C15-Loss Compensatory Mutations in EPCI Sequences with ≥1 C15-Loss Mutations.

<u>C15-Loss Mutations</u>: P9-, P9L, S12P, S13I, C15F, C15R, C15L, C15G, C15Y, C15S, C15-, and the combination of P9S/P9T + S12F.

<u>C15-Loss Compensatory Mutations</u>: R21I, W64C, F59C, S71C, F79C, C136-, C136F, C136H, C136R, C136Y, P139S, P139T, G142C, D142C, Y144C, Y145C, S151C, W152C

| Total EPCI C136 with Coverage at all C136-Loss and C136-Loss Compensator y Mut Sites | Total EPCI Seqs with C136-Loss Muts | Pct of EPCI Seqs with ≥1 C136-Loss Muts | Total EPCI Seqs with ≥1 C136-Loss Muts and ≥1 C136-Loss Compensatory Muts | Total EPCI Seqs with ≥1 C136-Loss Muts with Zero C136-Loss Compensatory Muts | -log10 Fisher Exact Test p-value for Enrichment of C136-Compensatory-Muts in EPCI Seqs with ≥1 C136-Loss Muts | Fisher Exact Test p-value for Enrichment of C136-Compensatory-Muts in EPCI Seqs with ≥1 C136-Loss Muts | Pct of EPCI Seqs with ≥1 C136-Loss Mutation that Possess ≥1 C136-loss Compensatory Muts | Pct of EPCI Seqs with Zero C136-Loss Muts that Possess ≥1 C136-loss Compensatory Muts | C136-Compensatory Mut Fold-Increase in C136-Loss EPCI Seqs Relative to non-C136-Loss EPCI Seqs |
|---|---|---|---|---|---|---|---|---|---|
| 3218 | 74 | 2.30% | 70 | 4 | $1.98 \times 10^{-68}$ | 67.70 | 94.59% | 7.79% | 12.14 |

**Supplementary Table 9**: Non-disulfide-altering mutations (non-DAMs) associated with EPCI sequences with one or more DAMs, listed by lowest Fisher's Exact Test p-value.

| Mutation | Match_Ct (EPCI seqs w/ ≥1 DAM + matchmut) | Total EPCI seqs w/ Mutation | % of seqs w/Mutation with ≥1 DAM | % of all non-DAM EPCI seqs with Mutation | Fold Increase of Mutation in EPCI seqs with ≥1 DAM relative to all other EPCI seqs | p-value, Fisher's Exact Test |
|---|---|---|---|---|---|---|
| S:A243- | 85 | 213 | 39.91 | 4.41 | 9.06 | $2.04 \times 10^{-19}$ |
| S:R403K | 54 | 124 | 43.55 | 2.34 | 18.63 | $2.65 \times 10^{-14}$ |
| S:T95I | 47 | 110 | 42.73 | 2.09 | 20.40 | $3.73 \times 10^{-12}$ |
| S:D215G | 35 | 70 | 50.00 | 1.15 | 43.54 | $1.20 \times 10^{-11}$ |
| S:R21I | 14 | 15 | 93.33 | 0.03 | 2895.20 | $8.23 \times 10^{-11}$ |
| S:R190S | 21 | 42 | 50.00 | 0.68 | 73.24 | $2.40 \times 10^{-7}$ |
| S:A688V | 19 | 36 | 52.78 | 0.55 | 95.68 | $3.23 \times 10^{-7}$ |
| S:R190K | 15 | 26 | 57.69 | 0.36 | 162.17 | $1.49 \times 10^{-6}$ |
| S:N211- | 26 | 68 | 38.24 | 1.38 | 27.77 | $5.51 \times 10^{-6}$ |
| S:V642G | 23 | 57 | 40.35 | 1.11 | 36.33 | $7.15 \times 10^{-6}$ |
| S:G213E | 19 | 44 | 43.18 | 0.81 | 53.10 | $1.58 \times 10^{-5}$ |
| S:L212I | 25 | 68 | 36.76 | 1.41 | 26.08 | $1.96 \times 10^{-5}$ |
| M:D3Y | 16 | 34 | 47.06 | 0.58 | 80.63 | $2.20 \times 10^{-5}$ |

| | | | | | | |
|---|---|---|---|---|---|---|
| S:E406Q | 16 | 35 | 45.71 | 0.62 | 74.18 | $3.49 \times 10^{-5}$ |
| S:D215H | 13 | 25 | 52.00 | 0.39 | 134.03 | $3.90 \times 10^{-5}$ |
| S:H681R | 17 | 39 | 43.59 | 0.71 | 61.01 | $4.13 \times 10^{-5}$ |
| S:P217L | 10 | 16 | 62.50 | 0.19 | 323.13 | $4.39 \times 10^{-5}$ |
| M:A2V | 24 | 69 | 34.78 | 1.48 | 23.57 | $8.60 \times 10^{-5}$ |
| S:E554K | 11 | 20 | 55.00 | 0.29 | 189.32 | $8.86 \times 10^{-5}$ |
| S:P631S | 10 | 17 | 58.82 | 0.23 | 260.59 | $9.22 \times 10^{-5}$ |
| S:P621S | 25 | 75 | 33.33 | 1.64 | 20.29 | $1.35 \times 10^{-4}$ |
| S:T299I | 30 | 98 | 30.61 | 2.25 | 13.60 | $1.58 \times 10^{-4}$ |
| S:D339A | 5 | 5 | 100.00 | 0.00 | >3113.0 | $1.66 \times 10^{-4}$ |
| S:T791I | 20 | 55 | 36.36 | 1.14 | 31.82 | $1.85 \times 10^{-4}$ |
| S:I212- | 21 | 62 | 33.87 | 1.34 | 25.25 | $4.00 \times 10^{-4}$ |
| S:K529N | 8 | 14 | 57.14 | 0.19 | 295.62 | $7.37 \times 10^{-4}$ |
| S:L176F | 18 | 52 | 34.62 | 1.11 | 31.21 | $8.40 \times 10^{-4}$ |
| S:R158I | 5 | 6 | 83.33 | 0.03 | 2593.33 | $8.63 \times 10^{-4}$ |

**Supplementary Table S10**: Full mutation-pattern detection results for EPCI and APCI datasets.
https://github.com/ryhisner/MP_EPCI_and_APCI_MPD_Results

**Supplementary Table S11**: Sensitivity Test Results for EPCI and APCI datasets utilizing stricter inclusion requirements for EPCI and APCI sequences.
https://github.com/ryhisner/MP_Substitutions__Sensitivity_Test_Results_v1
https://github.com/ryhisner/MP_Substitutions__Sensitivity_Test_Results_v2
https://github.com/ryhisner/MP_Positions__Sensitivity_Test_Results_v1
https://github.com/ryhisner/MP_Positions__Sensitivity_Test_Results_v2

**Supplementary Table S12**: The full, detailed BAL-MP results and the custom Julia code used in the analysis.
https://github.com/ryhisner/BAL_MP_Results

**Supplementary Table S13**: The complete, detailed NTD-Disulfide results and the code used for the analysis.
https://github.com/ryhisner/NTD_Disulfide_Results

**Supplementary Table S14**: The complete, detailed MDP-function results for the EPCI and APCI datasets and the distribution of Nextstrain clades, Pango lineages, and “Megaclades” for each mutation pattern can be found at the following Github repositories.
https://github.com/ryhisner/MP_Substitutions__Main_Results
https://github.com/ryhisner/MP_Positions__Main_Results
